# A critical role of stable grain filling rate in maximizing rice yield revealed by whole plant carbon nitrogen interaction modeling

**DOI:** 10.1101/2020.10.06.329029

**Authors:** Tian-Gen Chang, Zhong-Wei Wei, Zai Shi, Yi Xiao, Honglong Zhao, Shuo-Qi Chang, Mingnan Qu, Qingfeng Song, Faming Chen, Fenfen Miao, Xin-Guang Zhu

## Abstract

Crop yield is co-determined by potential size of the harvest organ, photosynthetic potential of source organs, and pattern of partitioning and use of photosynthates among sink organs. Given a sufficient potential size of the harvest organ at flowering, how to fully fill them remains a central challenge in crop breeding for high yields. Here, we develop a kinetic model of rice grain filling, scaling primary biochemical and biophysical processes to whole-plant carbon and nitrogen dynamics. Predicted post-anthesis physiological and agronomic behaviors validate experimental observations under six endogenous and external perturbations. By large scale *in silico* screening, we show here that a stable grain filling rate from flowering to harvest is required to maximize grain yield, which is validated here in two independent super-high yielding rice cultivars (~21 t ha^-1^ rough rice yield at 14% moisture). On the other hand, we show grain yields in an elite rice cultivar may increase by about 30-40% by stabilizing its grain filling rate. Intriguingly, we have found that the sum of grain filling rates around 15 and 38 days after flowering largely determines grain yield, and have further developed a novel *in situ* approach quantifying grain filling rates and grain yield precisely with the measurements of ear respiratory rates (r>0.93). Potential post-anthesis molecular targets to maximize rice yield include delaying leaf senescence, enhancing leaf sucrose synthesis and export, limiting root growth, strengthening stem starch synthesis, accelerating endosperm starch synthesis, and moderating endosperm cell division. Our study provides an effective computational framework for post-anthesis crop physiology research and ideotype design.

## INTRODUCTION

To meet the increasing global food demand, there is a tremendous need to greatly increase crop yields (Long, Marshall-Colon, & Zhu, 2015; Ray, Mueller, West, & Foley, 2013). With the identification of a number of superior alleles controlling grain yield/quality related traits, and the great advances in genome editing technology, molecular breeding by rational design is becoming a new important route for accelerated crop improvement (Peleman & van der Voort, 2003; Qian, Guo, Smith, & Li, 2016; Van Der Straeten et al., 2020; H. Yu et al., 2021). However, a dilemma presented is that in crop breeding practice, incorporation of superior alleles into a germplasm does not always lead to higher yield. This is because these superior alleles may not improve the most limiting factor(s) of the given germplasm, or even create new limiting factors (Garcia-Molina & Leister, 2020; Uga, 2021). Thus, a key remaining issue is a mechanistic understanding and quantification of the idealized morphological and physiological traits of a plant variety, known as crop ideotype (Donald, 1968), as an objective during rational crop improvement (Bailey-Serres, Parker, Ainsworth, Oldroyd, & Schroeder, 2019; Tian-Gen Chang, Chang, Song, Perveen, & Zhu, 2019; Paul, Watson, & Griffiths, 2019; Rötter, Tao, Höhn, & Palosuo, 2015; Uga, 2021).

Crop yield is co-determined by 1) canopy photosynthesis, which reflects the “source capacity”, 2) the potential size of all fertile spikelets, which reflects the “sink capacity”, and 3) the whole-plant source sink relationship (Z. Li, Pinson, Stansel, & Paterson, 1998; Xing & Zhang, 2010; S.-M. Yu, Lo, & Ho, 2015). To increase the canopy photosynthesis of rice, a staple crop feeding half of the world population, improving plant morphology has been extensively explored in studies of genetics, and through modeling and breeding during the past 60 years (Shaobing Peng, Khush, Virk, Tang, & Zou, 2008; Q. F. Song, Zhang, & Zhu, 2013; Yonghong Wang & Li, 2008; Yuan, 2017). Recently, much progress has also been made in increasing rice canopy photosynthesis by genetically improving leaf photosynthetic rate, including engineering photosystems (J.-H. Chen et al., 2020; Khan et al., 2020; Xia Li et al., 2020; Soda et al., 2018), Rubisco (Yoon et al., 2020), photorespiratory bypass (Shen et al., 2019), stomatal conductance (Yin Wang et al., 2014) and photosynthetic related transcription factors (Ambavaram et al., 2014; Perveen et al., 2020; Shen et al., 2019; Yin Wang et al., 2014; Yoon et al., 2020). To enhance potential size of all fertile spikelets, rice geneticists and breeders have made great strides in developing rice lines bearing more ears (up to 30 per plant), or more spikelets (up to 400 per ear), or larger grains (up to 40 mg per spikelets) (Ashikari et al., 2005; X. Huang et al., 2010; Xueyong Li et al., 2003; Makino et al., 2020). Some of these rice lines have very high yield potential (Okamura et al., 2018; Sheehy, Dionora, & Mitchell, 2001). However, the filled-grain ratio of these rice lines is usually low (<85%) and hence leads to a large yield gap, i.e., large difference between the observed and the potential yields (Kato, 2004; Okamura et al., 2018; Sheehy et al., 2001; Shen et al., 2019; J. Yang & Zhang, 2010; Yoshinaga, Takai, Arai-Sanoh, Ishimaru, & Kondo, 2013).

Given certain canopy photosynthetic capacity and large potential grain sink size at flowering, designing a physiological ideotype to fill spikelets as much as possible before harvest remains a central challenge to narrow the rice yield gap (Okamura et al., 2018; S Peng, Cassman, Virmani, Sheehy, & Khush, 1999; Sheehy et al., 2001; J. Yang & Zhang, 2010). Grain filling is determined by post-anthesis whole-plant source sink relationship, which reflects the interplay between biochemical and biophysical activities in source, sink and transport organs (Fujita et al., 2013; Q. Wang et al., 2018; Wei et al., 2018; J. Yang & Zhang, 2010). Over the past decades, a large amount of descriptive data has accumulated on the dynamics of plant physiological status during rice grain filling under different environmental conditions, experimental perturbations and genetic manipulations. However, results from individual field experiments have been often disparate or even conflicting. For example, by removing part of spikelets on an ear, the percentage of filled grains had significantly increased, which suggested that poor grain filling might be due to a limitation of carbohydrates (Kato, 2004); whereas, soluble sugars were found to be higher in the usually poorly filled inferior spikelets at the early grain-filling stage, demonstrating that carbohydrates might not be the limiting factor for this process (J. Yang, Zhang, Wang, Liu, & Wang, 2006). In another case, based on the correlation between grain filling parameters and yield components in 15 different genotypes, it was concluded that the grain filling rate was much more important than the grain filling duration (Jones, Peterson, & Geng, 1979); however, observations from six field-grown tropical irrigated rice, in another study, demonstrated that the grain filling duration, instead of the grain filling rate, was significantly positively correlated with the grain yield (W. Yang, Peng, Dionisio-Sese, Laza, & Visperas, 2008). Other debates on gaining higher yield during grain filling include whether plants should have higher or lower root:shoot ratio (Ma, Li, Xu, & Huang, 2010; Nada & Abogadallah, 2016), more or less leaf nitrogen remobilization (Lee et al., 2020; Shin et al., 2020; Q. Wang et al., 2018; J. Yu, Zhen, Li, Li, & Xu, 2019), higher or lower biomass at the flowering stage (Dingkuhn, Schnier, De Datta, Dorffling, & Javellana, 1991; Kar & Kumar, 2014), and more pre-anthesis stored non-structural carbohydrates or higher post-anthesis photosynthesis (Laza, Peng, Akita, & Saka, 2003; Liu et al., 2020). These contrasting opinions reflect the complexity of interplay between source, sink and transport organs. Therefore, a comprehensive framework of source sink relationship is needed to achieve consensus on 1) what is the biochemical and biophysical basis of source sink coordination, 2) what are the key factors controlling source sink coordination, and 3) how to coordinate source and sink activities for a physiological ideotype to realize the attainable yield projected based on canopy photosynthetic potential (T-G. Chang & Zhu, 2017; Fernie et al., 2020; Paul et al., 2019; Shaobing Peng et al., 2008; Rossi, Bermudez, & Carrari, 2015; White, Rogers, Rees, & Osborne, 2015).

Systems modeling is recognized as a promising means to understand the growth of plants and guide the design of crop ideotype (T-G. Chang & Zhu, 2017; Fernie et al., 2020; Hammer, Messina, Wu, & Cooper, 2019; Marshall-Colon et al., 2017). A few attempts on building a plant-level mechanistic model have been made for wheat (Barillot, Chambon, & Andrieu, 2016), Arabidopsis (Chew et al., 2014) and barley (Grafahrend-Belau et al., 2013), which aim to simulate plant growth of certain germplasms under certain conditions or perturbations. However, these models usually have many parameters, and difficulties are encountered when we attempt to perform large scale *in silico* simulation for plant ideotype design under multiple scenarios. How to keep biological reality and parsimony in a mechanistic model simultaneously to guide future crop improvement remains a major challenge in the current crop systems modeling community (Hammer et al., 2019). To fill this gap, we have built a kinetic model of rice grain filling based on modeling whole plant carbon nitrogen interaction. We have then validated this model by simulating plant physiological behaviors under both normal conditions and six different external/endogenous perturbations during rice grain filling. By *in silico* evolutionary experiments, we have further identified both molecular targets and macro-physiological features of rice grain filling for high yield. In particular, a stable grain filling rate is found to be necessary to achieve rice yield potential under different photosynthetic capacities. We have further predicted and validated a novel *in situ* approach quantifying grain filling pattern and grain yield precisely with the measurements of ear respiratory rates. Overall, our current study provides an effective computational framework for postanthesis crop physiology research and crop ideotype design.

## RESULTS

### A theoretical framework of rice grain filling

#### The construction of a kinetic model of whole plant carbon nitrogen interaction during rice grain filling

Carbon and nitrogen metabolism is the basis of plant growth and development (Zhang, Zhou, Burnap, & Peng, 2018). To predict grain yield formation directly from the molecular processes in different source, sink and transport organs, we develop here a kinetic model of Whole plAnt Carbon Nitrogen Interaction (WACNI; **Figure 1a**). In contrast to methods that had only considered phloem sucrose transport from leaf to grain (Seki et al., 2014), or relied on preset sink growth pattern (X Yin & van Laar, 2005), our model considers both carbon and nitrogen, and kinetically simulates rates of major basic biochemical and biophysical processes in a plant using ordinary differential equations.

**Figure 1.**
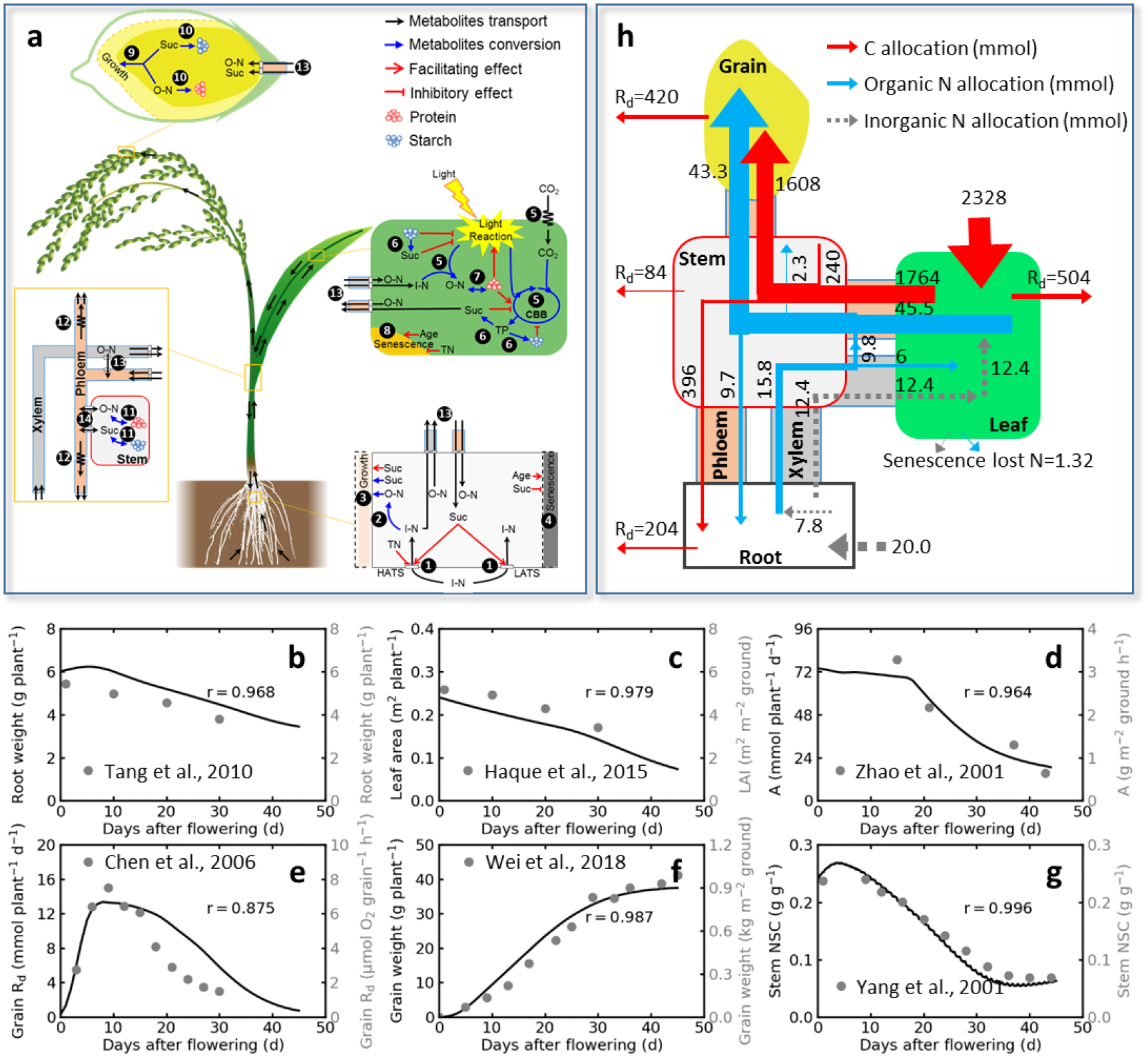
Characterizing plant physiological dynamics and carbon/nitrogen budget from flowering to harvest using a kinetic model *WACNI*. (**a**) Schematic illustration of the model. The 14 groups of simulated processes are: 1, root nitrogen uptake; 2, root nitrogen assimilation; 3, root growth; 4, root senescence; 5, leaf CO_2_ and nitrogen assimilation; 6, leaf triose phosphate, sucrose and starch interconversion; 7, leaf organic nitrogen and protein interconversion; 8, leaf senescence; 9, grain volume growth (cell division); 10, grain starch and protein synthesis; 11, stem sucrose and starch, organic nitrogen and protein interconversion; 12, phloem long distance transport; 13, transporter-dependent short distance transport; 14, symplastic diffusion between phloem and stem. (**b-g**) Plant physiological dynamics and carbon/nitrogen budget during grain filling. **b**, root dry weight (data from (Tang et al., 2010)); **c**, leaf area (data from (Haque et al., 2015)); **d**, diurnal canopy photosynthesis (data from (Q. Zhao et al., 2001)); **e**, grain respiratory rate (data from (J. Chen et al., 2006)); **f**, grain dry weight (data from (Wei et al., 2018)); **g**, stem non-structural carbohydrate content (data from (J. Yang et al., 2001)). The filled circles are measured data, the solid lines are simulated data. **h**, plant carbon and nitrogen budget throughout the grain filling period. The simulated data was from a virtual rice plant generated using default model parameter values (see details in **Supplementary Table 2**) Abbreviations: Suc, sucrose; TP, triose phosphate; I-N, inorganic nitrogen; O-N, freeform organic nitrogen; TN, total mobile nitrogen, including I-N, O-N and protein; HATS, high-affinity nitrogen transport system; LATS, low-affinity nitrogen transport system; CBB, Calvin–Benson–Bassham cycle; Rd, dark respiration; C, carbon; N, nitrogen.

The model comprises of six modules, which are the root, the leaf, the grain, the stem (including culm and sheath), the vascular transport system (xylem and phloem), and the respiration. Each module comprises a few previously well-established sub-models; and new sub-models are developed where no established sub-models are available (**Supplementary methods**; **Supplementary Table 1**). Fourteen types of biochemical and biophysical processes involved in different source, sink and transport organs are mathematically represented in the model; these processes include assimilation, transport, and utilization of six representative primary metabolites, i.e., triose phosphate (TP), sucrose (Suc), starch, inorganic nitrogen (I-N, including NH_4_^+^ and NO_3_^-^), free forms of organic nitrogen (O-N, including amino acids and amides), and proteins (**Figure 1a**). The model further incorporates known interactions between these metabolites and plant developmental processes, such as root growth, grain volume growth (endosperm cell division), grain filling (starch and protein synthesis in endosperm), root senescence, and leaf senescence (see details in **Supplementary methods**). Inputs for the model include the apparent kinetic parameters of each process, initial metabolite concentrations, and initial carbon mass and nitrogen mass in different organs at the flowering stage. A large volume of literature on rice was surveyed to generate a comprehensive parameter dataset of natural variation; and the default value of each parameter was determined empirically from the above generated dataset to form a virtual rice plant (**Supplementary Table 2**).

#### Characterizing plant physiological dynamics and carbon/nitrogen budget from flowering to harvest using the model

Firstly, the simulated physiological behaviors of the virtual rice plant successfully mimicked the real-world rice plants reported previously in multiple studies, including those on root weight (**Figure 1b**, r=0.968; r, Pearson correlation coefficient, the same below), leaf area (**Figure 1c**, r=0.979), rates of canopy photosynthesis (**Figure 1d**, r=0.964), grain respiratory rate (**Figure 1e**, r=0.875), grain dry weight (**Figure 1f**, r=0.987) and stem non-structural carbohydrates (**Figure 1g**, r=0.996).

Moreover, based on the virtual rice plant, a plant-level comprehensive carbon and nitrogen budget throughout the whole grain filling period was simulated (**Figure 1h**). For the carbon budget of a rice plant (with 12 tillers and 160 spikelets per tiller, in a canopy where plant spacing is 20 × 20 cm), it was predicted that total photosynthetic carbon intake through leaves amounting to 2328 mmol in 50 days from flowering to harvest, of which 1764 mmol was partitioned to sink organs and 504 mmol was lost through leaf respiration. Furthermore, 21.6% of the post-anthesis photosynthetically fixed carbon was predicted to be respired by the leaf itself and 52.1% by the entire plant (see detailed procedures for the calculation of values here and hereafter in **Materials and methods**), which were close to the previous estimates of ~20% and 40-60% in cultivated plants, respectively (Tcherkez & Ribas-Carbó, 2012; Whitfield, Connor, & Hall, 1989). The apparent contribution of pre-anthesis stem non-structural carbohydrate storage to grain yield, i.e., the ratio between the loss of stem dry weight during grain filling and the grain yield at harvest, was predicted to be 27.3%, which coincides with the observed values in the range of 0-40% (Yoshida, 1981). However, the actual contribution of pre-anthesis stem non-structural carbohydrate storage to grain yield, i.e., the ratio between carbohydrates produced by the remobilization of pre-anthesis stem storage and the sum of carbohydrates from remobilization and from post-anthesis photosynthesis, was much lower (12.0%). This notion also coincides with a recent field study on 12 rice cultivars from the three main rice varietal groups in China (Wei et al., 2018).

As for the nitrogen budget during grain filling, it was predicted that total nitrogen intake through root amounting to 20 mmol, accounting for 29.0% of the whole growing season nitrogen uptake, which is close to the reported value of about 30% (Ida, Ohsugi, Sasaki, Aoki, & Yamagishi, 2009). Furthermore, the nitrogen harvest index, defined as the ratio between the amount of nitrogen stored in the grain at harvest and the total plant nitrogen uptake during the whole growing season, was 62.8%, which is in the known range of 30-77% based on a large scale survey on irrigated lowland rice in tropical and subtropical Asia (Witt et al., 1999).

### Predicting plant physiological and agronomic responses to external perturbations during grain filling

In this section, rice plant physiological and agronomic responses to various external perturbations (e.g., different soil nitrogen application rates, incident light intensities and air CO_2_ concentrations) were studied *in silico* and compared to observations *in vivo*.

#### Carbon and nitrogen dynamics in leaves and grains during grain filling can be reproduced *in silico* without re-allocation of enzyme activities across different nitrogen regimes

Our model successfully reproduced carbon and nitrogen dynamics in leaves and grains throughout the grain filling period for all nitrogen treatments ranging from no nitrogen to high nitrogen, including the grain dry weight gain (**Figure 2a-d**, r=0.971-0.998), the leaf nitrogen concentration dynamics (**Supplementary Figure 1a-d**, r=0.970-0.992) and the grain nitrogen concentration dynamics (**Supplementary Figure 1e-h**, r=0.956-0.991). It is worth mentioning here that these predictions were made without adapting model parameters representing activities of metabolic and transport processes to different nitrogen regimes. Thus, these accurate predictions indicate that even if the activities of enzymes may change under different nitrogen application rates in the above study, these potential changes are not the major players in determining carbon and nitrogen dynamics in leaves and grains during grain filling.

**Figure 2.**
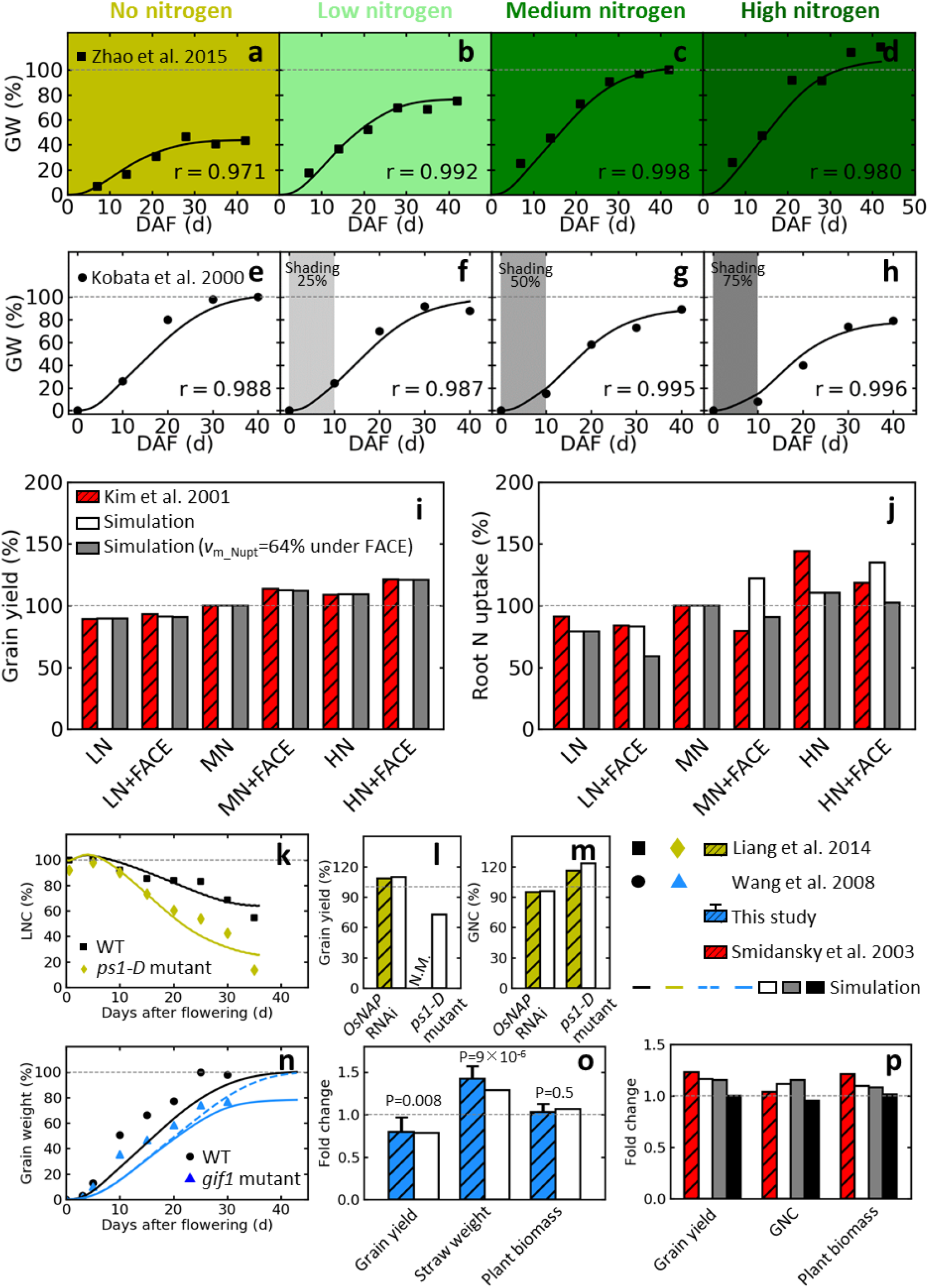
Predicting plant physiological and agronomic responses to external and endogenous perturbations during grain filling. (**a-d**) Grain filling under different nitrogen application rates. **a**, no nitrogen; **b**, low level nitrogen; **c**, medium level nitrogen; and **d**, high level nitrogen. The dynamic changes of leaf and grain nitrogen concentrations are shown in **Supplementary Figure 1**. The filled squares are measured data from Y Zhao et al. (2015), the solid lines are simulated data. Data were normalized by dividing by the final grain weight in the “medium nitrogen” group. (**e-h**) Grain filling under different light regimes. **e**, normal condition; conditions of shading 25% (**f**), 50% (**g**), and 75% (**h**) of the solar radiation during the first 10 days after flowering. Results on shading treatments followed by spacing are shown in **Supplementary Figure 2**. The filled circles are measured data from Kobata et al. (2000), the solid lines are simulated data. Data were normalized by dividing by the final grain weight under normal condition. (**i-j**) Grain yield and root nitrogen uptake during grain filling for plants grown under different air CO_2_ concentrations and soil nitrogen application rates. The red/striped bars are measured data from Kim, Lieffering, Miura, Kobayashi, and Okada (2001), the white bars are simulated data, the grey bars are simulated data with the additional assumption of the maximum root nitrogen uptake rate under FACE conditions being 64% of that under ambient conditions. Data were normalized by dividing by the values in the “MN” group. **(k-m)** Grain yield, GNC and dynamic change of LNC of the wild type (WT), the *OsNAP* gain-of-function mutant (*ps1-D*) with faster leaf senescence, and the *OsNAP* RNAi with slower leaf senescence. The black lines in panel a are simulated values for WT and yellow lines for *ps1-D* mutant. Data were normalized by dividing by the values of WT at flowering (for LNC) or at harvest (for grain yield and GNC). **(n-o)** Grain filling and agronomic traits of the wild type (WT) and the *gif1* mutant with impaired grain sucrose unloading. Data were normalized by dividing by the values of WT at harvest. The dashed blue line in panel n illustrates predicted grain filling of a hypothetical “new *gif1* mutant” with the same grain sink size to that of WT. Experimental data in panel o are mean ± sd (n=10); and the*p*-values show the probability between the difference of mean values in the *gif1* mutant and in the WT, and were calculated from two-tailed *t*-test. (**p**) Grain yield, GNC and aboveground plant biomass of the *Sh2r6hs* transgenic plant with enhanced endosperm ADP-glucose pyrophosphorylase activity. The grey bars are simulated values by changing only the plant size at the flowering stage for the *Sh2r6hs* transgenic plant; the black bars are simulated values by changing only *ν*_m_grain_Star_syn_ during grain filling; the white bars are simulated values by changing both the plant size and *ν*_m_grain_Star_syn_. Abbreviations: DAF, days after flowering; FACE, free air CO_2_ enrichment; GNC, grain nitrogen concentration; HN, high nitrogen; GW, grain dry weight; LNC, leaf nitrogen concentration; LN, low nitrogen; MN, medium nitrogen; N.M., not measured.

#### Low light during early grain filling period affects grain weight gain pattern but does NOT compromise rice yield potential

For grain filling under different light regimes, an increased penalty for grain dry weight 10 days after flowering was predicted, as well as at harvest, when 25%, 50% or 75% of the solar irradiance was shaded during the first 10 days after flowering (**Figure 2e-h**, r=0.987-0.996). In addition, our model correctly predicted an acceleration of grain filling beyond 10 days after flowering when extra light penetrates into the canopy as a result of plant thinning (**Supplementary Figure 2b-d**, r=0.994-0.999), supporting the idea that short-term impaired grain filling by low light-induced shortage of assimilates can almost be fully compensated if sufficient resource is provided later (Kobata, Sugawara, & Takatu, 2000).

#### Inhibited root nitrogen uptake capacity during grain filling under elevated air CO2 concentration underlies reduced plant nitrogen budget

For post-anthesis rice plants grown under different atmospheric CO_2_ concentrations and soil nitrogen application rates, our model successfully reproduced the enhancement of grain yield with increasing nitrogen application rate under both ambient CO_2_ level and free air CO_2_ enrichment (FACE; the white bars in **Figure 2i**; r=0.992 is between the white bars and the red/striped bars). However, when plants with the same nitrogen application rates were compared, our model predicted higher total root nitrogen uptake during the grain filling period of plants grown under FACE than that of plants grown under ambient CO_2_ concentration (the white bars in **Figure 2j**), which is inconsistent with the experimental observations (the red/striped bars in **Figure 2j**; r=0.318 is between the white bars and the red/striped bars). Earlier, it was shown that the maximum nitrogen uptake rate per root dry weight under FACE was reduced by 31-41% from the flowering stage to the harvest stage compared to that of plants grown under ambient CO_2_ concentration (Shimono & Bunce, 2009). Following this observation, if mean percentage reduction of maximum root nitrogen uptake rate, of 36%, was set under FACE in the model, the response pattern of root nitrogen uptake under FACE, during the grain filling period, could be explained much better (see the grey bars in **Figure 2j**; r=0.713 is between the grey bars and the red/striped bars). These results highlight the important role of root nitrogen uptake activity in affecting plant nitrogen budget under FACE during grain filling.

### Predicting plant physiological and agronomic responses to endogenous perturbations during grain filling

In this section, we present our data on plant physiological and agronomic responses of rice to various endogenous perturbations in source, transport and sink organs.

#### Prolonged leaf functional duration benefits grain yield with a slight reduction in grain nitrogen concentration by enhancing later-stage photosynthesis

We first simulated impacts on grain yield related traits when senescence of the major carbon source, leaf, was perturbed. Based on the work of Liang et al. (2014), three scenarios were examined, and we indeed showed: (i) the enhanced leaf protein degradation activity mimicking a dominant premature leaf senescence mutant, prematurely senile 1 (the “*ps1-D* mutant” group in **Figure 2k-m**); (ii) the medium leaf protein degradation activity mimicking the wild type plant (the “WT” group in **Figure 2k-m**); and (iii) the reduced leaf protein degradation activity mimicking the NAC-like, activated by apetala3/pistillata (*OsNAP*) gene RNAi plant (the “*OsNAP* RNAi” group in **Figure 2l-m**). A faster leaf nitrogen loss in the first scenario was predicted due to accelerated leaf protein degradation rate, mimicking the observed faster decrease in chlorophyll content of the *ps1-D* mutant (**Figure 2k**). Compared to WT, an increase of grain yield at the expense of a slight decrease in grain nitrogen concentration for the *OsNAP* RNAi plant, as well as the opposite for the *ps1-D* mutant, was predicted, coinciding well with the experimental observations (**Figure 2l, m**). A comprehensive characterization of source, sink and transport activities in these plants further suggested that the enhanced grain yield of the *OsNAP* RNAi plant was mainly contributed by the higher canopy photosynthesis in the later grain filling period with higher leaf nitrogen concentration, whereas the slightly reduced grain nitrogen concentration was mainly attributed to the decreased nitrogen concentration in the phloem with less leaf nitrogen remobilization (**Supplementary Figure 3; Figure 2k**).

#### Phloem-to-grain sucrose unloading capacity is NOT necessarily positively correlated with grain yield

We then simulated consequences of perturbing metabolite transport. E. Wang et al. (2008) had reported that a gene, GRAIN INCOMPLETE FILLING 1 (*GIF1*), which encodes a cell-wall invertase, is required for sucrose transport to the grain. In agreement with the experimental result, grain dry weight gain was predicted to be slower for the *gif1* mutant, which had a decreased maximum phloem-to-grain sucrose unloading rate (*ν*_m_grain_Suc_ul_; **Figure 2n**). Furthermore, it was predicted that although the grain yield was impaired in the *gif1* mutant, its straw weight (a sum of weight of leaves and stems) was higher, which compensated for the decrease of grain yield and led to a similar aboveground plant biomass of the *gif1* mutant with that of the WT at harvest (the white bars in **Figure 2o**). Although relevant data was not reported in the original paper, results from our field experiments using seeds provided by the authors has confirmed this prediction (**Figure 2o**). Namely, accumulation of photosynthates during grain filling was not impaired in the *gif1* mutant. Decrease of grain yield in the *gif1* mutant may be the result of sink-size limitation, as the dry weight of a single grain had decreased in the *gif1* mutant (E. Wang et al., 2008). To test this hypothesis, we generated a hypothetical “new *gif1* mutant” with the same grain sink size to that of WT. As a result, this “new *gif1* mutant” gained a similar grain yield to that of WT, although with a slower grain filling rate (dashed line in **Figure 2n**). Additionally, although E. Wang et al. (2008) had proposed that overexpressing the *GIF1* gene might increase the yield potential through improved grain-filling, the grain yield of *GIF1* overexpression plants was not presented. Using our model, we predicted that when plants were sink limited, increasing *ν*_m_grain_Suc_ul_ must enhance both the grain filling rate and the grain yield, whereas decreasing *ν*_m_grain_Suc_ul_ must impair the grain filling rate, as well as the grain yield (**Supplementary Figure 4**). Nevertheless, when plants are source limited, increasing *ν*_m_grain_Suc_ul_ might not increase yield, but even decrease it, mainly due to a faster sink growth induced accelerated leaf nitrogen remobilization, and consequently earlier leaf senescence (**Supplementary Figure 5**). Consistent with this notion, overexpression of *PbSWEET4*, a sugar transporter gene in pear, is known to cause early senescence in leaves (Ni et al., 2020). Similarly, overexpression of *OsSWEET5* in rice, which encodes a sugar transporter protein in leaves, stem, root and floral organs, has been shown to cause precocious leaf senescence (Zhou et al., 2014).

#### Enhanced endosperm ADP-glucose pyrophosphorylase activity cannot fully explain the mechanism for rice with ectopic expression of Sh2r6hs to gain increased grain yield

We finally simulated consequences of altering the activity of grain storage metabolism. Endosperm-specific ectopic expression of a modified maize ADP-glucose pyrophosphorylase (AGPase) large subunit sequence (*Sh2r6hs*) had been shown to enhance the activity of starch synthesis in the endosperm of both wheat and rice (Eric D Smidansky et al., 2002; E. D. Smidansky, Martin, Hannah, Fischer, & Giroux, 2003). Compared to that in WT, the predicted changes of the grain yield, the grain nitrogen concentration and the aboveground plant biomass of the *in silico Sh2r6hs* transgenic plant (the white bars in **Figure 2p**) matched the experimental observations (the red/striped bars in **Figure 2p**). Further, we noted that although the *Sh2r6hs* was expressed under the control of an endosperm-specific promoter, the size of the transgenic plant at flowering was shown to differ from that of WT. Thus, we asked whether the increase of the grain yield was mainly caused by the increased rate of maximum grain starch synthesis (*ν*_m_grain_Star_syn_) during grain filling, or was it mainly caused by the plant size change at the flowering stage. To answer this question, we simulated three different scenarios for the *Sh2r6hs* transgenic plant: (i) we changed its *ν*_m_ grain_Star_syn_ during grain filling only (the black bars in **Figure 2p**); (ii) we changed its plant size at the flowering stage only (the grey bars in **Figure 2p**); (iii) we changed both (the white bars in **Figure 2p**). As a result, simulations matched experimental observations for both the second and the third scenarios, but not for the first scenario (**Figure 2p**). These data demonstrate that the increased grain yield in the *Sh2r6hs* transgenic plant is mainly contributed by a change of the plant size at the flowering stage, rather than being mainly contributed by the ~50% enhancement of the maximum activity of endosperm starch synthesis during grain filling. In support of this notion, there is increasing evidence that crop yield responses are largely determined by the role of AGPase in affecting vegetative tissues and the number of grains formed at flowering (Fahy et al., 2018; Hannah et al., 2012; Saripalli & Gupta, 2015).

### *In silico* design of rice grain filling ideotype for super-high yield

#### Influences of capacities of reaction and diffusion processes on grain yield and yield-related agronomic traits are diverse

To systematically explore the influences of basic reaction and diffusion processes on post-anthesis physiology, we performed sensitivity analysis on 28 parameters controlling the maximum rate of reaction and diffusion processes. We classified these 28 parameters into four categories, based on the predicted response patterns of grain yield to parameter changes. Specifically, (i) eleven parameters that increase grain yield, when they increase from 0.1-fold to 10-fold of their default values, were termed as universal yield enhancers (UYEs); (ii) four parameters that decrease grain yield, when they increase from 0.1-fold to 10-fold of their default values, were termed as universal yield inhibitors (UYIs); (iii) six parameters that non-linearly affect grain yield were termed as “conditional yield enhancers” (CYEs); (iv) seven parameters having negligible influence on grain yield were termed as “weak yield regulators” (WYRs) (see grain yield response to typical UYE, UYI, CYE and WYR in **Figure 3a-d**, respectively; see grain yield response to all parameters in **Supplementary Figure 6**). Besides grain yield, responses of other seven major agronomic traits, i.e., grain nitrogen concentration, photosynthesis, root nitrogen uptake, grain filling rate, grain filling duration, harvest index, and nitrogen harvest index, upon 0.1-fold to 10-fold change of the 28 parameters, are documented in details in **Supplementary Figures 7-13**.

**Figure 3.**
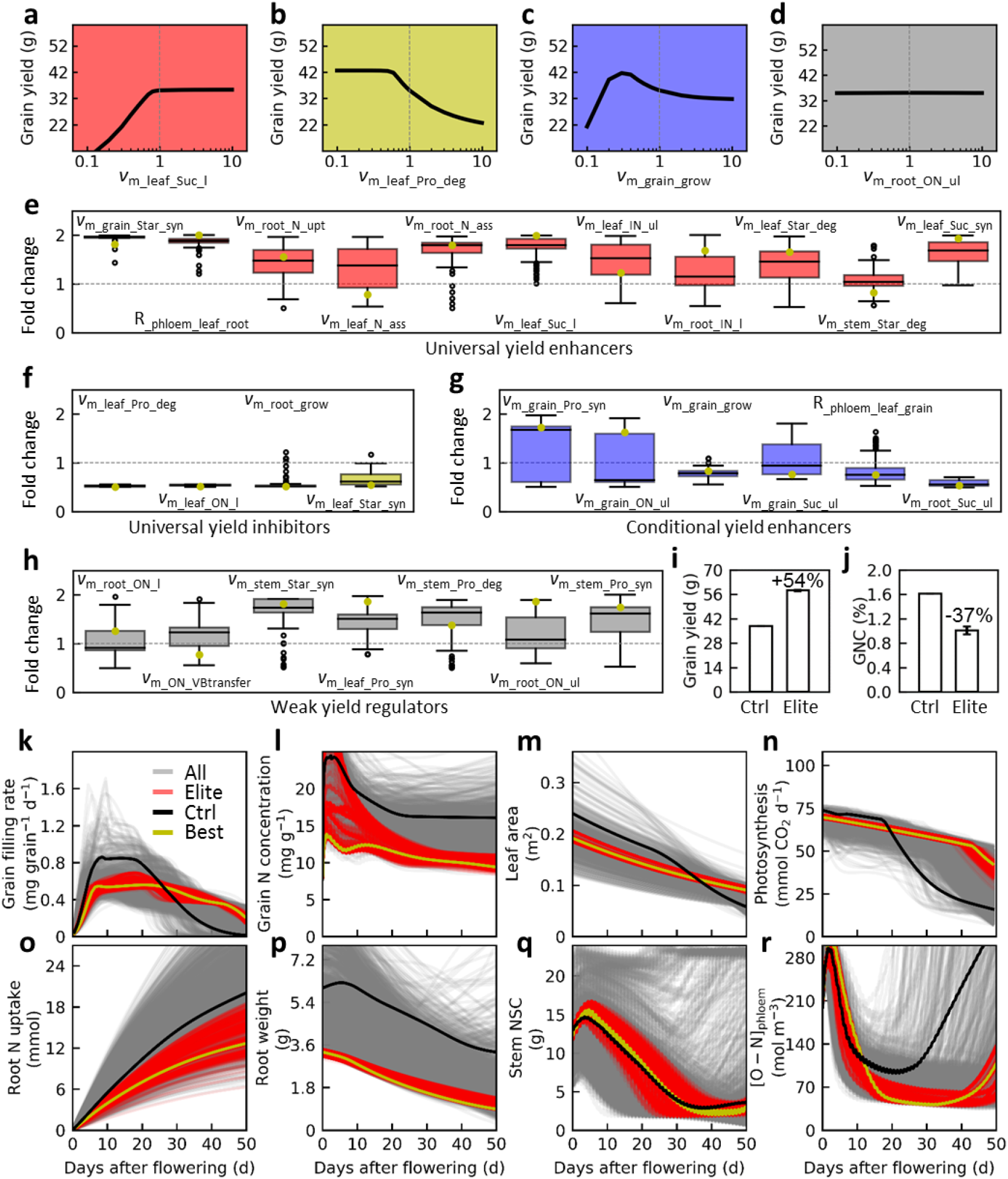
Designing rice physiological ideotype for super-high yield. **(a-d)** Predicting and grouping the effects of manipulating capacities of reaction and diffusion processes on grain yield and yield-related agronomic traits. **a**, leaf sucrose loading capacity (*ν*_m_leaf_Suc_l_) shows a universal yield enhancer which can monotonically increase grain yield when its value ranges from [0.1, 10]-fold to its default value. **b**, leaf protein degradation capacity *ν*_m_leaf_Pro_deg_ shows a universal yield inhibitor which can monotonically decrease grain yield. **c**, grain growth capacity *ν*_m_grain_grow_ shows a conditional yield enhancer which can non-linearly influence grain yield. **d**, root organic nitrogen unloading capacity *ν*_m_root_ON_ul_ shows a weak yield regulator which has negligible effects on grain yield. **(e-h)** Distribution of biochemical and biophysical parameter fold changes for *in silico* elite rice individuals with super-high yield (n=500). The parameters include universal yield enhancers (**a**), universal yield inhibitors (**b**), conditional yield enhancers (**c**) and weak yield regulators (**d**). The yellow dots illustrate parameter values for the highest yielding individual among elite individuals. The parameter fold change was calculated by dividing by the default value of a parameter. **(i-j)** Comparison of grain yield per plant (**i**) and grain nitrogen concentration (**j**) for elite individuals (Elite) and the control individual with default parameter values (Ctrl). **(k-r)** Comparison of macroscopic trait changes of the control individual, elite individuals, the highest yielding individual (Best), and all individuals (All) generated during *in silico* evolution. Macroscopic traits include single-grain filling rate (**k**), grain nitrogen concentration (**l**), leaf area (**m**), photosynthetic rate (**n**), total accumulated root nitrogen uptake (**o**), root weight (**p**), amount of stem non-structural carbohydrates (NSC; **q**), and phloem organic nitrogen concentration ([O-N]_phloem_; **r**). Parameter values for elite individuals are listed in **Supplementary Datasets 1**.

#### Designing rice physiological ideotype for super-high yield

To search for optimal combinations of parameter values maximizing grain yield, we combined our model with a genetic algorithm. Briefly, we first randomly varied the values of model parameters in a range from 50% to 200% to their default values to generate five independent populations; then, we “evolved” these five populations using high grain yield as the selection target (**Supplementary Figure 14**). The top 1% individuals ranking by grain yield in each population were then merged to form an elite plant group with 500 rice individuals (the “Elite” group in **Figure 3** and **Supplementary Figures 15-16**). In general, for individuals in the elite group, the parameter values of UYEs were up-tuned (**Figure 3e**), the parameter values of UYIs were down-tuned (**Figure 3f**), but the parameter values of CYEs and WYRs showed no specific pattern (**Figure 3g, h**). Further, the initial partitioning of carbon and nitrogen among organs at the flowering stage for the elite individuals was distinct from the default values. As an example, the grain number increased while the root size decreased dramatically for elite individuals (**Supplementary Figure 15**). Finally, as compared to the control individuals, the *in silico* optimization resulted in a remarkable (54%) increase of grain yield, at the expense of a 37% decrease in grain nitrogen concentration (**Figure 3i-j**).

Elite individuals show distinct patterns of macroscopic physiological changes compared to other individuals. For example, the rate of single-grain filling in the elite individuals was relatively low during the first 7 days after flowering, and was moderate in 7 to 25 days after flowering; remarkably, it decreased at much slower pace, starting around 20 days after flowering to the harvest stage, when compared to the control and most other individuals (**Figure 3k**). This stable grain filling rate resulted in a nearly linear grain yield gain for the elite individuals rather than the sigmoid curves, as observed for the control and most other individuals (**Supplementary Figure 16**). Furthermore, the grain nitrogen concentration was lower for elite individuals (**Figure 3l**), and they had slower rate of decrease in both leaf area and photosynthetic rate (**Figure 3m-n**), a feature commonly known as *stay green* (Thomas & Ougham, 2014). Although elite individuals had much smaller root mass throughout the grain filling period (**Figure 3p**), some elite individuals had similar root nitrogen uptake as the control individual did (**Figure 3o**), as a result, we suggest, of their higher maximum root nitrogen uptake rate (*ν*_m_root_N_upt_ in **Figure 3e**). Finally, elite individuals used up most of the stem non-structural carbohydrate at harvest (**Figure 3q**), as well as maintained a low level of organic nitrogen concentration in phloem ([O-N]_phloem_) from around 10 days after flowering to the harvest stage (**Figure 3r**).

### Stable grain filling rate is required to achieve rice yield potential under different photosynthetic capacities

#### An inherent relation between grain filling rate, grain respiratory rate and grain yield

The simulations, presented above, show that to gain super-high yield, grain filling rate, both on a grain basis and on a land area basis, should not be too high or too low in the early grain filling period, whereas they should be maintained in the mid-grain filling period and decrease as slowly as possible in the late grain filling period (**Figure 3g; Figure S16**). To confirm this notion, we grew distinct rice lines in different eco-zones (in total:14 groups; **Supplementary Figure 17**). The rice lines used in this research included *japonica*, *indica* inbred rice cultivars, as well as hybrid rice cultivars, and produced dried brown grain yield of 5 to 15 t ha^−1^ (**Supplementary Figure 17**). Intriguingly, both the distinct super-high yielding rice cultivars had almost identical grain yield gain pattern to that predicted by *in silico* analysis (**Figure 4a**). To further explore the connection between the rate of grain filling and the grain yield, we studied the relation between mean/instantaneous grain filling rates and the grain yield, using all 50,000 rice individuals from the above mentioned five *in silico* evolutionary populations, and examined these findings using all 14 groups of field data. We found: (i) mean grain filling rates of ≤15 days after flowering were negatively or hardly correlated with the grain yield, whereas the same rates ≥22 days after flowering were significantly positively correlated with the grain yield (**Figure 4b, c**, r>0.68, P<0.01); (ii) instantaneous grain filling rate at 36 days after flowering were strongly positively correlated with the grain yield (**Figure 4d**, r=0.89); (iii) a sum of instantaneous grain filling rates at 15 and 38 days after flowering accurately predicted the grain yield (**Figure 4e**, r=0.96).

**Figure 4.**
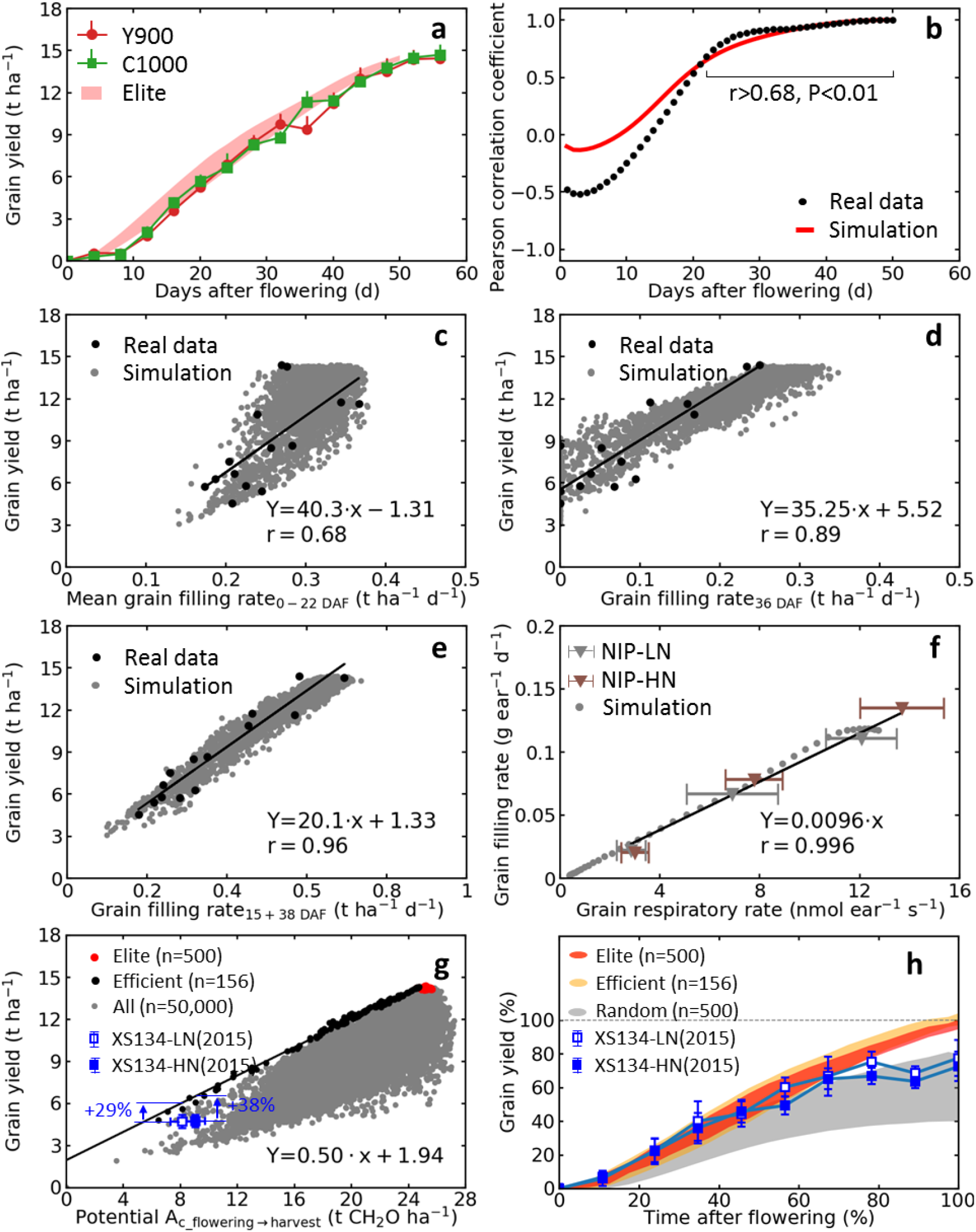
Stable grain filling rate is required to achieve rice yield potential under different photosynthetic capacities. **(a)** Grain filling pattern of the *in silico* elite individuals (Elite) and two hybrid rice cultivars *Y-Liang-You 900* (*Y900*) and *Chao You 1000* (*C1000*) grown in Yongsheng county, Yunnan province, China that reach super-high dried brown grain yield of about 15 t ha^−1^. **(b-c)** Relation between mean grain filling rates during the first *i* days after flowering and the grain yield. **b**, Pearson correlation coefficients between mean grain filling rate during the first *i* days after flowering and grain yield. The correlations were significant at a level of P<0.01 from 22 days after flowering till harvest. **c**, an illustration of correlation between the mean grain filling rate during the first 22 days after flowering and grain yield. **(d-e)** Relation between instantaneous grain filling rates and the grain yield. **d**, correlation between instantaneous grain filling rate on the 36^th^ day after flowering and grain yield. **e**, correlation between grain yield and a sum of instantaneous grain filling rates on the 15^th^ and 38^th^ day after flowering. **(f)** Relation between instantaneous grain filling rates and grain respiratory rates (mean ± sd, n=5). For the “Simulation” group, data from the *in silico* control individual were used. **(g)** Relation between potential photosynthate accumulation from flowering to harvest and grain yield. The potential photosynthate accumulation is an integration of daily potential gross photosynthesis throughout the grain filling period; and daily potential gross photosynthesis is calculated based photosynthetic capacity at the flowering stage. The *in silico* “Elite” and “All” individuals were defined in **Figure 3**; and the *in silico* “Efficient” individuals refer to individuals having highest grain yields with different potential photosynthate accumulation. Experimentally, the *japonica* rice cultivar *Xiu-Shui 134* grown under low nitrogen (*XS134-LN*) and high nitrogen (*XS134-HN*) treatments in 2015 were shown (mean ± sd, n=5). **(h)** Grain filling pattern of the *in silico* and *in vivo* rices. Grain filling duration and the potential grain yield are normalized to 1 for all rice individuals. The potential grain yields were calculated based on potential photosynthate accumulation using the equation shown in panel g. The filled regions represent different groups of *in silico* individuals with 95% confidence intervals (mean ± 1.96sd). The *in silico* “Random” individuals refer to individuals in the first generations of five evolutionary populations which were generated by randomly varying the values of the parameters. Experimentally, the grain filling patterns of the *XS134-LN* and *XS134-HN* treatments in 2015 were shown (mean ± sd, n=5).

Given the close association between the grain filling rate and the grain yield, how should we quantify grain filling rate *in situ*? Again, from the *in silico* simulation, we noticed a similar change pattern of the grain filling rate and the grain respiratory rate beyond 10 days after flowering, when active grain filling takes place (filled grey circles in **Figure 4f**). To test this finding, we measured respiratory rates of an intact ear at 12, 17 and 21 days after flowering in *Nipponbare*, a rice cultivar grown under two nitrogen conditions, using a custom-built cuvette (Tian-Gen Chang et al., 2020). Here, we observe that the correlation between the rates of grain respiration and grain filling is indeed extremely strong (**Figure 4f**, r=0.996). Given the strong correlation between the grain respiratory rate and the grain filling rate, logically, the sum of grain respiratory rates at 15 and 38 days after flowering should predict the grain yield as what has been proven for grain filling rates (**Figure 4e**). In agreement of the above, a strong correlation between ear respiratory rates at 15 and 38 days after flowering and ear dry weight at harvest was consistently found (**Supplementary Figure 18**, r=0.933).

#### Maximizing grain yield under different photosynthetic capacities

We further asked as to what could be the upper limits of grain yield for rice with different photosynthetic capacities at flowering and different grain filling durations. To answer this question, we studied the relation between grain yield and potential photosynthate accumulation from the time of flowering to harvest, using the 50,000 *in silico* rice individuals (**Figure 4g**). Knowing that the potential photosynthate accumulation is an integration of daily potential gross photosynthesis throughout the grain filling period, we identified 156 individuals with highest yields under different levels of potential photosynthate accumulation (the “Efficient” group in **Figure 4g**). We further deduced an expression converting potential photosynthate accumulation *x* to potential grain yield *Y* using these efficient individuals: *Y*=0.50*x*+1.94 (**Figure 4g**). Based on this calculation, the grain yields could be increased to reach the potential grain yields for the elite *japonica* rice cultivar under low nitrogen (by 29%) and high nitrogen (by 38%) conditions (**Figure 4g**).

We further studied the key physiological features of the efficient individuals (**Supplementary Figure 19**). As the efficient individuals have very different grain yields from each other, we normalized the grain filling duration and the potential grain yield to 1. Remarkably, the efficient individuals, as well as the elite individuals, had a stable grain filling rate from flowering to harvest, whereas, as expected, the randomly generated *in silico* individuals and the field-grown rice cultivar did not show this grain filling pattern (**Figure 4h**).

## DISCUSSION

Despite the descriptive research of source sink interaction in single cells or organs, the plant level coordination of processes in different source, transport and sink organs has been less studied, most likely, due to difficulties in interpretation of data in such a complex network (Sonnewald et al., 2020). Crop systems models have been used, in the past, in designing agronomic practices (Deng et al., 2012; Y. Wang et al., 2017), testing hypotheses regarding the responses of crops to climate change (Chenu et al., 2017; Q. Song, Srinivasan, Long, & Zhu, 2019), guiding selection of traits for breeding (Tuberosa, 2012) and assessing different strategies to adapt to future climate change (B. Peng, Guan, Tang, Ainsworth, & Zhou, 2020). Here, we have introduced WACNI, a new computational model of rice grain filling. Although as a highly simplified theoretical framework, it is remarkable that the *in silico* model reproduces major physiological changes *in vivo* under different conditions and perturbations (**Figures 1, 2; Supplementary Figures 1, 2**), thus providing a comprehensive view of rice grain filling. A few distinct features of the model may underlie its superior performance under multiple scenarios. First of all, the model simulates plant level physiological dynamic changes during grain filling, as an emergent property, from basic biochemical and biophysical processes of carbon and nitrogen in different plant organs. This *bottom-up* mechanistic approach clearly differs from the approaches using either preset rules on the relation between the source and sink organs, or the empirical correlation derived from the measured data. Concurrently, a comprehensive survey was conducted on the natural variation of the morphological and physiological traits in rice from a large amount of studies rather than from a single source, thus enabling an unbiased estimation of the model parameters (**Supplementary Table 2**). Last but not least, although being simplified, the model essentially embraces all plant organs, major primary metabolism of both carbon and nitrogen, and long-established feed-back and feed-forward regulations of activities in source and sink organs, thus allowing it to largely reproduce the complicated source sink interactions during rice grain filling (see **Supplementary methods**). Currently, the model has not been thoroughly parameterized for specific rice cultivars due to the incompleteness of data. The cultivar-specific parameterization will be greatly facilitated by the emerging of comprehensive datasets on whole plant level metabolic fluxes based on fluxomics (Salon et al., 2017). In addition, since many processes described in the model are rather generic among plants, the model may be adapted for other plants, especially for other cereal crops, such as wheat and maize. Finally, the framework presented here may support future development of more sophisticated crop systems models charactering plant signal transduction and regulation (S.-M. Yu et al., 2015), or elaborating 3D structures and functions of roots and shoots (Tian-Gen Chang, Zhao, et al., 2019; Postma et al., 2017; Watanabe et al., 2005).

With the theoretical framework, we may, indeed, revisit the potential targets for higher crop yield during grain filling that have been discussed for over four decades, such as functional *stay green* of leaves, higher phloem-to-grain sucrose unloading rate, higher leaf sucrose export rate, higher endosperm starch synthesis activity, more efficient remobilization of stem starch reserves, higher grain filling rate, and longer grain filling duration (Braun, Wang, & Ruan, 2014; Fahy et al., 2018; Jones et al., 1979; Saripalli & Gupta, 2015; Thomas & Howarth, 2000; W. Yang et al., 2008). In brief, our theoretical framework helps us predict non-intuitive plant physiological and agronomic responses when parameters controlling plant morphology and metabolism are changed by endogenous or external factors or both; evaluate the impacts on plant physiological dynamics of each of the multiple changes in plant morphology and physiology produced by a single endogenous or external perturbation; and identify the major mechanism underlying grain yield response (**Figure 2; Supplementary Figures 1-5**). All these results demonstrate the promise of mechanistic modelling of providing more in-depth insight into biological mechanisms, rather than merely serving as a plant growth simulator.

By linking basic biochemical and biophysical processes to plant physiological changes all the way up to agronomic traits, our framework offers an opportunity to design grain filling ideotype for maximizing crop yield. Given a gross canopy photosynthesis capacity of ~1750 mmol m^−2^ d^−1^ (**Figure 3n**, ~70 mmol per plant d^−1^ and 25 plants m^−2^) at flowering, an optimization of source sink and transport related processes *in silico* results in a standard yield of up to 21 t ha^−1^ (“rough” rice, which has 14% moisture content and 20% husk weight (Ahmaruzzaman & Gupta, 2011; Yoshinaga et al., 2013)). In such a rice line, a balanced source sink relationship is realized from re-allocation of the initial partitioning of carbon and nitrogen among organs and the fine-tuning of capacities of different basic metabolic processes (**Figure 3e-h; Supplementary Figure 15**). The optimal grain filling pattern leading to the standard rice yield of 21 t ha^−1^, designed *in silico*, is validated *in vivo* by using state-of-the-art high-yield cultivars (China super hybrid rice cultivar *Y-Liang-You 900* and *Chao You 1000*)(Cheung, 2014; M. Huang et al., 2017) and growing them in one of the most suitable eco-zones for super-high yield of rice (Yongsheng county, Yunnan province, China)(Z. Chen, Wang, Wen, & Wang, 2007) (**Figure 4a**). We emphasize that 21 t ha^−1^ is not the ultimate limit of rice yield, since, e.g., a further increase in canopy photosynthesis at flowering and extension of grain filling duration may lead to higher yields.

More generally, for maximizing grain yields with different photosynthetic capacities at flowering, stable grain filling rate, slower decrease of photosynthetic capacity, and full use of stem non-structural carbohydrates are essential macro-physiological features (**Figure 3k-r; Supplementary Figure 19**). Potential common post-anthesis molecular targets include delaying leaf senescence (decreasing *ν*_m_leaf_Pro_deg_ and *ν*_m_leaf_ON_l_), enhancing leaf sucrose synthesis and export (increasing *ν*_m_leaf_Suc_l_ and *ν*_m_leaf_Suc_syn_), limiting root sucrose consumption (increasing R_ phloem_leaf_root and decreasing *ν*_m_root_Suc_ul_), strengthening stem starch synthesis (increasing *ν*_m_stem_Star_syn_), accelerating endosperm starch synthesis (increasing *ν*_m_grain_Star_syn_), and moderating endosperm cell division (fine-tuning *ν*_m_grain_grow_) (**Supplementary Figure 20**; **Figure 3e-h**). Mining more genes and quantitative trait loci (QTLs) related to these molecular targets (**Supplementary Table 3**), stacking superior alleles of these genes and QTLs using advanced genomics technology (Qian et al., 2016), and developing smart and precision crop cultivation route (Saiz-Rubio & Rovira-Más, 2020) are promising approaches to realize the designed physiological ideotype. Theoretically, optimizing grain filling patterns in an elite rice cultivar under different nitrogen treatments may increase its grain yield by about 30-40% (**Figure 4g, h**), implying the huge potential of coordinating source sink relationship during grain filling for higher yield. Given the fact that it is no longer an obstacle to create sufficient amount of spikelets at flowering (Kato, 2004; Okamura et al., 2018; S Peng et al., 1999; Qian et al., 2016; J. Yang & Zhang, 2010), optimizing grain filling pattern is a major task for greater yield in the future beyond the current focus of improving canopy photosynthesis. In this aspect, measuring ear respiratory rates, especially around 15 and 38 days after flowering, can serve as a novel efficient approach for field *in situ* phenotyping of grain filling pattern and grain yield (**Figure 4f; Supplementary Figure 18**). In view of rapid advances in rice functional genomic and genome editing technologies (Y. Li et al., 2018; Nadakuduti & Enciso-Rodríguez, 2021), greater understanding of carbon and nitrogen fluxes between different organs and plant physiological changes during grain filling could pave the way for rational design of high-yield crops.

## Materials and methods

### Model development

The model developed here is depicted diagrammatically in **Figure 1a**. Essentially, after the reaction diagrams were established, differential equations, rate equations, and algebraic equations representing conserved quantities were developed. The model was implemented in MATLAB and solved using ode15s (the MathWorks Inc., Natick, MA, USA). The model computed changes in the metabolite concentrations by the differences between rates of fluxes generating and consuming metabolites.

Fourteen different types of primary biochemical/biophysical processes were incorporated in the model (**Figure 1a**). The rate equations of these processes in different root, leaf, grain, and stem organs were described in detail in **Supplementary methods**.

### Model parameterization

The maximum velocities (*ν*_max_) for processes simulated in the model were derived from a vast amount of genetic, molecular, biochemical and plant physiological research in rice; while most of the apparent Michaelis-constants (*K*_m_) and equilibrium constants (*K*_e_) were estimated based on initial metabolite concentrations in each organ. A detailed description of the parameters, their values, and the comprehensive parameterization procedures used are documented in **Supplementary Table 2**.

### Model performance on multiple datasets

#### Simulating typical plant physiological changes during rice grain filling (Figure 1b-g)

The model was validated by comparing the relative changes of physiological traits in different organs during the grain filling period between model predictions with default parameter settings and the experimental data obtained from literature under normal conditions (**Figure 1b-g**). Specifically, the dynamic change of root weight (**Figure 1b**) was extracted from *Table 2* in Tang et al. (2010), and the average values of seven combinations of two-line hybrid rice were used; the dynamic change of leaf area (**Figure 1c**) was extracted from *Figure 1(d)* in Haque, Pramanik, Biswas, Iftekharuddaula, and Hasanuzzaman (2015), and the average values of three hybrid and inbred varieties in 2008-2009 were used; the dynamic change of canopy photosynthesis (**Figure 1d**) was extracted from *Fig. b* in Q. Zhao, Huang, and Ling (2001), and the average values of treatments F1-F7 were used; the dynamic change of grain respiration (**Figure 1e**) was extracted from *Fig. 3* in J. Chen, Wang, Chen, Wang, and MO (2006), and the average values of cultivars *Dali* and *IR36* in the control groups were used; the dynamic change of grain dry weight (**Figure 1f**) was extracted from *Fig. 2* in Wei et al. (2018), and the average values of cultivars Yongyou 13/17 grown in 2013 were used; the dynamic change of stem non-structural carbohydrates (**Figure 1g**) was extracted from *Fig. 4B* in J. Yang, Zhang, Wang, Zhu, and Wang (2001), and the average values of normal nitrogen with well-watered and water deficit stressed treatments of cultivar *Yangdao 4* were used. All the extracted experimental data are tabulated in **Supplementary Datasets 2**.

#### Simulating plant carbon and nitrogen allocation during rice grain filling (Figure 1h)

Specifically, plant total dry mass at flowering (PTM0) is calculated as:

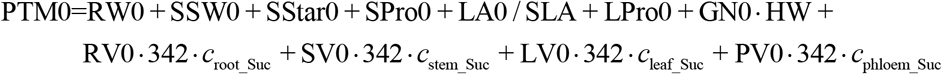

where, RV0, LV0 and PV0 are root volume, leaf volume and phloem volume, respectively; *c*_x_y_ are the concentration of metabolite *y* in organ *x* (mol m^−3^). Other parameters are defined in **Supplementary Table 2**.

Plant total nitrogen content at flowering (PTN0) is calculated as:

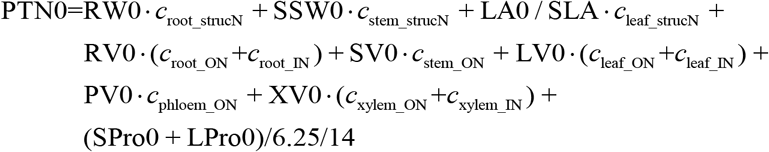

where, *c*_x_strucN_ are concentration of structural nitrogen in organ *x* (mol g^−1^). The factor 6.25 converts protein dry mass (g) to nitrogen dry mass (g) (Krul, 2019), and the factor 14 converts nitrogen dry mass (g) to nitrogen mole mass (mol; the same below). With default model parameter values, PTN0 amounts to 49 mmol per plant. For carbon budget from flowering to harvest, we have:

- the ratio between the post-anthesis respiration of the leaf and the post-anthesis photosynthesis = 504/2328*100% = 21.6%;
- the ratio between the post-anthesis respiration of the entire plant and the post anthesis photosynthesis = [420+504+84+204]/2328*100% = 52.1%;
- the “apparent contribution” of pre-anthesis stem non-structural carbohydrate storage to grain yield, i.e., the ratio between the loss of stem dry weight during grain filling and the grain yield at harvest = [240+84]/[1608-420]*100% = 27.3%;
- the actual contribution of pre-anthesis stem non-structural carbohydrate storage to grain yield, i.e., the ratio between carbohydrates produced by remobilization of pre-anthesis stem storage and the sum of carbohydrates from remobilization and post-anthesis photosynthesis = 240/[1764+240]*100% = 12.0%;
- the actual contribution of post-anthesis photosynthetically fixed carbon to grain filling = 1764/[1764+240]*100% = 88.0%.

For nitrogen budget from flowering to harvest, we have:

- the ratio between post-anthesis root nitrogen uptake and whole growing season nitrogen uptake = 20/[20+49]*100% = 29.0%;
- the ratio between the amount of nitrogen stored in the grain at harvest and the total plant nitrogen uptake during the whole growing season = 43.3/[20+49]*100% = 62.8%.

#### Simulating the effects of various endogenous and external perturbations on rice grain filling and yield (Figure 2)

The model was further applied to study effects of various endogenous and external perturbations on rice grain filling and yield; and, the experimental data were collected from multiple sources (**Supplementary Datasets 2**). The detailed model parameterization procedures in different case studies are documented in **Supplementary methods.**

### Sensitivity analysis of model parameters (Figure 3a-d; Supplementary Figures 6-13)

During parameter sensitivity analysis, 26 model parameters controlling capacities of biochemical processes (*ν*_max_*_) and two parameters controlling phloem resistance (R__phloem_leaf_root_ and R__phloem_leaf_grain_) were perturbed within [0.1, 10] fold to their default values individually. Grain number per ear was set to 190 for each group to allow sufficient sink size. Grain yield and values of other seven agronomic traits were calculated for each simulation. The agronomic traits are whole grain-filling season total canopy photosynthesis, whole grain-filling season total root nitrogen uptake, grain filling duration, grain nitrogen concentration at harvest, average grain filling rate from flowering to harvest, dry matter harvest index and nitrogen harvest index. The grain filling duration (GFD) is determined as the time interval between the flowering day and the day when total grain dry weight (TGDW1) reaches 95% of its value at the end of the simulation. The average grain filling rate (GFR) is calculated as:

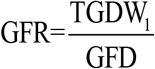

### Identification of optimal parameter combinations to maximize grain yield using a genetic algorithm (Figures 3e-r, 4; Supplementary Figures 14-16, 19, 20)

A genetic algorithm was used to identify the optimal value combinations of parameters to maximize grain yield. In brief, the algorithm mimics the process of natural biological evolution. The above mentioned 28 molecular biochemical and biophysical parameters, together with partitioning of initial mass among root weight, leaf area and grain number, were “perturbed” independently by randomly multiplying a scaling coefficient ranging from 0.5 to 2. Grain yield was used as the selection pressure in the algorithm.

To avoid a local optimum at the end of the evolution, five independent evolutionary populations were constructed with the following environmental variables: ambient CO_2_ concentration of 400 μbar; soil I-N concentration of 0.14 mol m^−3^; and grain-filling season of 50 days; for light intensity, air temperature and humidity, see **Supplementary Figure 21**. Each of the evolutionary populations included 100 generations with 100 individuals in each generation. As mentioned above, each parameter was restricted to vary between 0.5 to 2-fold of its default value. For the sake of comparison between lines, we restricted the total plant equivalent carbon and nitrogen at flowering to be the same for all individuals in the evolutionary populations. Specifically,

- Total plant equivalent carbon at flowering is a constant:

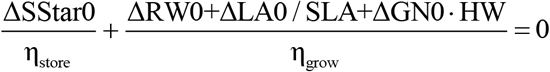

where parameters are defined in **Supplementary Table 2**.
- Total leaf nitrogen at flowering is a constant:

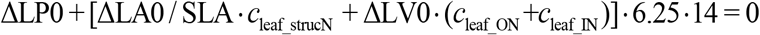

where, *c*_leaf_ON_ and *c*_leaf_IN_ are concentrations of leaf organic nitrogen and inorganic nitrogen, respectively; *c*leaf_strucN is nitrogen concentration in leaf structural matter (mol g^−1^).
- Total nitrogen in root, stem and grain at flowering is a constant:

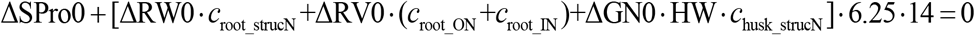

where, *c*_root_ON_ and *c*_root_IN_ are concentrations of root organic nitrogen and inorganic nitrogen, respectively; *c*_root_strucN_ and *c*_husk_strucN_ are nitrogen concentrations in root and husk structural matter (mol g^−1^), respectively.

### Field experiments

#### Plant biomass and grain yield of the wild type and *gif1* mutant (Figure 2o)

The *gif1* mutant was provided by courtesy of Dr. Zuhua He. The mutant is originated from the gamma radiation–induced mutant population of a *japonica* rice cultivar *Zhonghua 11* (E. Wang et al., 2008). The wild type *Zhonghua 11* and *gif1* mutant were planted in rows 20 cm apart and cultivated in standard paddy conditions at the experimental station in Songjiang district, Shanghai, China (latitude 30°56’44” N, 121°8’1” E). Plant biomass and grain yield of the wild type and *gif1* mutant were measured at harvest.

#### Grain filling and yield of rice cultivars grown in different ecozones (Figure 4; Supplementary Figure 17)

##### Shanghai, 2020

Four different rice cultivars, two *japonica* inbred rice cultivars *Nipponbare* (*NIP*) and *Xiu-Shui 134* (*XS134*), an *indica* inbred rice cultivar *9311*, and an *indica* hybrid rice cultivar *Y-Liang-You 900* (*Y900*), were grown with a planting space between hills of 0.20 × 0.20 m and cultivated under high and low nitrogen treatments at the experimental station in Songjiang district, Shanghai in 2020. For the high-nitrogen (HN) treatment, nitrogen fertilizer was applied at a rate of 152 kg ha^−1^; potassium (K_2_O) was applied at a rate of 180 kg ha^−1^. Phosphate (P_2_O_5_) was applied at a rate of 55 kg ha^−1^. For the low-nitrogen (LN) treatment, nitrogen, potassium and phosphate fertilizers were applied at a rate of 55 kg ha^−1^.

At flowering, 200 tillers with ears of similar length were labeled from four plots for each rice cultivar grown in each treatment. Fifteen labeled tillers were then randomly sampled, dried and weighed every 3-7 days from flowering to harvest. At harvest, 30 plants were randomly sampled for each rice cultivar grown in each treatment, and the dried straw biomass and ear biomass were measured separately. The dried rough grain yield is calculated as:

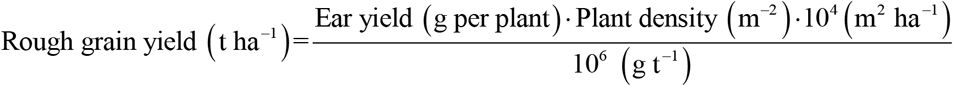

The dried brown grain yield is calculated as:

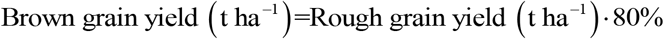

##### Hunan, 2020

Three high-yield *indica* hybrid rice cultivars, *Liang-You-Pei-Jiu* (*LYP9*), *Y900*, and *Chao You 1000* (*C1000*), were grown with a planting space between hills of 0.20 × 0.30 m and cultivated in standard paddy conditions at the experimental station in Longhui county, Hunan province, China (latitude 27°32’30” N, 110 °56’27” E) in 2020. Nitrogen fertilizer was applied at a rate of 300 kg ha^−1^; potassium (K_2_O) was applied at a rate of 150 kg ha^−1^; phosphate (P_2_O_5_) was applied at a rate of 300 kg ha^−1^. At flowering, 250 tillers with ears of similar length were labeled from five plots for each rice cultivar. Fifteen labeled tillers were then randomly sampled every 4 days from flowering to harvest. At harvest, 30 plants were randomly sampled and dried for each rice cultivar, and then the straw biomass and ear biomass were measured separately. The dried rough and brown grain yields are calculated as described above.

##### Yunnan, 2020

The same three high-yield *indica* hybrid rice cultivars were used as in Longhui, Hunan. Plants were grown with a planting space between hills of 0.20 × 0.30 m and cultivated in standard paddy conditions at the experimental station in Yongsheng county, Yunnan province, China (latitude 26°27’31” N, 100 °40’47” E) in 2020. Nitrogen fertilizer was applied at a rate of 330 kg ha^−1^; potassium (K_2_O) was applied at a rate of 165 kg ha^−1^; phosphate (P_2_O_5_) was applied at a rate of 330 kg ha^−1^. Other information and measurements were the same as that in Hunan, 2020.

#### Fitting grain filling pattern

To obtain mean grain filling rate during first *i* days after flowering, and instant grain filling rate on *i* day after flowering, we fitted dry weight gain with time of an ear (*W*) of rice cultivar *Nipponbare* using a beta growth function (Xinyou Yin, Goudriaan, Lantinga, Vos, & Spiertz, 2003):

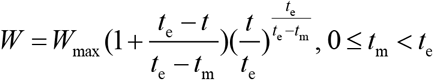

where, *W*_max_ is the final dry weight of an ear, which is reached at time *t*_e_; *t*_m_ is the time when maximum instant grain filling rate is reached.

For other rice cultivars with heavy ears with numerous spikelets, superior grains and inferior grains were filled asynchronously (J. Yang & Zhang, 2010; J. Yang et al., 2006);thus, we fitted dry weight gain with time of an ear (*W*) of these cultivars using a combination of two beta growth functions:

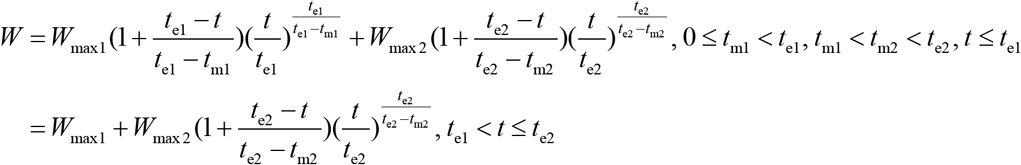

The mean grain filling rate during the first *i* days after flowering 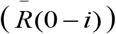 is defined as:

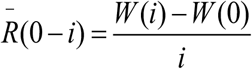

where, *W*(*i*) is dry weight of an ear on *i* th day after flowering. The instant grain filling rate on the *i^th^* day after flowering (*R*(*i*)) is defined as:

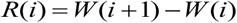

#### Grain respiratory rate during grain filling and the grain dry weight of an entire ear at harvest (Figure 4f; Supplementary Figure 18)

##### Shanghai, 2016

Seven rice cultivars were grown, in 2016, for experiments: *Y900, C1000, Shan-You 63* (*SY63*), *9311, XS134, Yong-You 538* (*YY538*) and *Yong-You 17* (*YY17*). Detailed experimental setup was described previously (Tian-Gen Chang et al., 2020). In brief, twenty tillers bearing ears with similar length were labeled at flowering for each cultivar. Respiratory rate of an entire ear was measured from flowering to harvest at a time interval of 4-15 days. Each measurement consisted five biological replicates. Five labeled ears were sampled, dried and weighed at harvest for each cultivar to determined the ear weight.

##### Shanghai, 2020

Rice cultivar *Nipponbare*, grown under low nitrogen and high nitrogen conditions as described above, was used. For each nitrogen condition, respiratory rates of five labeled ears were determined at 12, 17 and 21 days after flowering using the same protocol as described above.

#### Canopy photosynthesis, grain filling pattern and grain yield of rice cultivar XS134 (Figure 4g, h; Supplementary Figure 22)

##### Shanghai, 2015

Detailed field managements and measurements were described earlier (Tian-Gen Chang, Zhao, et al., 2019). In brief, plants of rice cultivar *XS134* were grown with a planting space between hills of 0.20 × 0.20 m and cultivated under low and high nitrogen conditions at the experimental station in Shanghai. Weather data, including photosynthetically active radiation (PAR), relative humidity and air temperature, were recorded every 10 minutes for the whole grain filling season (**Supplementary Datasets 3**).

At flowering, 100 tillers with ears of similar length were labeled for each treatment. Five labeled tillers were then randomly sampled every 5-6 days from flowering (September 12) to harvest (October 29). The samples were dried and the leaf biomass, stem biomass and ear biomass were measured separately.

Canopy-level gas exchange was measured with the Canopy Photosynthesis and Transpiration Measurement System (CAPTS), which comprises transparent chambers, sensors, and a control unit for data logging and storage (Q. Song, Xiao, Xiao, & Zhu, 2016). For each treatment, the canopy photosynthetic rates were recorded every 10 min for 3 independent plots at one day after flowering. At harvest, all plants covered by the chambers were harvested and threshed manually. The grains were then dried in an oven to determine the rough grain yield (kg m^−2^). The dried brown grain yield was then calculated as above described.

The measured diurnal net canopy photosynthetic rate (*A*_cn_) was fitted over incident solar photosynthetically active radiation (PAR) intensity (*I*) using the following equation:

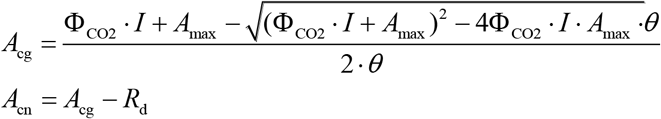

where, Φ_CO2_ is the maximum canopy apparent photosynthetic CO_2_ quantum yield, *A*_max_ is the maximum canopy photosynthetic rate, *θ* is an empirical coefficient (**Supplementary Figure 22**).

Daily total accumulated gross photosynthesis (*A*_cg tot_) was calculated based on PAR at different time (*t*) of a day:

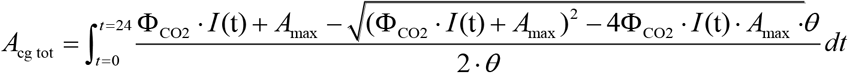

where, *I*(*t*) is incident solar PAR intensity at time *t*.

The potential photosynthate accumulation (*A*_c_fiowering→harvest_) is an integration of daily gross photosynthesis for all days (*D*) from flowering to harvest:

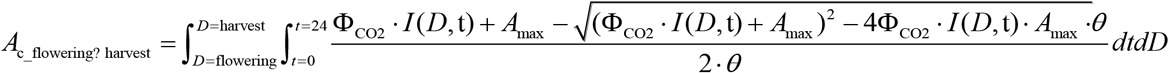

where, *I*(D, t) is incident solar PAR intensity at time *t* on day *D*.

## Supporting information

Datasets S1 paras_of_elites

Datasets S2 experimental_data_from_literature

Datasets S3 grain_filling_weather_for_2015

## Data and software availability

Experimental data extracted from literature, used in model-data comparison, are tabulated in **Supplementary Datasets 2**.

Source codes used for this study, together with the operation commands, are freely available for non-commercial use at https://github.com/rootchang/WACNI-rice.git.

## Acknowledgements

This research was financially supported by the open research fund of the state key laboratory of hybrid rice to T.C. (Hunan Hybrid Rice Research Center), the national natural science foundation of China to T.C. (Grant No: 32000285) and Q.S. (Grant No: 31970378), the Chinese Academy of Science strategic leading project to X.Z. (Grant No: XDB27020105) and the national key research and development project to T.C. (Grant No: 2018YFA0900600). The authors thank Zuhua He for providing the *gif1* mutant seeds; and Govindjee for constructive comments and improving the language of the manuscript; Youfa Liu and Xinyu Liu for assistance in experiments.

## Author contributions

X.Z. and T.C. designed the study; T.C. designed the model and the computational framework and analysed the data; Z.W., Z.S., T.C., H.Z., Y.X., S.C., M.Q., Q.S., F.C. and F.M. carried out the field experiments; T.C. and X.Z. drafted the manuscript; all authors provided critical feedback and helped shape the research, analysis and manuscript.

## Competing interests

The authors declare no competing financial interests.

## Notes

### Competing Interest Statement

The authors have declared no competing interest.

### Summary of Updates

The manuscript has been major revised. We have added a result sub-section (sub-section 4) to clarify the significance of grain filling rate to rice yield from both experimental perspective and theoretical perspective. The discussion section has been sharpened. The Materials and methods section has been expanded. Supplemental files updated with more experimental data provided.

## References

Ahmaruzzaman, M., & Gupta, V. K. (2011). Rice husk and its ash as low-cost adsorbents in water and wastewater treatment. Industrial & Engineering Chemistry Research, 50(24), 13589–13613. doi:10.1021/ie201477c

Ambavaram, M. M. R., Basu, S., Krishnan, A., Ramegowda, V., Batlang, U., Rahman, L., … Pereira, A. (2014). Coordinated regulation of photosynthesis in rice increases yield and tolerance to environmental stress. Nature Communications, 5, 5302. doi:10.1038/ncomms6302

Ashikari, M., Sakakibara, H., Lin, S., Yamamoto, T., Takashi, T., Nishimura, A., … Matsuoka, M. (2005). Cytokinin oxidase regulates rice grain production. Science, 309(5735), 741–745.

Bailey-Serres, J., Parker, J. E., Ainsworth, E. A., Oldroyd, G. E. D., & Schroeder, J. I. (2019). Genetic strategies for improving crop yields. Nature, 575(7781), 109–118. doi:10.1038/s41586-019-1679-0

Barillot, R., Chambon, C., & Andrieu, B. (2016). CN-Wheat, a functional–structural model of carbon and nitrogen metabolism in wheat culms after anthesis. I. Model description. Annals of Botany, 118(5), mcw144.

Braun, D. M., Wang, L., & Ruan, Y.-L. (2014). Understanding and manipulating sucrose phloem loading, unloading, metabolism, and signalling to enhance crop yield and food security. Journal of Experimental Botany, 65(7), 1713–1735. doi:10.1093/jxb/ert416

Chang, T.-G., Chang, S., Song, Q.-F., Perveen, S., & Zhu, X.-G. (2019). Systems models, phenomics and genomics: three pillars for developing high-yielding photosynthetically efficient crops. In Silico Plants, 1(1). doi:10.1093/insilicoplants/diy003

Chang, T.-G., Song, Q.-F., Zhao, H.-L., Chang, S., Xin, C., Qu, M., & Zhu, X.-G. (2020). An in situ approach to characterizing photosynthetic gas exchange of rice panicle. Plant Methods, 16(1), 92. doi:10.1186/s13007-020-00633-1

Chang, T.-G., Zhao, H., Wang, N., Song, Q.-F., Xiao, Y., Qu, M., & Zhu, X.-G. (2019). A three-dimensional canopy photosynthesis model in rice with a complete description of the canopy architecture, leaf physiology, and mechanical properties. Journal of Experimental Botany, 70(9), 2479–2490. doi:10.1093/jxb/ery430

Chang, T.-G., & Zhu, X.-G. (2017). Source-sink interaction: a century old concept under the light of modern molecular systems biology. Journal of Experimental Botany, 68(16), 4417–4431. doi:10.1093/jxb/erx002

Chen, J.-H., Chen, S.-T., He, N.-Y., Wang, Q.-L., Zhao, Y., Gao, W., & Guo, F.-Q. (2020). Nuclear-encoded synthesis of the D1 subunit of photosystem II increases photosynthetic efficiency and crop yield. Nature Plants, 6(5), 570–580. doi:10.1038/s41477-020-0629-z

Chen, J., Wang, Z., Chen, G., Wang, Y.-x., & Mo, Y.-w. (2006). Effects of various nitrogen fertilizer treatments on filling and respiratory rate of rice caryopsis. Chinese Journal of Rice Science, 20(4), 396–400.

Chen, Z., Wang, H.-G., Wen, Z.-J., & Wang, Y. (2007). Life sciences and biotechnology in China. Philosophical Transactions of the Royal Society of London, Series B: Biological Sciences, 362(1482), 947–957. doi:doi:10.1098/rstb.2007.2025

Chenu, K., Porter, J. R., Martre, P., Basso, B., Chapman, S. C., Ewert, F., … Asseng, S. (2017). Contribution of crop models to adaptation in wheat. Trends in Plant Science, 22(6), 472–490. doi:10.1016/j.tplants.2017.02.003

Cheung, F. (2014). Yield: The search for the rice of the future. Nature, 514(7524), S60–S61.

Chew, Y. H., Wenden, B., Flis, A., Mengin, V., Taylor, J., Davey, C. L., … De Reffye, P. (2014). Multiscale digital Arabidopsis predicts individual organ and whole-organism growth. Proceedings of the National academy of Sciences of the United States of America, 111(39), E4127–E4136.

Deng, J., Ran, J., Wang, Z., Fan, Z., Wang, G., Ji, M., … Brown, J. H. (2012). Models and tests of optimal density and maximal yield for crop plants. Proceedings of the National academy of Sciences of the United States of America, 109(39), 15823–15828. doi:10.1073/pnas.1210955109

Dingkuhn, M., Schnier, H., De Datta, S., Dorffling, K., & Javellana, C. (1991). Relationships between ripening-phase productivity and crop duration, canopy photosynthesis and senescence in transplanted and direct-seeded lowland rice. Field Crops Research, 26(3-4), 327–345.

Donald, C. M. (1968). The breeding of crop ideotypes. Euphytica, 17(3), 385–403.

Fahy, B., Siddiqui, H., David, L. C., Powers, S. J., Borrill, P., Uauy, C., & Smith, A. M. (2018). Final grain weight is not limited by the activity of key starch-synthesising enzymes during grain filling in wheat. Journal of Experimental Botany, 69(22), 5461–5475. doi:10.1093/jxb/ery314

Fernie, A. R., Bachem, C. W. B., Helariutta, Y., Neuhaus, H. E., Prat, S., Ruan, Y.-L., … Sonnewald, U. (2020). Synchronization of developmental, molecular and metabolic aspects of source-sink interactions. Nature Plants, 6(2), 55–66. doi:10.1038/s41477-020-0590-x

Fujita, D., Trijatmiko, K. R., Tagle, A. G., Sapasap, M. V., Koide, Y., Sasaki, K., … Kobayashi, N. (2013). *NAL1* allele from a rice landrace greatly increases yield in modern *indica* cultivars. Proceedings of the National academy of Sciences of the United States of America, 110(51), 20431–20436. doi:10.1073/pnas.1310790110

Garcia-Molina, A., & Leister, D. (2020). Accelerated relaxation of photoprotection impairs biomass accumulation in Arabidopsis. Nature Plants, 6(1), 9–12. doi:10.1038/s41477-019-0572-z

Grafahrend-Belau, E., Junker, A., Eschenröder, A., Müller, J., Schreiber, F., & Junker, B. H. (2013). Multiscale metabolic modeling: dynamic flux balance analysis on a whole-plant scale. Plant Physiology, 163(2), 637–647.

Hammer, G., Messina, C., Wu, A., & Cooper, M. (2019). Biological reality and parsimony in crop models—why we need both in crop improvement! In Silico Plants, 1(1). doi:10.1093/insilicoplants/diz010

Hannah, L. C., Futch, B., Bing, J., Shaw, J. R., Boehlein, S., Stewart, J. D., … Greene, T. (2012). A shrunken-2 transgene increases maize yield by acting in maternal tissues to increase the frequency of seed development. The Plant Cell, 24(6), 2352–2363. doi:10.1105/tpc.112.100602

Haque, M. M., Pramanik, H. R., Biswas, J. K., Iftekharuddaula, K., & Hasanuzzaman, M. (2015). Comparative performance of hybrid and elite inbred rice varieties with respect to their source-sink relationship. The Scientific World Journal, 2015.

Huang, M., Jiang, P., Shan, S., Gao, W., Ma, G., Zou, Y., … Yuan, L. (2017). Higher yields of hybrid rice do not depend on nitrogen fertilization under moderate to high soil fertility conditions. Rice, 10(1), 43. doi:10.1186/s12284-017-0182-1

Huang, X., Wei, X., Sang, T., Zhao, Q., Feng, Q., Zhao, Y., … Zhang, Z. (2010). Genome-wide association studies of 14 agronomic traits in rice landraces. Nature Genetics, 42(11), 961–967.

Ida, M., Ohsugi, R., Sasaki, H., Aoki, N., & Yamagishi, T. (2009). Contribution of nitrogen absorbed during ripening period to grain filling in a high-yielding rice variety, Takanari. Plant Production Science, 12(2), 176–184. doi:10.1626/pps.12.176

Jones, D., Peterson, M., & Geng, S. (1979). Association between grain filling rate and duration and yield components in rice. Crop Science, 19(5), 641–644.

Kar, G., & Kumar, A. (2014). Forecasting rainfed rice yield with biomass of early phenophases, peak intercepted PAR and ground based remotely sensed vegetation indices. Journal of Agrometeorology, 16, 94–103.

Kato, T. (2004). Effect of spikelet removal on the grain filling of Akenohoshi, a rice cultivar with numerous spikelets in a panicle. Journal of Agricultural Science, 142(2), 177–181.

Khan, N., Essemine, J., Hamdani, S., Qu, M., Lyu, M.-J. A., Perveen, S., … Zhu, X.-G. (2020). Natural variation in the fast phase of chlorophyll a fluorescence induction curve (OJIP) in a global rice minicore panel. Photosynthesis Research. doi:10.1007/s11120-020-00794-z

Kim, H., Lieffering, M., Miura, S., Kobayashi, K., & Okada, M. (2001). Growth and nitrogen uptake of CO_2_ - enriched rice under field conditions. New Phytologist, 150(2), 223–229.

Kobata, T., Sugawara, M., & Takatu, S. (2000). Shading during the early grain filling period does not affect potential grain dry matter increase in rice. Agronomy Journal, 92(3), 411–417.

Krul, E. S. (2019). Calculation of nitrogen-to-protein conversion factors: a review with a focus on soy protein. Journal of the American Oil Chemists’ Society, 96(4), 339–364. doi:https://doi.org/10.1002/aocs.12196

Laza, M. R., Peng, S., Akita, S., & Saka, H. (2003). Contribution of biomass partitioning and translocation to grain yield under sub-optimum growing conditions in irrigated rice. Plant Production Science, 6(1), 28–35. doi:10.1626/pps.6.28

Lee, S., Marmagne, A., Park, J., Fabien, C., Yim, Y., Kim, S.-j., … Nam, H. G. (2020). Concurrent activation of *OsAMT1;2* and *OsGOGAT1* in rice leads to enhanced nitrogen use efficiency under nitrogen limitation. The Plant Journal, 103(1), 7–20. doi:https://doi.org/10.1111/tpj.14794

Li, X., Qian, Q., Fu, Z., Wang, Y., Xiong, G., Zeng, D., … Li, J. (2003). Control of tillering in rice. Nature, 422(6932), 618–621. doi:10.1038/nature01518

Li, X., Wang, P., Li, J., Wei, S., Yan, Y., Yang, J., … Zhou, W. (2020). Maize *GOLDEN2-LIKE* genes enhance biomass and grain yields in rice by improving photosynthesis and reducing photoinhibition. Communications Biology, 3(1), 151–151. doi:10.1038/s42003-020-0887-3

Li, Y., Xiao, J., Chen, L., Huang, X., Cheng, Z., Han, B., … Wu, C. (2018). Rice functional genomics research: past decade and future. Molecular Plant, 11(3), 359–380. doi:https://doi.org/10.1016/?.molp.2018.01.007

Li, Z., Pinson, S. R. M., Stansel, J. W., & Paterson, A. H. (1998). Genetic dissection of the source-sink relationship affecting fecundity and yield in rice (shape *Oryza sativa* L.). Molecular Breeding, 4(5), 419–426.

Liang, C., Wang, Y., Zhu, Y., Tang, J., Hu, B., Liu, L., … Chu, J. (2014). OsNAP connects abscisic acid and leaf senescence by fine-tuning abscisic acid biosynthesis and directly targeting senescence-associated genes in rice. Proceedings of the National academy of Sciences of the United States of America, 111(27), 10013–10018.

Liu, Y., Zhu, X., He, X., Li, C., Chang, T., Chang, S., … Zhang, Y. (2020). Scheduling of nitrogen fertilizer topdressing during panicle differentiation to improve grain yield of rice with a long growth duration. Scientific Reports, 10(1), 15197. doi:10.1038/s41598-020-71983-y

Long, S. P., Marshall-Colon, A., & Zhu, X.-G. (2015). Meeting the global food demand of the future by engineering crop photosynthesis and yield potential. Cell, 161(1), 56–66.

Ma, S.-C., Li, F.-M., Xu, B.-C., & Huang, Z.-B. (2010). Effect of lowering the root/shoot ratio by pruning roots on water use efficiency and grain yield of winter wheat. Field Crops Research, 115(2), 158–164.

Makino, A., Kaneta, Y., Obara, M., Ishiyama, K., Kanno, K., Kondo, E., … Mae, T. (2020). High yielding ability of a large-grain rice cultivar, Akita 63. Scientific Reports, 10(1), 12231. doi:10.1038/s41598-020-69289-0

Marshall-Colon, A., Long, S. P., Allen, D. K., Allen, G., Beard, D. A., Benes, B., … Hart, J. C. (2017). Crops *in silico*: generating virtual crops using an integrative and multi-scale modeling platform. Frontiers in Plant Science, 8(786).

Nada, R. M., & Abogadallah, G. M. (2016). Restricting the above ground sink corrects the root/shoot ratio and substantially boosts the yield potential per panicle in field - grown rice *(Oryza sativa*L.). Physiologia Plantarum, 156(4), 371–386. doi:https://doi.org/10.1111/ppl.12377

Nadakuduti, S. S., & Enciso-Rodríguez, F. (2021). Advances in genome editing with CRISPR systems and transformation technologies for plant DNA manipulation. Frontiers in Plant Science, 11(2267). doi:10.3389/fpls.2020.637159

Ni, J., Li, J., Zhu, R., Zhang, M., Qi, K., Zhang, S., & Wu, J. (2020). Overexpression of sugar transporter gene *PbSWEET4* of pear causes sugar reduce and early senescence in leaves. Gene, 743, 144582. doi:10.1016/?.gene.2020.144582

Okamura, M., Arai-Sanoh, Y., Yoshida, H., Mukouyama, T., Adachi, S., Yabe, S., … Kondo, M. (2018). Characterization of high-yielding rice cultivars with different grain-filling properties to clarify limiting factors for improving grain yield. Field Crops Research, 219, 139–147. doi:https://doi.org/10.1016/j.fcr.2018.01.035

Paul, M. J., Watson, A., & Griffiths, C. A. (2019). Linking fundamental science to crop improvement through understanding source and sink traits and their integration for yield enhancement. Journal of Experimental Botany, 71(7), 2270–2280. doi:10.1093/jxb/erz480

Peleman, J. D., & van der Voort, J. R. (2003). Breeding by design. Trends in Plant Science, 8(7), 330–334. doi:10.1016/S1360-1385(03)00134-1

Peng, B., Guan, K., Tang, J., Ainsworth, E. A., & Zhou, W. (2020). Towards a multiscale crop modelling framework for climate change adaptation assessment. Nature Plants, 6, 338–348.

Peng, S., Cassman, K. G., Virmani, S., Sheehy, J., & Khush, G. (1999). Yield potential trends of tropical rice since the release of IR8 and the challenge of increasing rice yield potential. Crop Science, 39, 1552–1559. doi:10.2135/cropsci1999.3961552x

Peng, S., Khush, G. S., Virk, P., Tang, Q., & Zou, Y. (2008). Progress in ideotype breeding to increase rice yield potential. Field Crops Research, 108(1), 32–38.

Perveen, S., Qu, M., Chen, F., Essemine, J., Khan, N., Lyu, M.-J. A., … Zhu, X.-G. (2020). Overexpression of maize transcription factor mEmBP-1 increases photosynthesis, biomass, and yield in rice. Journal of Experimental Botany, 71(16), 4944–4957. doi:10.1093/jxb/eraa248

Postma, J. A., Kuppe, C., Owen, M. R., Mellor, N., Griffiths, M., Bennett, M. J., … Watt, M. (2017). OpenSimRoot: widening the scope and application of root architectural models. New Phytologist, 215(3), 1274–1286.

Qian, Q., Guo, L. B., Smith, S. M., & Li, J. Y. (2016). Breeding high-yield superior quality hybrid super rice by rational design. National Science Review, 3(3), 283–294. doi:10.1093/nsr/nww006

Ray, D. K., Mueller, N. D., West, P. C., & Foley, J. A. (2013). Yield trends are insufficient to double global crop production by 2050. PLoS One, 8(6), e66428.

Rossi, M., Bermudez, L., & Carrari, F. (2015). Crop yield: challenges from a metabolic perspective. Current Opinion in Plant Biology, 25, 79–89. doi:https://doi.org/10.1016/j.pbi.2015.05.004

Rötter, R. P., Tao, F., Höhn, J. G., & Palosuo, T. (2015). Use of crop simulation modelling to aid ideotype design of future cereal cultivars. Journal of Experimental Botany, 66(12), 3463–3476.

Saiz-Rubio, V., & Rovira-Más, F. (2020). From smart farming towards agriculture 5.0: a review on crop data management. Agronomy, 10(2). doi:10.3390/agronomy10020207

Salon, C., Avice, J.-C., Colombié, S., Dieuaide-Noubhani, M., Gallardo, K., Jeudy, C., … Rolin, D. (2017). Fluxomics links cellular functional analyses to whole-plant phenotyping. Journal of Experimental Botany, 68(9), 2083–2098. doi:10.1093/jxb/erx126

Saripalli, G., & Gupta, P. K. (2015). AGPase: its role in crop productivity with emphasis on heat tolerance in cereals. Theoretical and Applied Genetics, 128(10), 1893–1916. doi:10.1007/s00122-015-2565-2

Seki, M., Feugier, F. G., Song, X.-J., Ashikari, M., Nakamura, H., Ishiyama, K., … Satake, A. (2014). A mathematical model of phloem sucrose transport as a new tool for designing rice panicle structure for high grain yield. Plant and Cell Physiology, 56(4), 605–619. doi:10.1093/pcp/pcu191

Sheehy, J. E., Dionora, M. J. A., & Mitchell, P. L. (2001). Spikelet numbers, sink size and potential yield in rice. Field Crops Research, 71(2), 77–85. doi:https://doi.org/10.1016/S0378-4290(01)00145-9

Shen, B.-R., Wang, L.-M., Lin, X.-L., Yao, Z., Xu, H.-W., Zhu, C.-H., … Peng, X.-X. (2019). Engineering a new chloroplastic photorespiratory bypass to increase photosynthetic efficiency and productivity in rice. Molecular Plant. doi:https://doi.org/10.1016/j.molp.2018.11.013

Shimono, H., & Bunce, J. A. (2009). Acclimation of nitrogen uptake capacity of rice to elevated atmospheric CO_2_ concentration. Annals of Botany, 103(1), 87–94. doi:10.1093/aob/mcn209

Shin, D., Lee, S., Kim, T.-H., Lee, J.-H., Park, J., Lee, J., … Nam, H. G. (2020). Natural variations at the Stay-Green gene promoter control lifespan and yield in rice cultivars. Nature Communications, 11(1), 2819–2819. doi:10.1038/s41467-020-16573-2

Smidansky, E. D., Clancy, M., Meyer, F. D., Lanning, S. P., Blake, N. K., Talbert, L. E., & Giroux, M. J. (2002). Enhanced ADP-glucose pyrophosphorylase activity in wheat endosperm increases seed yield. Proceedings of the National academy of Sciences of the United States of America, 99(3), 1724–1729.

Smidansky, E. D., Martin, J. M., Hannah, L. C., Fischer, A. M., & Giroux, M. J. (2003). Seed yield and plant biomass increases in rice are conferred by deregulation of endosperm ADP-glucose pyrophosphorylase. Planta, 216(4), 656–664. doi:10.1007/s00425-002-0897-z

Soda, N., Gupta, B. K., Anwar, K., Sharan, A., Govindjee Singla-Pareek, S. L., & Pareek, A. (2018). Rice intermediate filament, OsIF, stabilizes photosynthetic machinery and yield under salinity and heat stress. Scientific Reports, 8(1), 4072–4072. doi:10.1038/s41598-018-22131-0

Song, Q., Srinivasan, V., Long, S. P., & Zhu, X.-G. (2019). Decomposition analysis on soybean productivity increase under elevated CO_2_ using 3-D canopy model reveals synergestic effects of CO_2_ and light in photosynthesis. Annals of Botany. doi:10.1093/aob/mcz163

Song, Q., Xiao, H., Xiao, X., & Zhu, X.-G. (2016). A new canopy photosynthesis and transpiration measurement system (CAPTS) for canopy gas exchange research. Agricultural and Forest Meteorology, 217, 101–107.

Song, Q. F., Zhang, G. L., & Zhu, X. G. (2013). Optimal crop canopy architecture to maximise canopy photosynthetic CO_2_ uptake under elevated CO_2_ - a theoretical study using a mechanistic model of canopy photosynthesis. Functional Plant Biology, 40(2), 109–124. doi:10.1071/FP12056

Sonnewald, U., Fernie, A. R., Gruissem, W., Schläpfer, P., Anjanappa, R. B., Chang, S.-H., … Zierer, W. (2020). The Cassava Source–Sink project: opportunities and challenges for crop improvement by metabolic engineering. The Plant Journal, n/a(n/a). doi:10.1111/tpj.14865

Tang, W.-B., Deng, H.-B., Xiao, Y.-H., Zhang, G.-L., Fan, K., Mo, H., & Chen, L.-Y. (2010). Root characteristics of high-yield C Liangyou rice combinations of two-line hybrid rice. Scientia Agricultura Sinica, 43(14), 2859–2868. doi:10.3864/j.issn.0578-1752.2010.14.004

Tcherkez, G. G. B., & Ribas-Carbó, M. (2012). Interactions between photosynthesis and day respiration. In F. Loreto, H. Medrano, & J. Flexas (Eds.), Terrestrial Photosynthesis in a Changing Environment: A Molecular, Physiological, and Ecological Approach (pp. 41–53). Cambridge: Cambridge University Press.

Thomas, H., & Howarth, C. J. (2000). Five ways to stay green. Journal of Experimental Botany, 51(suppl_1), 329–337. doi:10.1093/jexbot/51.suppl_1.329

Thomas, H., & Ougham, H. (2014). The stay-green trait. Journal of Experimental Botany, 65(14), 3889–3900. doi:10.1093/jxb/eru037

Tuberosa, R. (2012). Phenotyping for drought tolerance of crops in the genomics era. Frontiers in Physiology, 3, 347–347. doi:10.3389/fphys.2012.00347

Uga, Y. (2021). Challenges to design-oriented breeding of root system architecture adapted to climate change. Breeding science, 71(1), 3–12. doi:10.1270/jsbbs.20118

Van Der Straeten, D., Bhullar, N. K., De Steur, H., Gruissem, W., MacKenzie, D., Pfeiffer, W., … Bouis, H. (2020). Multiplying the efficiency and impact of biofortification through metabolic engineering. Nature Communications, 11(1), 5203. doi:10.1038/s41467-020-19020-4

Wang, E., Wang, J., Zhu, X., Hao, W., Wang, L., Li, Q., … Lin, H. (2008). Control of rice grain-filling and yield by a gene with a potential signature of domestication. Nature Genetics, 40(11), 1370–1374.

Wang, Q., Nian, J., Xie, X., Yu, H., Zhang, J., Bai, J., … Chen, L. (2018). Genetic variations in *ARE1* mediate grain yield by modulating nitrogen utilization in rice. Nature Communications, 9(1).

Wang, Y., & Li, J. (2008). Molecular basis of plant architecture. Annual Review of Plant Biology, 59, 253–279.

Wang, Y., Noguchi, K., Ono, N., Inoue, S., Terashima, I., & Kinoshita, T. (2014). Overexpression of plasma membrane H^+^-ATPase in guard cells promotes light-induced stomatal opening and enhances plant growth. Proceedings of the National academy of Sciences of the United States of America, 111(1), 533–538.

Wang, Y., Song, Q. F., Jaiswal, D., de Souza, A. P., Long, S. P., & Zhu, X. G. (2017). Development of a three-dimensional ray-tracing model of sugarcane canopy photosynthesis and its application in assessing impacts of varied row spacing. Bioenergy Research, 10(3), 626–634. doi:10.1007/s12155-017-9823-x

Watanabe, T., Hanan, J. S., Room, P. M., Hasegawa, T., Nakagawa, H., & Takahashi, W. (2005). Rice morphogenesis and plant architecture: Measurement, specification and the reconstruction of structural development by 3D architectural modelling. Annals of Botany, 95(7), 1131–1143. doi:Doi 10.1093/Aob/Mci136

Wei, H., Meng, T., Li, X., Dai, Q., Zhang, H., & Yin, X. (2018). Sink-source relationship during rice grain filling is associated with grain nitrogen concentration. Field Crops Research, 215, 23–38. doi:https://doi.org/10.1016/j.fcr.2017.09.029

White, A. C., Rogers, A., Rees, M., & Osborne, C. P. (2015). How can we make plants grow faster? A source-sink perspective on growth rate. Journal of Experimental Botany, 67(1), 31–45. doi:10.1093/jxb/erv447

Whitfield, D. M., Connor, D. J., & Hall, A. J. (1989). Carbon dioxide balance of sunflower *(Helianthus annuus)* subjected to water stress during grain-filling. Field Crops Research, 20(1), 65–80. doi:https://doi.org/10.1016/0378-4290(89)90024-5

Witt, C., Dobermann, A., Abdulrachman, S., Gines, H. C., Guanghuo, W., Nagarajan, R., … Olk, D. C. (1999). Internal nutrient efficiencies of irrigated lowland rice in tropical and subtropical Asia. Field Crops Research, 63(2), 113–138. doi:10.1016/S0378-4290(99)00031-3

Xing, Y., & Zhang, Q. (2010). Genetic and molecular bases of rice yield. Annual Review of Plant Biology, 61, 421–442.

Yang, J., & Zhang, J. (2010). Grain-filling problem in ‘super’ rice. Journal of Experimental Botany, 61(1), 1–5.

Yang, J., Zhang, J., Wang, Z., Liu, K., & Wang, P. (2006). Post-anthesis development of inferior and superior spikelets in rice in relation to abscisic acid and ethylene. Journal of Experimental Botany, 57(1), 149–160.

Yang, J., Zhang, J., Wang, Z., Zhu, Q., & Wang, W. (2001). Remobilization of carbon reserves in response to water deficit during grain filling of rice. Field Crops Research, 71(1), 47–55.

Yang, W., Peng, S., Dionisio-Sese, M. L., Laza, R. C., & Visperas, R. M. (2008). Grain filling duration, a crucial determinant of genotypic variation of grain yield in field-grown tropical irrigated rice. Field Crops Research, 105(3), 221–227.

Yin, X., Goudriaan, J., Lantinga, E. A., Vos, J., & Spiertz, H. J. (2003). A flexible sigmoid function of determinate growth. Annals of Botany, 91(3), 361–371.

Yin, X., & van Laar, H. (2005). Crop Systems Dynamics: An Ecophysiological Model of Genotype-by-Environment Interactions (GECROS). Wageningen Academic Pub., Wageningen.

Yoon, D.-K., Ishiyama, K., Suganami, M., Tazoe, Y., Watanabe, M., Imaruoka, S., … Makino, A. (2020). Transgenic rice overproducing Rubisco exhibits increased yields with improved nitrogen-use efficiency in an experimental paddy field. Nature Food, 1(2), 134–139. doi:10.1038/s43016-020-0033-x

Yoshida, S. (1981). Fundamentals of rice crop science: Los Banos, Philippines: International Rice Research Institute.

Yoshinaga, S., Takai, T., Arai-Sanoh, Y., Ishimaru, T., & Kondo, M. (2013). Varietal differences in sink production and grain-filling ability in recently developed high-yielding rice (*Oryza sativa*L.) varieties in Japan. Field Crops Research, 150(15), 74–82.

Yu, H., Lin, T., Meng, X., Du, H., Zhang, J., Liu, G., … Li, J. (2021). A route to *de novo* domestication of wild allotetraploid rice. Cell, 184(5), 1156–1170.e1114. doi:https://doi.org/10.1016/j.cell.2021.01.013

Yu, J., Zhen, X., Li, X., Li, N., & Xu, F. (2019). Increased autophagy of rice can increase yield and nitrogen use efficiency (NUE). Frontiers in Plant Science, 10(584). doi:10.3389/fpls.2019.00584

Yu, S.-M., Lo, S.-F., & Ho, T.-H. D. (2015). Source–sink communication: regulated by hormone, nutrient, and stress cross-signaling. Trends in Plant Science, 20(12), 844–857. doi:10.1016/j.tplants.2015.10.009

Yuan, L. (2017). Progress in super-hybrid rice breeding. The Crop Journal, 5(2), 100–102.

Zhang, C.-C., Zhou, C.-Z., Burnap, R. L., & Peng, L. (2018). Carbon/nitrogen metabolic balance: lessons from cyanobacteria. Trends in Plant Science, 23(12), 1116–1130. doi:https://doi.org/10.1016/j.tplants.2018.09.008

Zhao, Q., Huang, P., & Ling, Q. (2001). Relations between canopy apparent photosynthesis and store matter in stem and sheath between and yield and nitrogen regulations in rice. Scientia Agricultura Sinica, 34(3), 304–310.

Zhao, Y., Xi, M., Zhang, X., Lin, Z., Ding, C., Tang, S., … Ding, Y. (2015). Nitrogen effect on amino acid composition in leaf and grain of japonica rice during grain filling stage. Journal of Cereal Science, 64, 29–33.

Zhou, Y., Liu, L., Huang, W., Yuan, M., Zhou, F., Li, X., & Lin, Y. (2014). Overexpression of *OsSWEET5* in rice causes growth retardation and precocious senescence. PLoS One, 9(4), e94210–e94210. doi:10.1371/journal.pone.0094210

