## supplementary materials for "A critical role of stable grain filling rate in maximizing rice yield revealed by whole plant carbon nitrogen interaction modeling"

**This PDF file includes:**

Supplementary methods
Supplementary Figures 1 to 22
Supplementary Tables 1 to 3
References

**Other Supplementary Materials for this manuscript include the following:**

Supplementary Datasets 1 (Excel)
Supplementary Datasets 2 (Excel)
Supplementary Datasets 3 (Excel)

### Supplementary methods

#### Model development

The WACNI model developed in this work kinetically simulated the rates of major basic biochemical and physiological processes in a plant using ordinary differential equations. Fourteen different types of primary biochemical/biophysical processes were incorporated in the model (**Figure 1**). The sub-models used or developed for these processes were described in four different subsections according to the organs they present in, i.e., root, leaf, grains and stem (including culm and sheath) (Eqn 1.1-14.1). The respiration for each organ was described in the subsequent *Respiration* subsection (Eqn 15.1-15.4). A summary of all the sub-models used and developed here was shown in **Supplementary Table 1**.

##### *Root*

*I-N uptake and assimilation*: Two I-N uptake systems present in roots, i.e., the high affinity transport system (HATS) and the low affinity transport system (LATS); which together ensure I-N uptake at different soil nitrogen (N) concentrations (Glass et al., 2002). The uptake rates of HATS and LATS were modeled with Michaelis-Menten kinetics model and linear model, respectively (Malagoli et al., 2004) (Eqn 1.2). Sugar level also influences mineral transport as sugar provides the energy needed for these processes (Henry & Raper Jr, 1991; Pitman & Cram, 2013). The impact of root sucrose level on I-N uptake was modelled with a Monod function (Monod, 1949), i.e., the first item of Eqn (1.2). Root total N concentration has a feedback inhibitory effect ($\alpha_{N\_inhibit}$) on root HATS (Glass et al., 2002). Finally, these regulatory processes were incorporated into a single root I-N uptake equation to calculate the total root I-N uptake rate ($v_{root\_N\_upt}$):

$$\alpha_{N\_inhibit} = \begin{cases} 1 - \dfrac{[N]_{root}}{[N]_{inhibit\_up}}, [N]_{root} < [N]_{inhibit\_up} \\ 0, [N]_{root} \geq [N]_{inhibit\_up} \end{cases} \quad (1.1)$$

$$v_{\text{root\_N\_upt}} = \frac{[\text{Suc}]_{\text{root}}}{K_{\text{m1}} + [\text{Suc}]_{\text{root}}} \cdot (\alpha_{\text{N\_inhibit}} \cdot v_{\text{m\_root\_HATS}} \cdot \frac{[\text{I-N}]_{\text{soil}}}{K_{\text{m2}} + [\text{I-N}]_{\text{soil}}} + k_{\text{root\_LATS}} \cdot [\text{I-N}]_{\text{soil}}) \quad (1.2)$$

where, $[\text{I-N}]_{\text{soil}}$ is soil I-N concentration, $[\text{Suc}]_{\text{root}}$ is root sucrose concentration, $[\text{N}]_{\text{root}}$ is total root N concentration (including I-N and O-N). $[\text{N}]_{\text{inhibit\_up}}$ is the upper limit of total root N concentration above which N uptake by HATS ceases, $K_{\text{m1}}$ and $K_{\text{m2}}$ are Michaelis-constants, $v_{\text{m\_root\_HATS}}$ is root HATS maximum I-N uptake rate, $k_{\text{root\_LATS}}$ is root LATS I-N uptake rate coefficient (**Supplementary Table 2**).

Root N assimilation depends on energy and reducing power derived from sugar (Miflin & Lea, 1980). Hence, root I-N assimilation rate ($v_{\text{root\_N\_ass}}$) was modelled based on $[\text{Suc}]_{\text{root}}$ and $[\text{I-N}]_{\text{root}}$:

$$v_{\text{root\_N\_ass}} = v_{\text{m\_root\_N\_ass}} \cdot \frac{[\text{Suc}]_{\text{root}}}{K_{\text{m3}} + [\text{Suc}]_{\text{root}}} \cdot \frac{[\text{I-N}]_{\text{root}}}{K_{\text{m4}} + [\text{I-N}]_{\text{root}}} \quad (2.1)$$

where, $K_{\text{m3}}$ and $K_{\text{m4}}$ are Michaelis-constants, $v_{\text{m\_root\_N\_ass}}$ is root maximum I-N assimilation rate (**Supplementary Table 2**).

*Root growth***:** As sugar supply is a major factor limiting root growth (Radin, Parker, & Sell, 1978), and organ growth rate is positively correlated with the sugar level in the organ before reaching its maximum (Muller et al., 2011; Radin et al., 1978; Willaume & Pagès, 2011), a critical sucrose concentration ($[\text{Suc}]_{\text{grow\_low}}$) was set, below which root growth ceased. A Monod function was applied to describe the relation between root sucrose level and its growth rate ($v_{\text{root\_grow}}$):

$$v_{\text{root\_grow}} = \begin{cases} 0, [\text{Suc}]_{\text{root}} < [\text{Suc}]_{\text{grow\_low}} \\ v_{\text{m\_root\_grow}} \cdot \dfrac{[\text{Suc}]_{\text{root}} - [\text{Suc}]_{\text{grow\_low}}}{K_{\text{m5}} + ([\text{Suc}]_{\text{root}} - [\text{Suc}]_{\text{grow\_low}})}, [\text{Suc}]_{\text{root}} \geq [\text{Suc}]_{\text{grow\_low}} \end{cases} \quad (3.1)$$

where, $K_{\text{m5}}$ is Michaelis-constant, $v_{\text{m\_root\_grow}}$ is the root maximum relative growth rate (**Supplementary Table 2**).

*Root senescence***:** Root loss during senescence was modelled based on two factors, aging and carbohydrate supply. Specifically, there is a minimal (constant) relative senescence rate $\alpha_{\text{sene\_root}}$ (**Supplementary Table 2**) due to the aging of root (Asseng,

Richter, & Wessolek, 1997; Johnson & Thornley, 1985), but when sucrose concentration is lower than a critical level ($[\text{Suc}]_{\text{sene\_up}}$), root senescence rate ($v_{\text{root\_sene}}$) would be accelerated as a result of carbohydrate starvation (Fanello, Bartoli, & Guiamet, 2017):

$$v_{\text{root\_sene}} = \begin{cases} \alpha_{\text{root\_sene}} \cdot e^{-\beta_{\text{root\_sene}} \cdot ([\text{Suc}]_{\text{root}} - [\text{Suc}]_{\text{sene\_up}})}, [\text{Suc}]_{\text{root}} < [\text{Suc}]_{\text{sene\_up}} \\ \alpha_{\text{root\_sene\_}}, [\text{Suc}]_{\text{root}} \geq [\text{Suc}]_{\text{sene\_up}} \end{cases} \quad (4.1)$$

where, $\beta_{\text{root\_sene}}$ is an empirical coefficient (**Supplementary Table 2**).

***Leaf***

*Photosynthesis and N assimilation*: Leaf level light reaction, Calvin-Benson cycle and photorespiration were modelled based on the Farquhar-von Caemmerer-Berry (FvB) model (Von Caemmerer, 2000b). To scale up them to a canopy level, a sun-shade model described in De Pury and Farquhar (1997) was used (see below).

Leaf protein level (especially the level of enzyme Rubisco, which accounts for major leaf protein) has been shown to be linearly and positively correlated with leaf light reaction and dark reaction activity before reaching their maximum in many experiments (San-oh, Sugiyama, Yoshita, Ookawa, & Hirasawa, 2006; Tazoe, Noguchi, & Terashima, 2006), while leaf non-structural carbohydrates (NSC, refer to sucrose and starch in the current model) accumulation can inhibit photosynthesis (Azcón-Bieto, 1983) through multiple feedback regulatory pathways (Paul & Pellny, 2003; Stitt, 1991). These effects were considered by setting an activation coefficient ($\alpha_{\text{Pro\_promote}}$, Eqn 5.1) of leaf protein content ($[\text{Pro}]_{\text{leaf}}$) and an inhibition coefficient ($\alpha_{\text{NSC\_inhibit}}$, Eqn 5.2) of leaf NSC content ($[\text{NSC}]_{\text{leaf}}$) to parameters involved in both photosynthetic electron transport rate $v_J$ and potential $CO_2$ assimilation rate $v_{A0}$ (Eqn 5.3-5.8):

$$\alpha_{\text{Pro\_promote}} = \begin{cases} \dfrac{[\text{Pro}]_{\text{leaf}}}{[\text{Pro}]_{\text{promote\_up}}}, [\text{Pro}]_{\text{leaf}} < [\text{Pro}]_{\text{promote\_up}} \\ 1, [\text{Pro}]_{\text{leaf}} \geq [\text{Pro}]_{\text{promote\_up}} \end{cases} \quad (5.1)$$

$$\alpha_{\text{NSC\_inhibit}} = \begin{cases} 1, [\text{NSC}]_{\text{leaf}} < [\text{NSC}]_{\text{inhibit\_low}} \\ 1 - \dfrac{[\text{NSC}]_{\text{leaf}} - [\text{NSC}]_{\text{inhibit\_low}}}{[\text{NSC}]_{\text{inhibit\_up}} - [\text{NSC}]_{\text{inhibit\_low}}}, [\text{NSC}]_{\text{inhibit\_low}} \leq [\text{NSC}]_{\text{leaf}} \leq [\text{NSC}]_{\text{inhibit\_up}} \\ 0, [\text{NSC}]_{\text{leaf}} > [\text{NSC}]_{\text{inhibit\_up}} \end{cases}$$

(5.2)

$$v_{\mathrm{cmax}} = \alpha_{\mathrm{Pro\_promote}} \cdot \alpha_{\mathrm{NSC\_inhibit}} \cdot v_{\mathrm{cmax0}} \quad (5.3)$$

$$J_{\mathrm{max}} = \alpha_{\mathrm{Pro\_promote}} \cdot \alpha_{\mathrm{NSC\_inhibit}} \cdot J_{\mathrm{max0}} \quad (5.4)$$

$$\theta = \alpha_{\mathrm{Pro\_promote}} \cdot \theta_0 \quad (5.5)$$

$$\varphi = \alpha_{\mathrm{Pro\_promote}} \cdot \varphi_0 \quad (5.6)$$

$$v_{\mathrm{A0}} = v_{\mathrm{cmax}} \cdot \frac{[\mathrm{CO_2}]_{\mathrm{i}} - \Gamma}{K_{\mathrm{m6}} + [\mathrm{CO_2}]_{\mathrm{i}}} \quad (5.7)$$

$$v_{\mathrm{J}} = \frac{\varphi \cdot I + J_{\mathrm{max}} - \sqrt{(\varphi \cdot I + J_{\mathrm{max}})^2 - 4 \cdot \varphi \cdot \theta \cdot I \cdot J_{\mathrm{max}}}}{2\theta} \quad (5.8)$$

where, $K_{\mathrm{m6}}$ is Michaelis-constant; $[\mathrm{Pro}]_{\mathrm{promote\_up}}$, $[\mathrm{NSC}]_{\mathrm{inhibit\_up}}$ and $[\mathrm{NSC}]_{\mathrm{inhibit\_low}}$ are critical leaf protein and NSC content; $\varphi_0$=0.85*0.5 and $\theta_0$=0.7 are empirical constants (Von Caemmerer, 2000b); $v_{\mathrm{cmax0}}$ is the maximum Rubisco carboxylation rate, $J_{\mathrm{max0}}$ is the maximum electron transport rate (**Supplementary Table 2**); Γ is $CO_2$ compensation point in the absence of mitochondrial respiration (**Supplementary Table 2**). $I$ is irradiance absorbed by leaves. Whole-plant $I$, $v_{\mathrm{cmax0}}$ and $J_{\mathrm{max0}}$ were calculated separately for sunlit and shaded leaves, e.g., $I_{\mathrm{sun}}$, $I_{\mathrm{sh}}$, $v_{\mathrm{cmax0\_sun}}$, $v_{\mathrm{cmax0\_sh}}$, $J_{\mathrm{max0\_sun}}$ and $J_{\mathrm{max0\_sh}}$ in Eqn 5.14-5.19, and $v_{\mathrm{\_leaf\_N\_ass0}}$ in Eqn 5.23, based on light extinction profile within the canopy throughout the day following De Pury and Farquhar (1997):

$$\delta = -23.4 \frac{\pi}{180} \cos \frac{2\pi(\mathrm{DOY}+10)}{365} \quad (5.9)$$

$$\sin\beta = \sin\lambda \cdot \sin\delta + \cos\lambda \cdot \cos\delta \cdot \cos\frac{\pi \cdot (\tau - 12)}{12} \quad (5.10)$$

$$f_d = \frac{1 - a^{1/\sin\beta}}{1 + a^{1/\sin\beta}(1/f_a - 1)} \quad (5.11)$$

$$L_{\mathrm{C\_sun}} = \frac{1 - e^{-k_b \cdot L_{\mathrm{C}}}}{k_b}(1 - e^{-k_b \cdot L_{\mathrm{C}}}) \quad (5.12)$$

$$I_{\mathrm{C}} = \int_0^{L_{\mathrm{C}}} I_l dl = (1-\rho_{cb}) \cdot I_b \cdot (1 - e^{-k_b' \cdot L_{\mathrm{C}}}) + (1-\rho_{cd}) \cdot I_d \cdot (1 - e^{-k_d' \cdot L_{\mathrm{C}}}) \quad (5.13)$$

$$I_{\mathrm{sun}} = \int_0^{L_{\mathrm{C}}} I_{l\_\mathrm{sun}} \cdot f_{l\_\mathrm{sun}} dl = I_b \cdot (1-\sigma) \cdot (1-e^{-k_b \cdot L_{\mathrm{C}}}) + I_d \cdot (1-\rho_{cd}) \cdot (1-e^{(-k_d^{'}+k_b)\cdot L_{\mathrm{C}}}) \cdot \frac{k_d^{'}}{k_d^{'}+k_b}$$
$$+ I_b \cdot (1-\rho_{cb}) \cdot (1-e^{(-k_b^{'}+k_b)\cdot L_{\mathrm{C}}} \cdot \frac{k_b^{'}}{k_b^{'}+k_b}) - (1-\sigma) \cdot \frac{1-e^{-2k_b \cdot L_{\mathrm{C}}}}{2} \qquad (5.14)$$

$$I_{\mathrm{sh}} = I_{\mathrm{C}} - I_{\mathrm{sun}} \qquad (5.15)$$

$$v_{\mathrm{cmax0\_sun}} = v_{\mathrm{cmax0}} \cdot L_{\mathrm{C\_sun}} \qquad (5.16)$$

$$J_{\mathrm{max0\_sun}} = J_{\mathrm{max0}} \cdot L_{\mathrm{C\_sun}} \qquad (5.17)$$

$$v_{\mathrm{cmax0\_sh}} = v_{\mathrm{cmax0}} \cdot (L_{\mathrm{C}} - L_{\mathrm{C\_sun}}) \qquad (5.18)$$

$$J_{\mathrm{max0\_sh}} = J_{\mathrm{max0}} \cdot (L_{\mathrm{C}} - L_{\mathrm{C\_sun}}) \qquad (5.19)$$

where, $\delta$ is solar declination angle; DOY is current day of year; $\beta$ is solar elevation angle; $\lambda$ is the latitude of experimental base; $\tau$ is the local time of day; $f_d$ is the fraction of diffuse irradiance; $f_a$=0.425 is the forward scattering coefficient of PAR in atmosphere; $a$=0.75 is the atmospheric transmission coefficient of PAR; $L_{\mathrm{c}}$ is canopy leaf area index; $L_{\mathrm{c\_sun}}$ is the sunlit leaf area index; $k_b=k_{b0}/\sin\beta$ is the beam radiation extinction coefficient of canopy; $k'_b=0.9k_b$ is beam and scattered beam PAR extinction coefficient; $k_d$=0.78 is diffuse PAR extinction coefficient; $k'_d=0.9k_d$ is diffuse and scattered diffuse PAR extinction coefficient; $I_0$ is the total incident PAR intensity at the top of the canopy; $I_d=I_0 \cdot f_d$ is diffuse PAR intensity; $I_b=I_0 \cdot (1-f_d)$ is beam PAR intensity; $I_{\mathrm{C}}$ is irradiance absorbed by the canopy; $I_{\mathrm{sun}}$ is irradiance absorbed by the sunlit leaves; $I_{\mathrm{sh}}$ is irradiance absorbed by the shaded leaves; $\rho_{\mathrm{cb}}$=0.029 is canopy reflection coefficient for beam PAR; $\rho_{\mathrm{cd}}$=0.036 is canopy reflection coefficient for diffuse PAR; $\sigma$=0.15 is leaf scattering coefficient of PAR. For further details, see De Pury and Farquhar (1997).

Intercellular $CO_2$ concentration ($[CO_2]_i$) is determined by ambient $CO_2$ concentration ($[CO_2]_a$), leaf stomatal conductance ($g_s$) and leaf net photosynthetic rate ($A - R_{\mathrm{leaf}}$):

$$[\mathrm{CO}_2]_{\mathrm{i}} = [\mathrm{CO}_2]_{\mathrm{a}} - \frac{A - R_{\mathrm{leaf}}}{g_{\mathrm{s}}} \qquad (5.20)$$

where, leaf gross photosynthetic rate $A$ is calculated, as in Eqn (5.27), and respiratory rate $R_{leaf}$ is calculated, as described in the following *Respiration* subsection.

Leaf stomatal conductance ($g_s$) is influenced by many factors, e.g., $CO_2$ concentration, vapor pressure deficit, light intensity, leaf water potential, temperature and abscisic acid concentration (Farquhar & Sharkey, 1982; Jarvis, 1976; J. Zhang & Davies, 1990). Diffusion of $CO_2$ from ambient air to intercellular space was modeled following Leuning (1995):

$$g_s = g_0 + a_1 \cdot \frac{A}{[CO_2]_i - \Gamma} \cdot \frac{1}{1 + \frac{VPD_{leaf}}{VPD_0}} \tag{5.21}$$

where, $g_0$=0.01 is residual stomatal conductance, $a_1$=20 and $VPD_0$=0.35 are empirical constants (Leuning, 1995). Leaf vapor pressure deficit ($VPD_{leaf}$) is the difference between leaf vapor pressure (which is assumed to be saturated and determined by leaf temperature $T$) and actual air vapor pressure (which is determined by relative air humidity $RH_{air}$), which was calculated following Allen, Pereira, Raes, and Smith (1998):

$$VPD_{leaf} = 0.6108 \exp[\frac{17.27T}{T + 237.3}] \cdot (1 - RH_{air}) \tag{5.22}$$

As photosynthetic light reactions generate abundant ATP and reducing power (e.g., NADPH and Ferredoxin), necessary for I-N assimilation, leaves, as well as roots, act as major N assimilators (Andrews, 1986). Potential I-N assimilation rate ($v_{N\_ass0}$) was modeled based on leaf protein level and I-N concentration ($[\text{I-N}]_{leaf}$):

$$v_{leaf\_N\_ass0} = \alpha_{Pro\_promote} \cdot v_{m\_leaf\_N\_ass} \cdot \frac{[\text{I-N}]_{leaf}}{K_{m7} + [\text{I-N}]_{leaf}} \tag{5.23}$$

where, $K_{m7}$ is Michaelis-constant, $v_{m\_leaf\_N\_ass}$ is leaf maximum N assimilation rate (**Supplementary Table 2**).

Calvin-Benson cycle, photorespiration and I-N assimilation compete for reducing power derived from the light reactions when light is limiting (Nunes-Nesi, Fernie, & Stitt, 2010). The rate of formation of the reducing power ($v_{NADPH\_p}$) was calculated as follows:

$$v_{NADPH\_p} = \frac{v_J}{2} \tag{5.24}$$

Calvin-Benson cycle and photorespiration consume 2 NADPH per cycle, and, on average, I-N assimilation consumes 3 NADPH per N (if $NO_3^-$ is the N source, it needs 5 NADPH or equivalent reduction power; if $NH_4^+$ is the N source, it needs 1 NADPH or equivalent reduction power) (Walker, Strand, Kramer, & Cousins, 2014). Thus the potential NADPH consumption rate ($v_{NADPH\_c0}$) was calculated as follows:

$$v_{\mathrm{NADPH\_c0}} = 2v_{\mathrm{cmax}} \cdot \frac{[\mathrm{CO_2}]_\mathrm{i} + \Gamma}{K_{\mathrm{m6}} + [\mathrm{CO_2}]_\mathrm{i}} + 3v_{\mathrm{leaf\_N\_ass0}} \quad (5.25)$$

The actual $CO_2$ assimilation rate $A$ as well as I-N assimilation rate ($v_{leaf\_N\_ass}$) were finally determined by balancing NADPH production and consumption:

$$\delta = \begin{cases} \frac{v_{\mathrm{NADPH\_p}}}{v_{\mathrm{NADPH\_c0}}}, v_{\mathrm{NADPH\_p}} < v_{\mathrm{NADPH\_c0}} \\ 1, v_{\mathrm{NADPH\_p}} \geq v_{\mathrm{NADPH\_c0}} \end{cases} \quad (5.26)$$

$$A = \delta \cdot v_{\mathrm{A0}} \quad (5.27)$$

$$v_{\mathrm{leaf\_N\_ass}} = \delta \cdot v_{\mathrm{leaf\_N\_ass0}} \quad (5.28)$$

Final $CO_2$ and I-N assimilation rates were determined by solving the above equations (Eqn 5.1-5.28).

*Leaf sucrose synthesis and starch synthesis/degradation*: During the day, photosynthesis- derived triose phosphate (TP) convert to sucrose ($v_{leaf\_Suc\_syn}$) and starch ($v_{leaf\_Star\_syn}$) synchronously (Zhu, de Sturler, & Long, 2007):

$$v_{\mathrm{leaf\_Suc\_syn}} = v_{\mathrm{m\_leaf\_Suc\_syn}} \cdot \frac{[\mathrm{TP}]_{\mathrm{leaf}} - \frac{[\mathrm{Suc}]_{\mathrm{leaf}}}{K_{\mathrm{e1}}}}{K_{\mathrm{m8}} + [\mathrm{TP}]_{\mathrm{leaf}}} \quad (6.1)$$

$$v_{\mathrm{leaf\_Star\_syn}} = v_{\mathrm{m\_leaf\_Star\_syn}} \cdot \frac{[\mathrm{TP}]_{\mathrm{leaf}}}{K_{\mathrm{m9}} + [\mathrm{TP}]_{\mathrm{leaf}}} \quad (6.2)$$

where, $K_{m8}$, $K_{m9}$ and $K_{e1}$ are Michaelis-constants, $v_{m\_leaf\_Suc\_syn}$ is leaf maximum sucrose synthesis (TP to sucrose conversion) rate, and $v_{m\_leaf\_Star\_syn}$ is leaf maximum starch synthesis (TP to starch conversion) rate (**Supplementary Table 2**).

In the night, sucrose level decreases and daily synthesized starch degrades gradually (Pilkington et al., 2015). There are a number of models simulating nighttime starch degradation, most of which incorporate a circadian clock control of the process (Feugier & Satake, 2013; Scialdone et al., 2013; Seaton, Ebenhöh, Millar, & Pokhilko, 2014). For simplification, we used a simple substrates feedback regulation mechanism to simulate starch degradation rate ($v_{\mathrm{leaf\_Star\_deg}}$):

$$v_{\mathrm{leaf\_Star\_deg}} = \begin{cases} v_{\mathrm{m\_leaf\_Star\_deg}} \cdot \dfrac{[\mathrm{Star}]_{\mathrm{leaf}}}{K_{\mathrm{m10}} + [\mathrm{Star}]_{\mathrm{leaf}}} \cdot \dfrac{[\mathrm{Suc}]_{\mathrm{leaf\_Star\_deg\_low}} - [\mathrm{Suc}]_{\mathrm{leaf}}}{[\mathrm{Suc}]_{\mathrm{leaf\_Star\_deg\_low}} + [\mathrm{Suc}]_{\mathrm{leaf}}}, [\mathrm{Suc}]_{\mathrm{leaf}} < [\mathrm{Suc}]_{\mathrm{leaf\_Star\_deg\_low}} \\ 0, [\mathrm{Suc}]_{\mathrm{leaf}} \geq [\mathrm{Suc}]_{\mathrm{leaf\_Star\_deg\_low}} \end{cases} \tag{6.3}$$

where, $K_{\mathrm{m10}}$ is Michaelis-constant, $[\mathrm{Suc}]_{\mathrm{leaf\_Star\_deg\_low}}$ is critical leaf sucrose concentration below which starch starts to decompose, $v_{\mathrm{m\_leaf\_Star\_deg}}$ is leaf maximum starch degradation rate (**Supplementary Table 2**).

*Leaf protein synthesis and degradation*: Leaf O-N and protein are in dynamic balance through their inter-conversion (conversion rate $v_{\mathrm{leaf\_O\text{-}N2Pro}} > 0$ means a conversion from O-N to protein, and *vice versa*):

$$v_{\mathrm{leaf\_O\text{-}N2Pro}} = \begin{cases} v_{\mathrm{m\_leaf\_Pro\_syn}} \cdot \dfrac{[\mathrm{O\text{-}N}]_{\mathrm{leaf}} - [\mathrm{O\text{-}N}]_{\mathrm{leaf\_Pro\_syn\_low}}}{K_{\mathrm{m11}} + [\mathrm{O\text{-}N}]_{\mathrm{leaf}}}, [\mathrm{O\text{-}N}]_{\mathrm{leaf}} > [\mathrm{O\text{-}N}]_{\mathrm{leaf\_Pro\_syn\_low}} \\ -v_{\mathrm{m\_leaf\_Pro\_deg}} \cdot \dfrac{[\mathrm{O\text{-}N}]_{\mathrm{leaf\_Pro\_syn\_low}} - [\mathrm{O\text{-}N}]_{\mathrm{leaf}}}{K_{\mathrm{m11}} + [\mathrm{O\text{-}N}]_{\mathrm{leaf}}}, [\mathrm{O\text{-}N}]_{\mathrm{leaf}} \leq [\mathrm{O\text{-}N}]_{\mathrm{leaf\_Pro\_syn\_low}} \end{cases} \tag{6.4}$$

where, $K_{\mathrm{m11}}$ is Michaelis-constant, $[\mathrm{O\text{-}N}]_{\mathrm{leaf\_Pro\_syn\_low}}$ is critical leaf O-N concentration below which protein synthesis ceases and proteins starts to decompose, $v_{\mathrm{m\_leaf\_Pro\_syn}}$ is leaf maximum protein synthesis rate and $v_{\mathrm{m\_leaf\_Pro\_deg}}$ is leaf maximum protein degradation rate (**Supplementary Table 2**).

*Leaf senescence***:** Similar to that of root, there is a minimal (constant) relative leaf photosynthetic area ($S_{\mathrm{leaf}}$) loss rate $\alpha_{\mathrm{leaf\_sene}}$ (**Supplementary Table 2**) due to aging (X. Yin & H. van Laar, 2005). As leaf senescence is closely related to leaf N remobilization during grain filling (Masclaux - Daubresse, Reisdorf - Cren, & Orsel, 2008; Sinclair & De Wit, 1976), we propose that leaf senescence rate ($v_{\mathrm{leaf\_sene}}$) would be accelerated as a result of nitrogen starvation when leaf total nitrogen level ($[\mathrm{N}]_{\mathrm{leaf}}$; including I-N, O-N and protein) decrease below a critical level $[\mathrm{N}]_{\mathrm{sene\_up}}$:

$$v_{\text{leaf\_sene}} = \begin{cases} \alpha_{\text{leaf\_sene}} \cdot e^{-\beta_{\text{leaf\_sene}} \cdot ([\text{N}]_{\text{leaf}} - [\text{N}]_{\text{sene\_up}})}, [\text{N}]_{\text{leaf}} < [\text{N}]_{\text{sene\_up}} \\ \alpha_{\text{leaf\_sene}}, [\text{N}]_{\text{leaf}} \geq [\text{N}]_{\text{sene\_up}} \end{cases} \quad (7.1)$$

where, $\beta_{\text{leaf\_sene}}$ is an empirical coefficient (**Supplementary Table 2**).

***Grains***

In WACNI, grain volume expansion and grain filling occur simultaneously rather than being divided into two distinct phases, as it has been reported in both maize and rice that time for expression of enzymes involved in these two processes overlaps, and protein/starch granules are presented in endosperm cells at the very beginning of the grain filling period (Cai et al., 2018; Ober, Setter, Madison, Thompson, & Shapiro, 1991; Ohdan et al., 2005; Ou-Lee & Setter, 1985; Toyosawa et al., 2016).

*Grain volume expansion*: Grain volume expansion, as a result of endosperm cell division, was modelled based on current grain surface area since cell division occurs mainly within several outer layer cells of grain (Olsen, 2004). Assuming that developing grains have ellipsoid shape with the width, height and length being $w$, $w$ and $w \cdot r$ ($r$ is grain length width ratio, **Supplementary Table 2**), respectively, grain volume ($V_{\text{grain}}$) and grain surface area ($S_{\text{grain}}$) were estimated as described below:

$$V_{\text{grain}} = \frac{4}{3}\pi \cdot w^3 \cdot r \quad (8.1)$$

$$S_{\text{grain}} \approx 4\pi\left(\frac{1+2r^{1.6075}}{3}\right)^{\frac{1}{1.6075}} \cdot w^2 \quad (8.2)$$

Glume size ($V_{\text{glume}}$) or and pericarp size can physically restrict growth of developing grains (Z. Yang, Van Oosterom, Jordan, & Hammer, 2009), although the strength of constraint may differ among species (Tashiro & Wardlaw, 1989). This constraint effect ($\alpha_{\text{restrict}}$) was modelled as described below:

$$\alpha_{\text{restrict}} = \begin{cases} 1, V_{\text{grain}} < (1-\varepsilon) \cdot V_{\text{glume\_max}} \\ \frac{1}{\varepsilon}\left(1 - \frac{V_{\text{grain}}}{V_{\text{glume\_max}}}\right), V_{\text{grain}} \geq (1-\varepsilon) \cdot V_{\text{glume\_max}} \end{cases} \quad (8.3)$$

where, $(1\text{-}\varepsilon) \cdot V_{\text{glume}}$ is a critical grain volume above which grain growth will be physically restrained.

In addition, grain volume expansion was modelled based on current grain surface area

($S_{grain}$) since cell division occurs mainly within several outer layer cells of grains (Olsen, 2004). We further propose that carbon and nitrogen supply, i.e., the concentrations of sucrose and nitrogen in grain aqueous space ($[O\text{-}N]_{grain}$ and $[Suc]_{grain}$), together determine grain volume expansion rate ($v_{grain\_grow}$):

$$v_{grain\_grow} = \alpha_{restrict} \cdot v_{m\_grain\_grow} \cdot S_{grain} \frac{[O\text{-}N]_{grain}}{K_{m12} + [O\text{-}N]_{grain}} \cdot \frac{[Suc]_{grain}}{K_{m13} + [Suc]_{grain}} \quad (8.4)$$

where, $K_{m12}$ and $K_{m13}$ are Michaelis-constants, $v_{m\_grain\_grow}$ is grain maximum growth rate (**Supplementary Table 2**).

*Grain starch and protein storage*: The aqueous space volume ($V_{grain\_aq}$) expands with the division of endosperm cells, and reduces by storage starch and protein (that are insoluble) accumulation:

$$V_{grain\_aq} = V_{grain} - \frac{m_{grain\_Star}}{\rho_{Star}} - \frac{m_{grain\_Pro}}{\rho_{Pro}} \quad (9.1)$$

where, $V_{grain}$ is total grain volume, $m_{grain\_Star}$ and $m_{grain\_Pro}$ are biomass and $\rho_{Star}$ and $\rho_{Pro}$ are density of accumulated starch and protein in grain, respectively.

For grain filling, storage starch and protein synthesis rate ($v_{grain\_Star\_syn}$ and $v_{grain\_Pro\_syn}$) were modeled based on the sucrose and O-N concentrations in grain aqueous space following a Michaelis-Menten equation:

$$v_{grain\_Star\_syn} = v_{m\_grain\_Star\_syn} \cdot \frac{[Suc]_{grain}}{K_{m14} + [Suc]_{grain}} \quad (9.2)$$

$$v_{grain\_Pro\_syn} = v_{m\_grain\_Pro\_syn} \cdot \frac{[O\text{-}N]_{grain}}{K_{m15} + [O\text{-}N]_{grain}} \quad (9.3)$$

where, $K_{m14}$ and $K_{m15}$ are Michaelis-constants, $v_{m\_grain\_Star\_syn}$ is grain maximum starch synthesis rate, $v_{m\_grain\_Pro\_syn}$ is grain maximum protein synthesis rate (**Supplementary Table 2**).

***Stem***

*Stem sucrose and starch, O-N and protein homeostasis*: Sucrose and starch, O-N and protein are in dynamic balance in stem as a result of inter-conversion between them, respectively. The conversion rates between them ($v_{stem\_Suc2Star}$ and $v_{stem\_O\text{-}N2Pro}$) were modeled as described below:

$$V_{\text{stem\_aq}} = \text{SV0} - \frac{m_{\text{stem\_Star}}}{\rho_{\text{Star}}} - \frac{m_{\text{stem\_Pro}}}{\rho_{\text{Pro}}} \quad (11.1)$$

$$v_{\text{stem\_Suc2Star}} = \begin{cases} v_{\text{m\_stem\_Star\_syn}} \cdot \dfrac{[\text{Suc}]_{\text{stem}} - [\text{Suc}]_{\text{stem\_Star\_syn\_low}}}{K_{\text{m16}} + [\text{Suc}]_{\text{stem}} - [\text{Suc}]_{\text{stem\_Star\_syn\_low}}}, [\text{Suc}]_{\text{stem}} > [\text{Suc}]_{\text{stem\_Star\_syn\_low}} \\ v_{\text{m\_stem\_Star\_deg}} \cdot \dfrac{[\text{Star}]_{\text{stem}}}{K_{\text{m17}} + [\text{Star}]_{\text{stem}}} \cdot \dfrac{[\text{Suc}]_{\text{stem}} - [\text{Suc}]_{\text{stem\_Star\_syn\_low}}}{K_{\text{m16}} + [\text{Suc}]_{\text{stem}} - [\text{Suc}]_{\text{stem\_Star\_syn\_low}}}, [\text{Suc}]_{\text{stem}} \le [\text{Suc}]_{\text{stem\_Star\_syn\_low}} \end{cases}$$

(11.2)

$$v_{\text{stem\_O-N2Pro}} = \begin{cases} v_{\text{m\_stem\_Pro\_syn}} \cdot \dfrac{[\text{O-N}]_{\text{stem}} - [\text{O-N}]_{\text{stem\_Pro\_syn\_low}}}{K_{\text{m18}} + [\text{O-N}]_{\text{stem}} - [\text{O-N}]_{\text{stem\_Pro\_syn\_low}}}, [\text{O-N}]_{\text{stem}} > [\text{O-N}]_{\text{stem\_Pro\_syn\_low}} \\ -v_{\text{m\_stem\_Pro\_deg}} \cdot \dfrac{[\text{Pro}]_{\text{stem}}}{K_{\text{m19}} + [\text{Pro}]_{\text{stem}}} \cdot \dfrac{[\text{O-N}]_{\text{stem}} - [\text{O-N}]_{\text{stem\_Pro\_syn\_low}}}{K_{\text{m18}} + [\text{O-N}]_{\text{stem}} - [\text{O-N}]_{\text{stem\_Pro\_syn\_low}}}, [\text{O-N}]_{\text{stem}} \le [\text{O-N}]_{\text{stem\_Pro\_syn\_low}} \end{cases}$$

(11.3)

where, $V_{\text{stem\_aq}}$ is the aqueous space volume of stem; $m_{\text{stem\_Star}}$ and $m_{\text{stem\_Pro}}$ represent the biomass of starch and protein in stem, respectively; $K_{\text{m16}}$, $K_{\text{m17}}$ $K_{\text{m18}}$ and $K_{\text{m19}}$ are Michaelis-constants; $v_{\text{m\_stem\_Star\_syn}}$ and $v_{\text{m\_stem\_Star\_deg}}$ are stem maximum starch synthesis and starch degradation rate, $v_{\text{m\_stem\_Pro\_syn}}$ and $v_{\text{m\_stem\_Pro\_deg}}$ are stem maximum protein synthesis and protein degradation rates, respectively (**Supplementary Table 2**).

*Xylem and phloem transport*: Phloem transport includes short distance transport that occurs at source or sink ends, i.e., material loading or unloading, and long distance transport along phloem sieve tubes. According to Münch's theory (Knoblauch et al., 2016), phloem long distance transport resistance is mainly determined by the physical property of the sieve tube and fluid in it. However, transporter proteins, whose efficiency is mainly determined by Michaelis-Menten kinetics, usually control the resistance of short distance transport. As solutes in xylem move with transpiration flow, which is not explicitly considered in our current model, solutes are assumed to be uniformly distributed along xylem path without concentration gradient. For phloem, we divided it into three different compartments and considered the transport resistance between them (see below).

*Flux between different phloem compartments*: Phloem path was divided into three compartments (see **Figure 1**), i.e., leaf phloem, grain phloem and root phloem. Flux in phloem is driven by osmotic pressure difference between the source and the sink; sucrose and amino acids are the main osmotic components in the phloem (Lalonde, Tegeder, Throne - Holst, Frommer, & Patrick, 2003). Flux rate ($v_{\mathrm{phloem\_flux\_leaf\_X}}$) between phloem compartments of leaf and organ X (X can be grain or root) was modeled as described below:

$$v_{\mathrm{phloem\_flux\_leaf\_X}} = \frac{([\mathrm{Suc}]_{\mathrm{leaf\_phloem}} + [\mathrm{O\text{-}N}]_{\mathrm{leaf\_phloem}}) - ([\mathrm{Suc}]_{\mathrm{X\_phloem}} + [\mathrm{O\text{-}N}]_{\mathrm{X\_phloem}})}{\mathrm{R}_{\mathrm{phloem\_leaf\_X}}} \qquad (12.1)$$

where, $\mathrm{R}_{\mathrm{phloem\_leaf\_X}}$ is the transport resistance between leaf phloem and X phloem (**Supplementary Table 2**).

*Loading and unloading*: The rate of substrate *s* loading and unloading at each source or sink organ X ($v_{\mathrm{X\_s\_l}}$ and $v_{\mathrm{X\_s\_ul}}$) were modeled with a simple equation which describes metabolites transport across membranes, similar to that in Y. Wang, Long, and Zhu (2014):

$$v_{\mathrm{X\_s\_l}} = v_{\mathrm{m\_X\_s\_l}} \cdot \frac{[\mathrm{s}]_{\mathrm{X}} - \dfrac{[\mathrm{s}]_{\mathrm{X}}}{K_{\mathrm{e\_X\_s\_l}}}}{K_{\mathrm{m\_X\_s\_l}} + [\mathrm{s}]_{\mathrm{X}}} \qquad (13.1)$$

$$v_{\mathrm{X\_s\_ul}} = v_{\mathrm{m\_X\_s\_ul}} \cdot \frac{[\mathrm{s}]_{\mathrm{tube}} - \dfrac{[\mathrm{s}]_{\mathrm{X}}}{K_{\mathrm{e\_X\_s\_ul}}}}{K_{\mathrm{m\_X\_s\_ul}} + [\mathrm{s}]_{\mathrm{tube}}} \qquad (13.2)$$

where, $K_{\mathrm{m\_X\_s\_l}}$ and $K_{\mathrm{m\_X\_s\_ul}}$ are Michaelis-constants; $v_{\mathrm{m\_X\_s\_l}}$ is the maximum loading rate and $v_{\mathrm{m\_X\_s\_ul}}$ is the maximum unloading rate of substrate *s* at organ X; $[\mathrm{s}]_{\mathrm{X}}$ and $[\mathrm{s}]_{\mathrm{tube}}$ are the concentrations of substrate *s* in organ X and in xylem or phloem tube, respectively.

These loading and unloading processes include root-to-shoot I-N and O-N loading (to xylem), shoot-to-root sucrose and O-N unloading (from phloem) at root; photosynthetic sucrose and O-N loading (to phloem), I-N and O-N unloading (from xylem) at leaf; sucrose and O-N unloading (from phloem) at grain; and xylem-to-phloem O-N transfer (see **Figure 1**, process 13).

*Phloem-stem diffusion*: As a simplification, stem was set to be located around leaf phloem only. Substrates (sucrose and O-N in this model) diffusion rate ($v_{\mathrm{phloem\_stem\_s\_d}}$) between leaf phloem and stem were determined by their diffusion property, similar to that in Y. Wang et al. (2014):

$$v_{\mathrm{phloem\_stem\_s\_d}} = \frac{[\mathrm{s}]_{\mathrm{phloem}} - [\mathrm{s}]_{\mathrm{stem}}}{\mathrm{R}_{\mathrm{\_phloem\_stem\_s}}} \qquad (14.1)$$

where, $[\mathrm{s}]_{\mathrm{phloem}}$ and $[\mathrm{s}]_{\mathrm{stem}}$ are the concentration of substrate *s* in leaf phloem and stem, respectively; $\mathrm{R}_{\mathrm{\_phloem\_stem\_s}}$ is the transfer resistance of substrate *s* (**Supplementary Table 2**).

***Respiration***

The respiratory rate was calculated for each organ ($R_{\mathrm{d\_root}}$, $R_{\mathrm{d\_leaf}}$, $R_{\mathrm{d\_grain}}$, $R_{\mathrm{d\_stem}}$), following the concept proposed by Cannell and Thornley (Cannell & Thornley, 2000; Thornley & Cannell, 2000):

$$\begin{aligned} R_{\mathrm{d\_root}} = [&\eta_{\mathrm{l}} \cdot (v_{\mathrm{root\_IN\_l}} + v_{\mathrm{root\_ON\_l}} + v_{\mathrm{root\_ON\_ul}} + v_{\mathrm{root\_Suc\_ul}}) + \eta_{\mathrm{N\_upt}} \cdot v_{\mathrm{root\_N\_upt}} \\ &+ \eta_{\mathrm{N\_ass}} \cdot v_{\mathrm{root\_N\_ass}} + \eta_{\mathrm{grow}} \cdot v_{\mathrm{root\_grow}}] \cdot \mathrm{RW} + \gamma_{\mathrm{root\_residual}} \cdot V_{\mathrm{root\_aq}} \end{aligned} \qquad (15.1)$$

$$\begin{aligned} R_{\mathrm{d\_leaf}} = [&\eta_{\mathrm{l}} \cdot (v_{\mathrm{leaf\_ON\_l}} + v_{\mathrm{leaf\_Suc\_l}} + v_{\mathrm{leaf\_IN\_ul}} + v_{\mathrm{leaf\_ON\_ul}}) + \eta_{\mathrm{store}} \cdot v_{\mathrm{leaf\_Star\_deg}} \\ &+ \eta_{\mathrm{Pro\_syn}} \cdot v_{\mathrm{leaf\_Pro\_syn}} + \eta_{\mathrm{Pro\_deg}} \cdot v_{\mathrm{leaf\_Pro\_deg}}] \cdot \mathrm{LA} + \gamma_{\mathrm{residual}} \cdot V_{\mathrm{leaf\_aq}} \end{aligned} \qquad (15.2)$$

$$\begin{aligned} R_{\mathrm{d\_grain}} = [&\eta_{\mathrm{ul}} \cdot (v_{\mathrm{grain\_ON\_ul}} + v_{\mathrm{grain\_Suc\_ul}}) + \eta_{\mathrm{grow}} \cdot v_{\mathrm{grain\_grow}}] \cdot S_{\mathrm{grain}} + (\eta_{\mathrm{store}} \cdot v_{\mathrm{grain\_Star\_syn}} \\ &+ \eta_{\mathrm{Pro\_syn}} \cdot v_{\mathrm{grain\_Pro\_syn}}) \cdot V_{\mathrm{grain\_aq}} + \gamma_{\mathrm{residual}} \cdot V_{\mathrm{grain}} \end{aligned} \qquad (15.3)$$

$$\begin{aligned} R_{\mathrm{d\_stem}} = [&\eta_{\mathrm{store}} \cdot (v_{\mathrm{stem\_Star\_deg}} + v_{\mathrm{stem\_Star\_syn}}) + \eta_{\mathrm{Pro\_syn}} \cdot v_{\mathrm{stem\_Pro\_syn}} + \eta_{\mathrm{Pro\_deg}} \cdot v_{\mathrm{stem\_Pro\_deg}} \\ &+ \gamma_{\mathrm{residual}}] \cdot V_{\mathrm{stem\_aq}} \end{aligned} \qquad (15.4)$$

where, $\eta_{\mathrm{l}}$, $\eta_{\mathrm{ul}}$, $\eta_{\mathrm{N\_upt}}$, $\eta_{\mathrm{N\_ass}}$, $\eta_{\mathrm{grow}}$, $\eta_{\mathrm{store}}$, $\eta_{\mathrm{Pro\_deg}}$ and $\eta_{\mathrm{Pro\_syn}}$ are respiration coefficients of loading, unloading, soil I-N uptake, N assimilation, growth, starch degdaration or synthesis, protein degradation and protein synthesis processes, respectively; $v_{\mathrm{X\_IN\_l}}$ and $v_{\mathrm{X\_ON\_l}}$ and $v_{\mathrm{X\_Suc\_l}}$, $v_{\mathrm{X\_IN\_ul}}$ and $v_{\mathrm{X\_ON\_ul}}$ and $v_{\mathrm{X\_Suc\_ul}}$, $v_{\mathrm{X\_N\_upt}}$, $v_{\mathrm{X\_N\_ass}}$, $v_{\mathrm{X\_grow}}$, $v_{\mathrm{X\_Star\_deg}}$ and $v_{\mathrm{X\_Star\_syn}}$, $v_{\mathrm{X\_Pro\_deg}}$ and $v_{\mathrm{X\_Pro\_syn}}$ are the corresponding reaction rates in organ X;

$V_{root\_aq}$, $V_{leaf\_aq}$, $V_{grain\_aq}$ and $V_{stem\_aq}$ are the aqueous space volume of root, leaf, grain and stem, respectively; $\gamma_{residual}$ is the residual respiration (respiration for other processes) coefficient for leaf, stem and grain; $\gamma_{root\_residual}$ is the residual respiration coefficient for root, which is twice larger than $\gamma_{residual}$ as a compensation for potential root exudation that is not explicitly represented in the model (**Supplementary Table 2**).

#### Model performance on multiple datasets

##### *Simulating the effects of different nitrogen fertilizer application rates on rice grain filling (Figure 2a-d; Supplementary Figure 1)*

To examine effects of different nitrogen fertilizer application rates on rice grain filling, experimental data were taken from Zhao et al. (2015) (**Supplementary Datasets 2**). These authors had reported four nitrogen top-dressing treatments during the ear development period, i.e., no nitrogen; low level (0.6 g nitrogen/pot), medium level (1.2 g nitrogen/pot), and high level (1.8 g nitrogen/pot) nitrogen. In simulation, soil nitrogen concentrations were set to be 20%, 50%, 100% and 150% of the default value for the four nitrogen treatments, respectively, with the assumption that there was still 20% of nitrogen in soil when no nitrogen was applied. Based on data in *Table 1* in Zhao et al. (2015), the ratio of leaf dry matter and dry weight of grains per plant among the four nitrogen treatments at 7 days after anthesis was 0.60 : 0.90 : 1.00 : 1.23, and 0.43 : 0.75 : 1.00 : 1.18, respectively. Therefore, we set the ratio of tiller number between these groups as that of leaf dry matter, i.e., 0.60 : 0.90 : 1.00 : 1.23, and set the ratio of grain number per ear between these groups by dividing dry weight of the grains by tiller number, i.e., 0.43/0.60 : 0.75/0.90 : 1.00 : 1.18/1.23. According to *Table 2* in Zhao et al. (2015), ratio of leaf protein amino acids per plant between the four nitrogen treatments at 7 days after anthesis were 0.40 : 0.75 : 1.00 : 1.21. Therefore, we set the ratio of leaf protein concentration between groups by dividing leaf protein amino acids by leaf dry matter, i.e., 0.40/0.60 : 0.75/0.90 : 1.00 :

1.21/1.23. We further set the same ratio of stem protein concentration between groups as that of leaf.

To compare the physiological traits during the grain filling period between model simulation and experimental data, the grain nitrogen concentration data were extracted from *Table 2* in Zhao et al. (2015), and the values of total amino acids were used for comparison; the leaf nitrogen concentration data were calculated by dividing the amount of leaf total amino acids by leaf dry matter from *Table 1* and *Table 2* in Zhao et al. (2015), and the grain dry weight data were extracted from *Table 1* in Zhao et al. (2015). For both model simulation and experimental data, the values of the leaf nitrogen concentration, the grain nitrogen concentration at 7 days after flowering, and the grain dry weight at harvest of the medium nitrogen treatment were used as a reference, and data at all time points and in all nitrogen treatment groups were normalized by dividing these values, respectively. Extracted experimental data are tabulated in **Supplementary Datasets 2**.

***Simulating the effects of shading and thinning on rice grain filling (Figure 2e-h; Supplementary Figure 2)***

To examine the effects of different light regimes on rice grain filling, the experimental data were collected from Kobata, Sugawara, and Takatu (2000) (**Supplementary Datasets 2**). The authors reported seven different shading/thinning treatments after flowering. Firstly, control plots were grown under normal condition throughout the entire grain filling period. Concurrently, three shading treatments were applied immediately after flowering, i.e., the heavy shaded treatment using double black cloth, the moderate shaded treatment using single black cloth, and the light shaded treatment using white cloth, which reduced full sun radiation by 74.4%, 48.1% and 25.1%, respectively. The shade frames were removed 10 days later. Then, half the plots were left untouched (**Figure 2e-h**), while the other half were thinned to every other plant to reduce the plant density by half for the rest of the grain filling period

(**Supplementary Figure 2**).

Default model parameters were used during simulation. Incident solar light intensity during the firstly 10 days after flowering were reduced to 75%, 50% and 25% for light, medium and heavy shaded treatments, respectively; and the planting density beyond 10 days after flowering were halved for the thinning treatment groups. The experimental data of grain dry weight changes during the grain filling period for all seven treatments were extracted from *Fig. 2* in Kobata et al. (2000). For both model simulation and experimental data, the values of grain dry weight at harvest of the "normal condition" groups were used as a reference, and data at all time points and for all treatments were normalized by dividing them, respectively. The extracted experimental data are tabulated in **Supplementary Datasets 2**.

***Simulating the interaction effects of different nitrogen fertilizer application rates and air $CO_2$ concentrations on rice grain filling (Figure 2i-j)***

For simulating the interaction effects of different nitrogen fertilizer application rates and air $CO_2$ concentrations on rice grain filling, experimental data were collected from Kim, Lieffering, Miura, Kobayashi, and Okada (2001) (**Supplementary Datasets 2**). The authors reported six different combinations of nitrogen fertilizer application rate and air $CO_2$ concentration treatments. Namely, nitrogen was supplied as ammonium sulphate at three application rates: 4 g $m^{-2}$ (low; "LN"), 8-9 g $m^{-2}$ (medium; "MN") and 12-15 g $m^{-2}$ (high; "HN"), respectively; air $CO_2$ concentration was controlled at two levels: ambient level of around 390 μmol $mol^{-1}$, free-air $CO_2$ enrichment of 230-365 μmol $mol^{-1}$ higher than that under ambient level ("FACE"). Correspondingly, in the simulation, we set the soil nitrogen concentrations to be 50%, 100% and 150% of the default value for "LN", "MN" and "HN" treatments, respectively; the air $CO_2$ concentrations were set to 390 and 690 μmol $mol^{-1}$ for the ambient and the "FACE" treatments, respectively.

According to *Table 1* in Kim et al. (2001), it is known that the ratio of plant total nitrogen, plant root dry matter and plant total dry matter between “LN”, “MN”, “HN”, “LN+FACE”, “MN+FACE” and “HN+FACE” groups at ear initiation were 0.83 : 1.00 : 1.25 : 0.85 : 1.15 : 1.53, 1.00 : 1.00 : 1.03 : 1.18 : 1.46 : 1.35 and 0.87 : 1.00 : 1.06 : 1.21 : 1.45 : 1.51, respectively. Therefore, the ratio of root weight between these groups at flowering was set as that of plant root dry matter measured at ear initiation; the ratio of leaf area and pre-flowering stem stored starch at flowering was set to be the same as that of measured plant total dry matter at ear initiation; and the ratio of leaf protein concentration and pre-flowering stem stored protein at flowering was set by dividing plant total nitrogen by plant total dry matter, i.e., 0.83/0.87 : 1.00 : 1.25/1.06 : 0.85/1.21 : 1.15/1.45 : 1.53/1.51.

According to *Table 2* in Kim et al. (2001), it is known that ratio of ear number, fertile spikelets and individual grain mass between “LN”, “MN”, “HN”, “LN+FACE”, “MN+FACE” and “HN+FACE” groups at harvest were 0.87 : 1.00 : 1.07 : 0.89 : 1.09 : 1.15, 0.83 : 1.00 : 1.13 : 0.85 : 1.09 : 1.30, and 1.07 : 1.00 : 0.98 : 1.09 : 1.04 : 0.93, respectively. Therefore, the ratio of tiller number and grain size between these groups was set as that of ear number and individual grain mass measured at harvest, respectively; the ratio of grain number per ear between these groups was set by dividing fertile spikelets by tiller number, i.e., 0.83/0.87 : 1.00 : 1.13/1.07 : 0.85/0.89 : 1.09/1.09 : 1.30/1.15. The ratio of leaf nitrogen concentration at flowering was set by dividing plant nitrogen uptake by plant total dry matter at ear initiation, which was 0.95 : 1.00 : 1.18 : 0.70 : 0.79 : 1.01. In addition, for the “MN” group, the measured ear number, the spikelet number per ear, and the fertiled rate of spikelets are 454.74 $m^{-2}$, 84 and 94.2%, respectively. As the default model parameters for ear number and fertiled rate of spikelets are 300 $m^{-2}$ and 87.5%, respectively, an equivalent grain number per ear was set to 137 (137 = 454.74*84*94.2%/300/87.5%) for the “MN” group.

During comparison of model prediction and experimental data, the grain yield at harvest was extracted from *Table 2* in Kim et al. (2001), the root nitrogen uptake throughout the grain filling period was approximated by the difference of plant total nitrogen at harvest and plant total nitrogen at ear initiation from *Table 1* in Kim et al. (2001). We simulated two scenarios, one was using adjusted parameters as above mentioned, with other parameters using the default values (the "Simulation" group in Figure 3t, u); the other one was with a further assumption that maximum root nitrogen uptake rate under "FACE" was reduced by 36% compared to that under ambient condition (the "Simulation ($v_{m\_Nupt}$=64% under FACE)" group in **Figure 2i, j**). For both model simulation and experimental data, the values of grain yield and root nitrogen uptake in the "MN" group were used as a reference, and data at all time points and for all treatments were normalized by dividing them, respectively. The extracted experimental data are tabulated in **Supplementary Datasets 2**.

##### *Simulating the effects of genetic manipulation of leaf senescence rate on rice grain filling (Figure 2k-m; Supplementary Figure 3)*

To examine the effects of genetic manipulation of leaf senescence rate on rice grain filling, the experimental data were collected from Liang et al. (2014) (**Supplementary Datasets 2**). The authors reported a gain-of-function mutant prematurely senile 1 (*ps1-D*) that demonstrated significant premature leaf senescence. Further genetic study shows that *PS1* encodes a plant-specific NAC transcription activator, *OsNAP*, the overexpression of which significantly promoted senescence, whereas knockdown of which would delay leaf senescence. Default model parameters were used for simulating wild type plants ("WT"), except that the $v_{m\_leaf\_Pro\_deg}$ was set to 50% of the default value to adapt to the observed leaf senescence pattern of "WT". According to *Fig. 1C* in Liang et al. (2014), the expression of chlorophyll degradation and leaf senescence process related genes in "*ps1-D* mutant" increased to 3-5 folds to that of "WT"; and according to *Fig. 4B-C* in Liang et al. (2014), *OsNAP* gene expression level in the leaf of "*OsNAP* RNAi" decreased to 0.2-0.3 fold, while ABA content increased to 1.4-1.9 folds to that of "WT". Therefore, we set $v_{m\_leaf\_Pro\_deg}$ of "*ps1-D* mutant" and "*OsNAP* RNAi" to be 0.5-fold and 4-folds of "WT", respectively.

According to *Table S4* in Liang et al. (2014), the ratio of dry weight of vegetative organs among "WT", "*ps1-D* mutant" and "*OsNAP* RNAi" groups was 1 : 0.74 : 1.11. Therefore, the ratio of plant size between these groups, including leaf area, root weight, stem stored starch/protein and grain number per ear, were set the same as that for dry weight of vegetative organs, i.e., 1 : 0.74 : 1.11.

To enable comparison between model predictions and experimental data, we extracted the leaf chrolophyll content dynamic changes of "WT" and "*ps1-D* mutant" during the grain filling period from *Fig. 1B* in Liang et al. (2014), which was used to represent relative change of leaf nitrogen concentration. The grain yield and grain nitrogen concentration at harvest were extracted from *Fig. 4F* and *Fig. 5B* in Liang et al. (2014), respectively. For both model simulation and experimental data, the values of leaf nitrogen concentration at flowering, and the grain yield and grain nitrogen concentration at harvest of "WT" were used as a reference, i.e., data at all time points and for other plants were normalized by dividing them, respectively. The extracted experimental data are tabulated in **Supplementary Datasets 2**.

***Simulating the effects of genetic manipulation of phloem-to-grain sucrose unloading capacity on rice grain filling (Figure 2n-o; Supplementary Figures 4, 5)***

To examine the effects of genetic manipulation of phloem-to-grain sucrose unloading capacity on rice grain filling, the experimental data were collected from E. Wang et al. (2008) (**Supplementary Datasets 2**). The authors reported a loss-of-function mutant grain incomplete filling 1 ("*gif1* mutant") that showed slower grain-filling rate than wild-type rice ("WT"). Further genetic study shows that *GIF1* encodes a cell-wall invertase, which is responsible for sucrose unloading into grain, particularly during early grain-filling.

According to *Figure 1h-j* in E. Wang et al. (2008), the sugar content at 5 days after pollination in "*gif1* mutant" decreased to 0.4-0.7 fold to that of "WT". Therefore, the maximum phloem-to-grain sucrose unloading rate of "*gif1* mutant" was set to be 0.5-

fold of “WT”; the maximum phloem-to-grain sucrose unloading rate of plants overexpressing *GIF1* from its native promoter (“*GIF1* OE”) was set to be 2-fold of “WT”.

According to *Figure 1g* and *Figure 4b* in E. Wang et al. (2008), the ratio of single filled grain dry weight at harvest between “WT”, “*gif1* mutant” and “*GIF1* OE” groups was 1 : 0.76 : 1.11. Therefore, the ratio of grain size between these groups was set to be the same as that for the measured grain dry weight at harvest, i.e., 1 : 0.76 : 1.11. The extracted experimental data are tabulated in **Supplementary Datasets 2**.

For both model simulation and experimental data, the grain dry weight, straw dry weight and plant biomass at harvest of “WT” were used as a reference, i.e., data at all time points and for other plants were normalized by dividing them, respectively. Note the plant grain yield, straw dry weight and plant biomass at harvest were measured in this study (see in **Methods**).

To test the effects of changing phloem-to-grain sucrose unloading capacity on plant physiological changes during the grain filling period for “sink limited” plants (with default model parameter settings except that the grain number per ear is set to 145), we simulated the grain dry weight, leaf area, leaf nitroge concentration, canopy photosynthesis, root dry weight and root nitrogen uptake throughout the grain filling period for plants with the maximum phloem-to-grain sucrose unloading rates of 200%, 100% and 50% to that of the default value (**Supplementary Figure 4**). For the “source limited” case, the grain number per ear is set to 175 (**Supplementary Figure 5**).

***Simulating the effects of genetic manipulation of grain starch synthesis activity on rice grain filling (Figure 2p)***

To examine the effects of genetic manipulation of grain starch synthesis activity on rice grain filling, the experimental data were collected from Smidansky, Martin,

Hannah, Fischer, and Giroux (2003) (**Supplementary Datasets 2**). The authors reported that transforming rice with a modified maize AGPase large subunit sequence (*Sh2r6hs*) in an endosperm-specific manner could specifically enhance activity of starch synthesis in endosperm. According to *Table 4* in Smidansky et al. (2003), the ratio of grain number per ear and tiller number between "WT" and *Sh2r6hs* transgene groups were 1 : 1.13 and 1 : 1.05, respectively. The ratio of grain size between two groups was set to be the same as that for grain weight, i.e., 1: 1.02. The ratio of leaf area, root weight, stem structural weight, stem starch storage and stem protein storage was set by dividing the total plant weight by the tiller number, i.e., 1 : 1.22/1.05. According to *Table 2* in Smidansky et al. (2003), the measured AGPase activity was 9-52% higher in *Sh2r6hs* transgenic grains than in untransformed grains between 10-20 days after anthesis. Therefore, the maximum grain starch synthesis rate $v_{\mathrm{m\_grain\_Star\_syn}}$ of *Sh2r6hs* transgenic plants was set to be 1.5-fold of "WT".

In total three scenarios were simulated, the first one was using adjusted plant size and $v_{\mathrm{m\_grain\_Star\_syn}}$ as above mentioned, with other parameters using the default values (the grey bars in **Figure 2p**); the second one was using only adjusted plant size with unchanged $v_{\mathrm{m\_grain\_Star\_syn}}$ (the black bars in **Figure 2p**); the third one was using only adjusted $v_{\mathrm{m\_grain\_Star\_syn}}$ with unchanged plant size (the white bars in **Figure 2p**).

To enable comparison between model predictions and experimental data, we extracted the grain yield, grain nitrogen concentration and harvest index at harvest of positive homozygotes and negative homozygotes of *Sh2r6hs* transgenic plants from *Table 4* in Smidansky et al. (2003). For both model simulation and experimental data, the grain yield, grain nitrogen concentration and harvest index at harvest of negative homozygotes of *Sh2r6hs* transgenic plants were used as a reference, i.e., data at all time points and for other plants were normalized by dividing them, respectively. The extracted experimental data are tabulated in **Supplementary Datasets 2**.

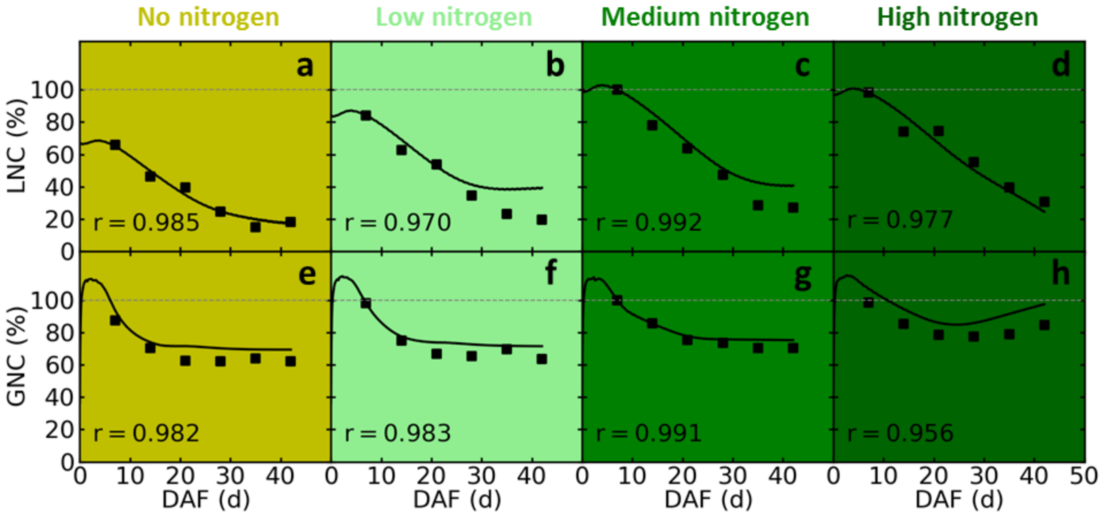


**Supplementary Figure 1. Dynamic changes of leaf nitrogen concentration (LNC) and grain nitrogen concentration (GNC) during grain filling under different nitrogen regimes.**

The filled squares are measured data from Zhao et al. (2015), the solid lines are simulated data. The dynamic changes of grain dry weight were shown in **Figure 2a-d**.

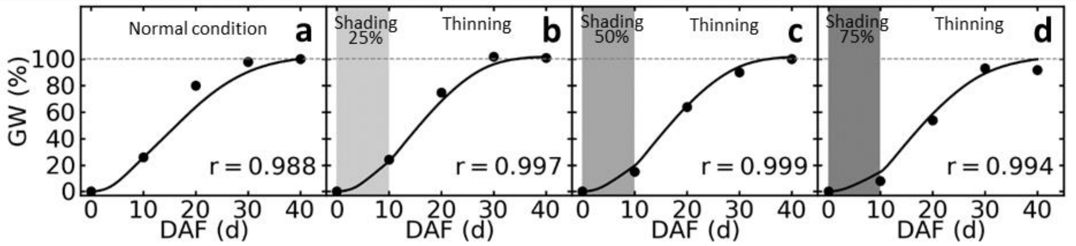


**Supplementary Figure 2. Dynamic changes of grain dry weight during grain filling under different light regimes.**

The filled circles are measured data from Kobata et al. (2000), the solid lines are simulated data. The dynamic changes of grain dry weight without plant thinning were shown in **Figure 2e-h**.

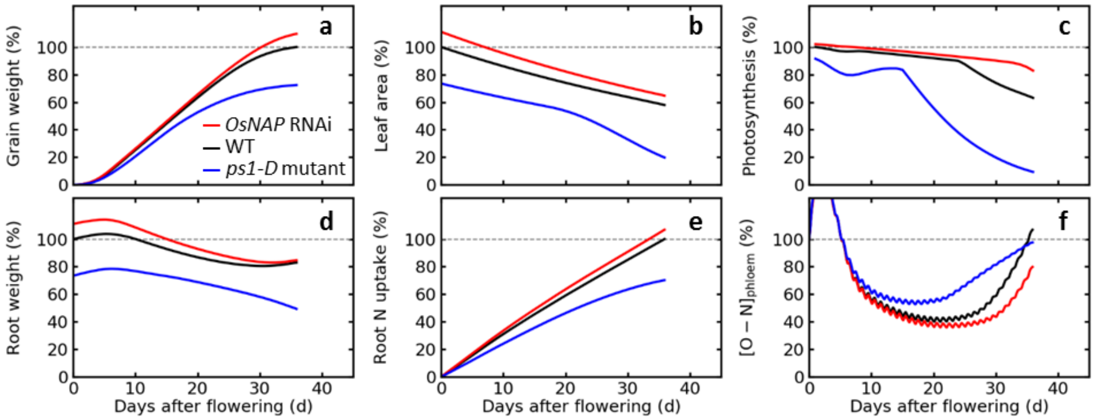


**Supplementary Figure 3. Predicted changes of plant physiological and agronomic traits for wild type (WT), *ps1-D* mutant and *OsNAP* RNAi plants during grain filling.**

The traits include plant grain weight (**a**), leaf area (**b**), daily total photosynthesis (**c**), root weight (**d**), total accumulated root nitrogen uptake (**e**), and phloem O-N concentration (**f**).

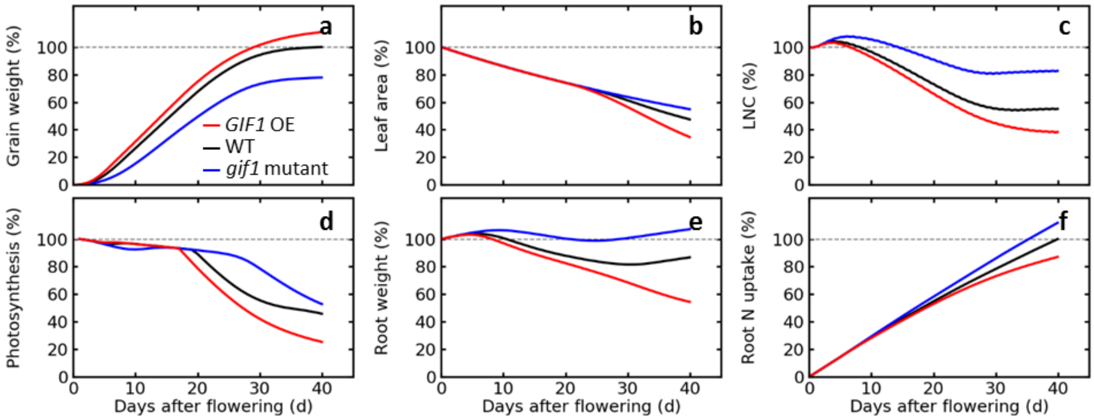


**Supplementary Figure 4. Predicted changes of plant physiological and agronomic traits for *in silico* wild type (WT), *gif1* mutant and *GIF1* overexpression plants in a "sink limited" scenario.**

In this scenario, the grain number per ear for WT was set to 145. The traits include plant grain weight (**a**), leaf area (**b**), leaf nitrogen concentration (**c**), daily total photosynthesis (**d**), root weight (**e**), total accumulated root nitrogen uptake (**f**).

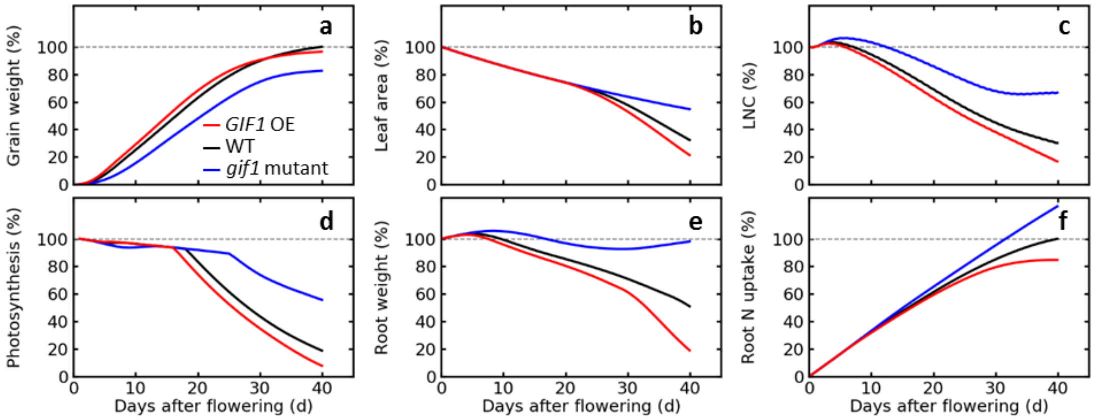


**Supplementary Figure 5. Predicted changes of plant physiological and agronomic traits for *in silico* wild type (WT), *gif1* mutant and *GIF1* overexpression plants in a "source limited" scenario.**

In this scenario, the grain number per ear for WT was set to 175. The traits include plant grain weight (**a**), leaf area (**b**), leaf nitrogen concentration (**c**), daily total photosynthesis (**d**), root weight (**e**), total accumulated root nitrogen uptake (**f**).

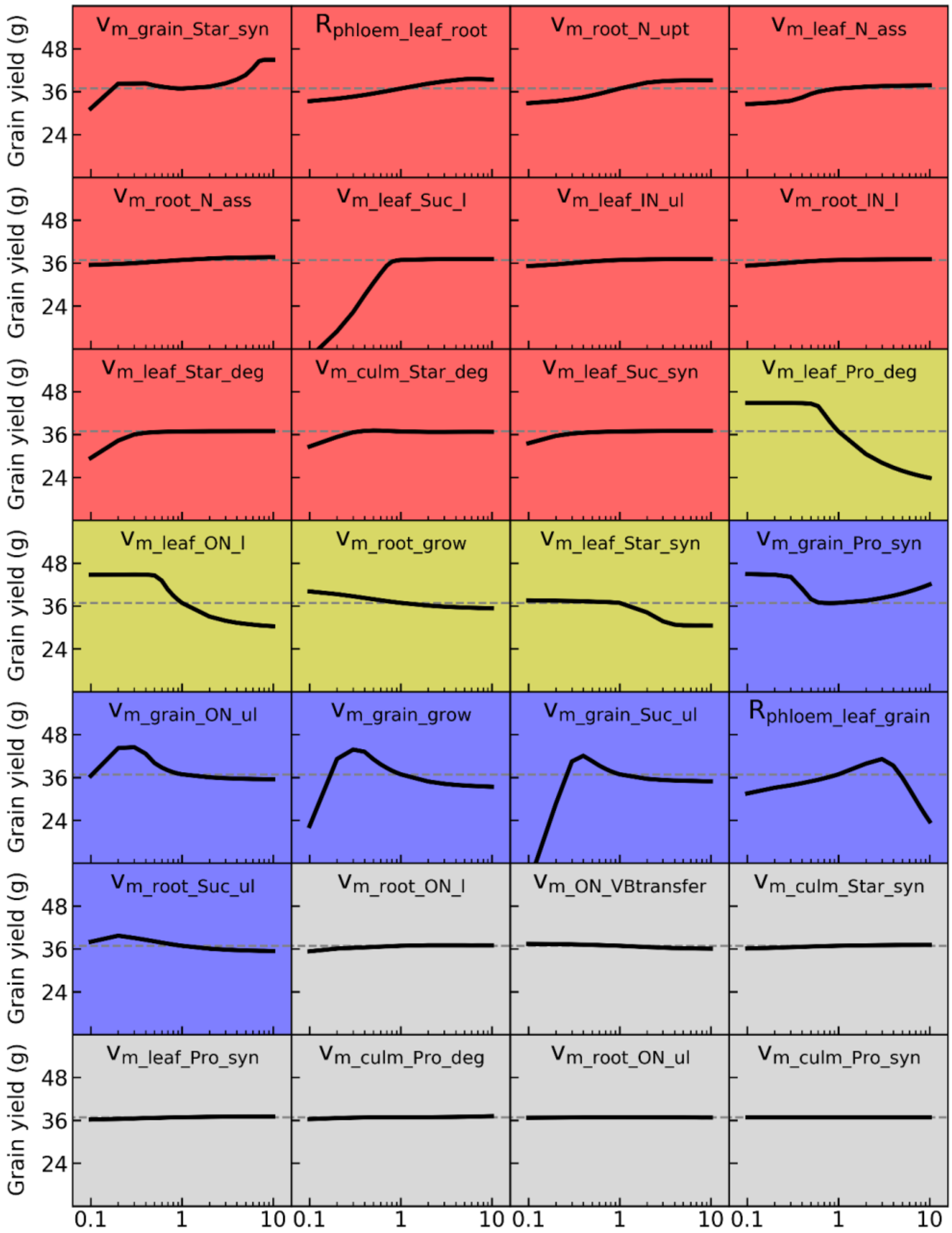


**Supplementary Figure 6. Predicting and grouping effects of manipulating capacities of 28 reaction and diffusion processes on grain yield.**
Eleven parameters shown on the red background that monotonically increase grain yield when they increase from 0.1-fold to 10-fold of their default values, were termed as universal yield enhancers (UYEs). Four parameters shown on the yellow background that monotonically decrease grain yield when they increase, were termed as universal yield inhibitors (UYIs). Six parameters shown on the blue background that non-linearly affect grain yield, were termed as conditional yield enhancers (CYEs). Seven parameters shown on the grey background having negligible influences on grain yield, were termed as weak yield regulators (WYRs).

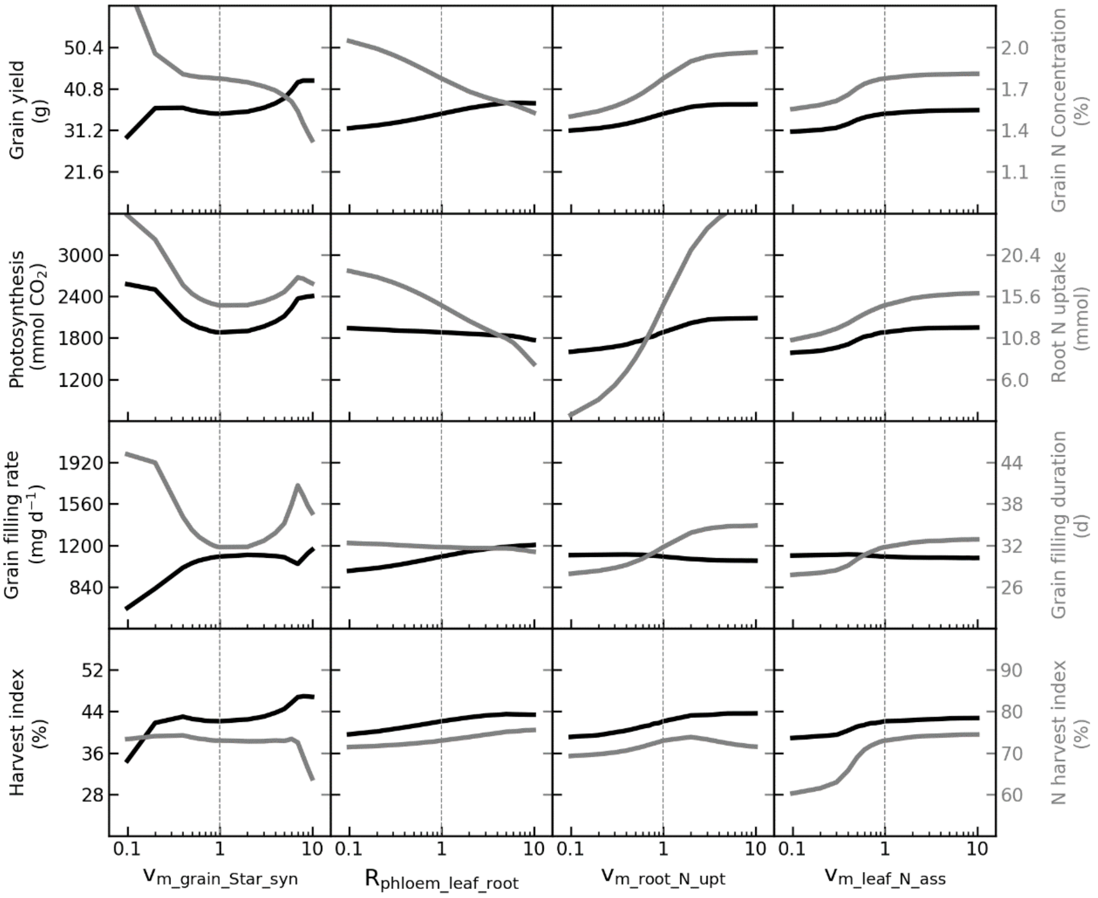


**Supplementary Figure 7. The responses of major agronomic traits to variation of model parameters $v_{m\_grain\_Star\_syn}$, $R_{\_phloem\_leaf\_root}$, $v_{m\_root\_N\_upt}$, $v_{m\_leaf\_N\_ass}$, respectively.**

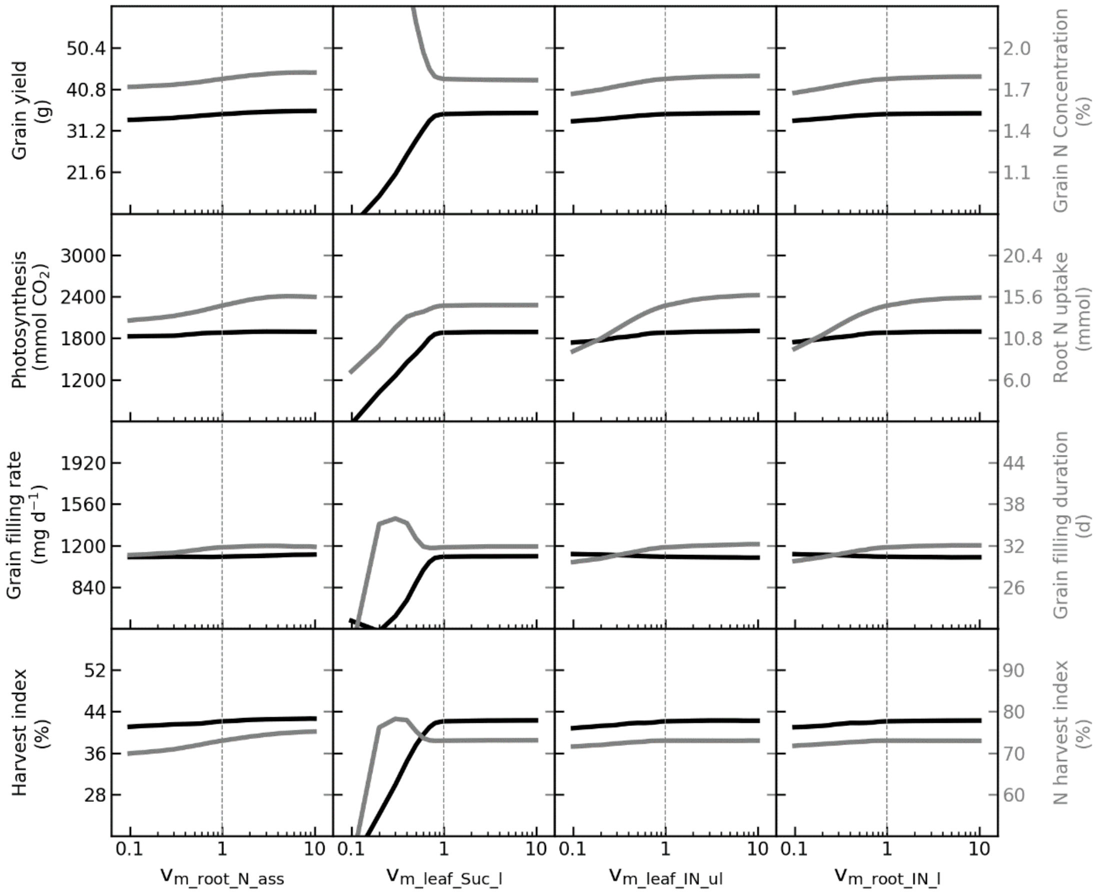


**Supplementary Figure 8. The responses of major agronomic traits to variation of model parameters $v_{m\_root\_N\_ass}$, $v_{m\_leaf\_Suc\_l}$, $v_{m\_leaf\_IN\_ul}$, $v_{m\_root\_IN\_l}$, respectively.**

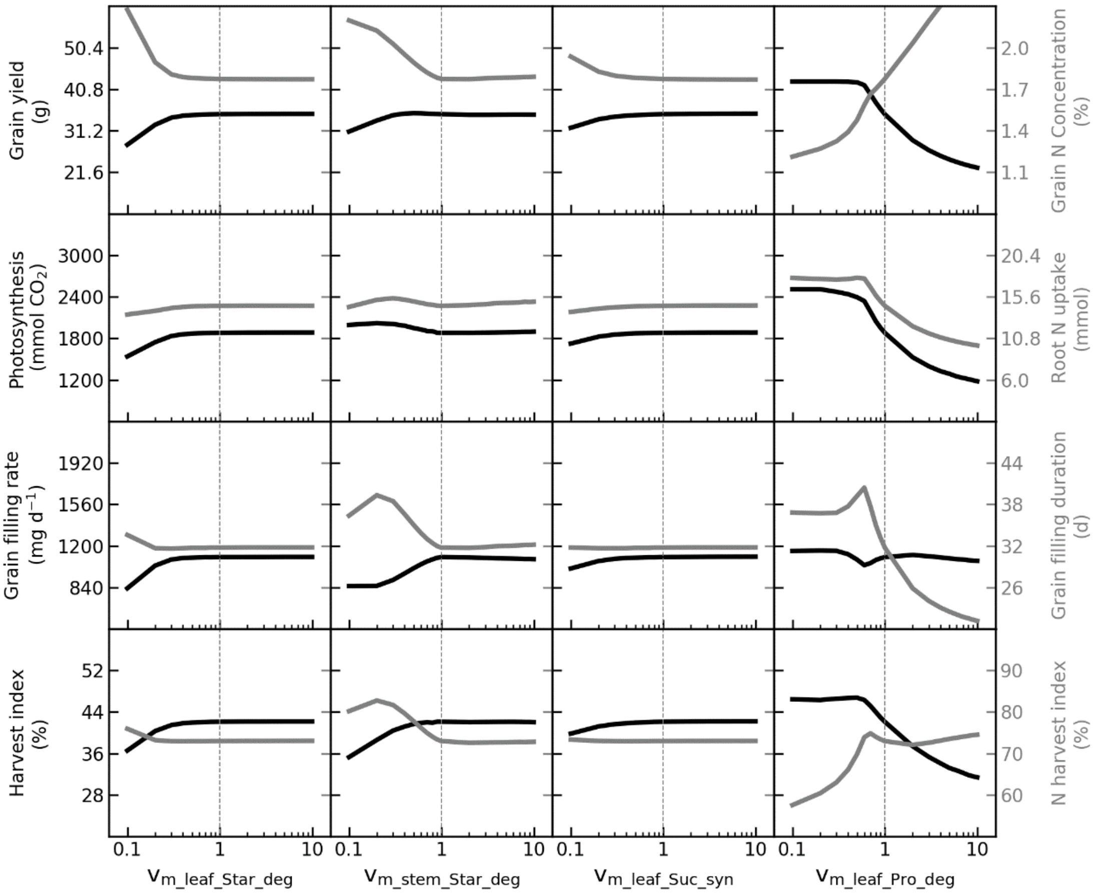


**Supplementary Figure 9. The responses of major agronomic traits to variation of model parameters $v_{m\_leaf\_Star\_deg}$, $v_{m\_stem\_Star\_deg}$, $v_{m\_leaf\_Suc\_syn}$, $v_{m\_leaf\_Pro\_deg}$, respectively.**

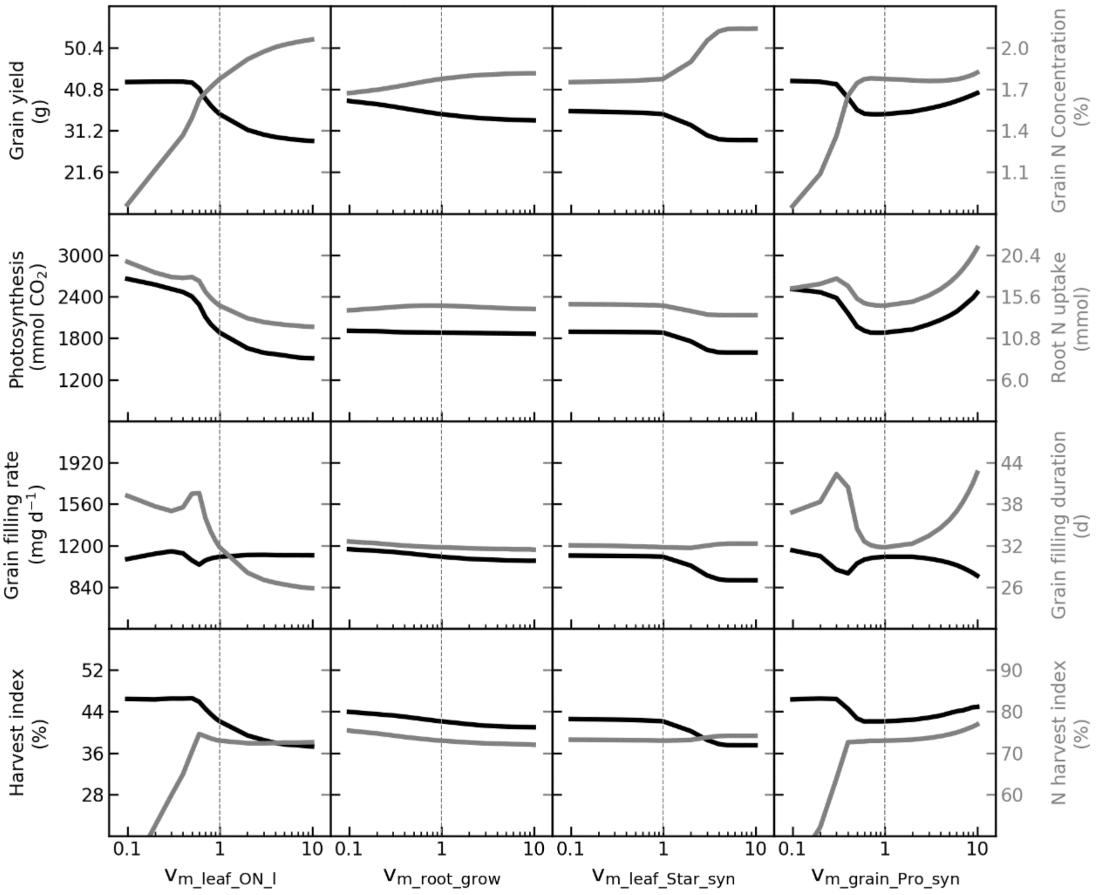


**Supplementary Figure 10. The responses of major agronomic traits to variation of model parameters $V_{m\_leaf\_ON\_l}$, $V_{m\_root\_grow}$, $V_{m\_leaf\_Star\_syn}$, $V_{m\_grain\_Pro\_syn}$, respectively.**

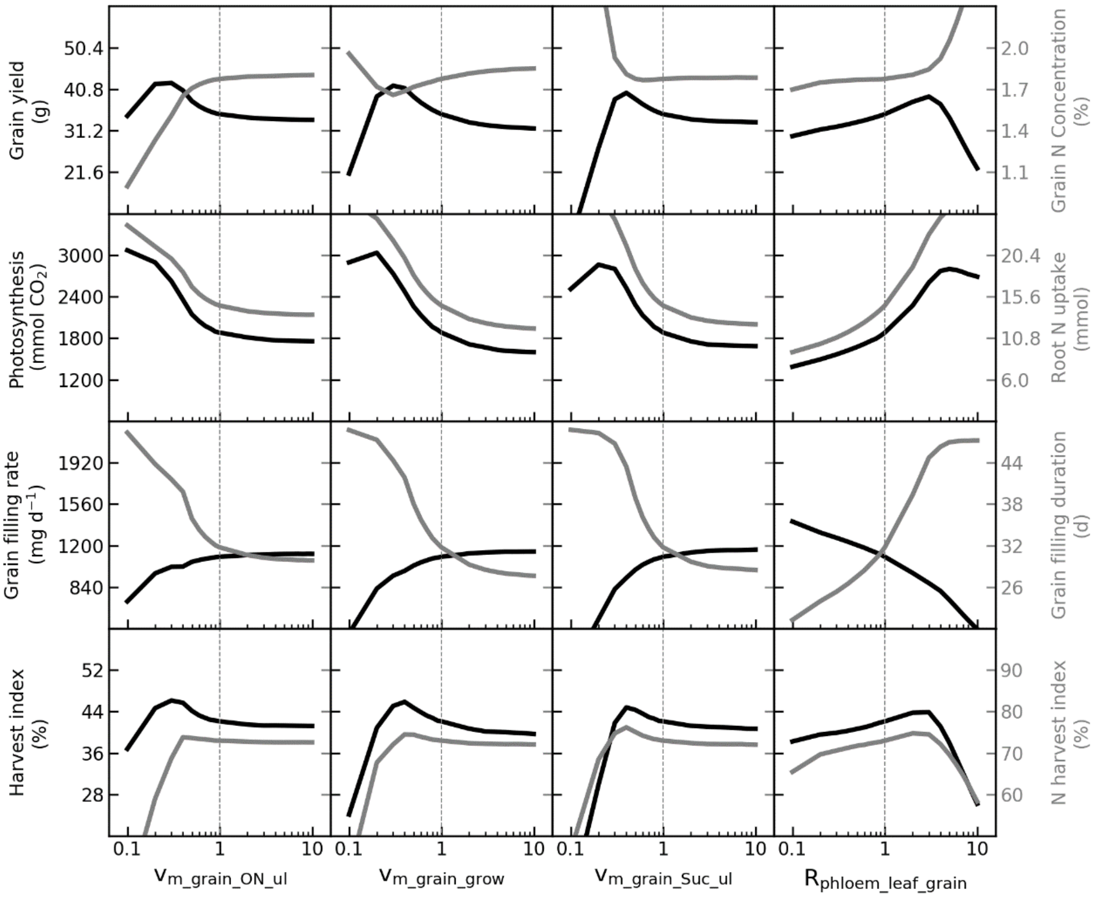


**Supplementary Figure 11. The responses of major agronomic traits to variation of model parameters $v_{m\_grain\_ON\_ul}$, $v_{m\_grain\_grow}$, $v_{m\_grain\_Suc\_ul}$, $R_{\_phloem\_leaf\_grain}$, respectively.**

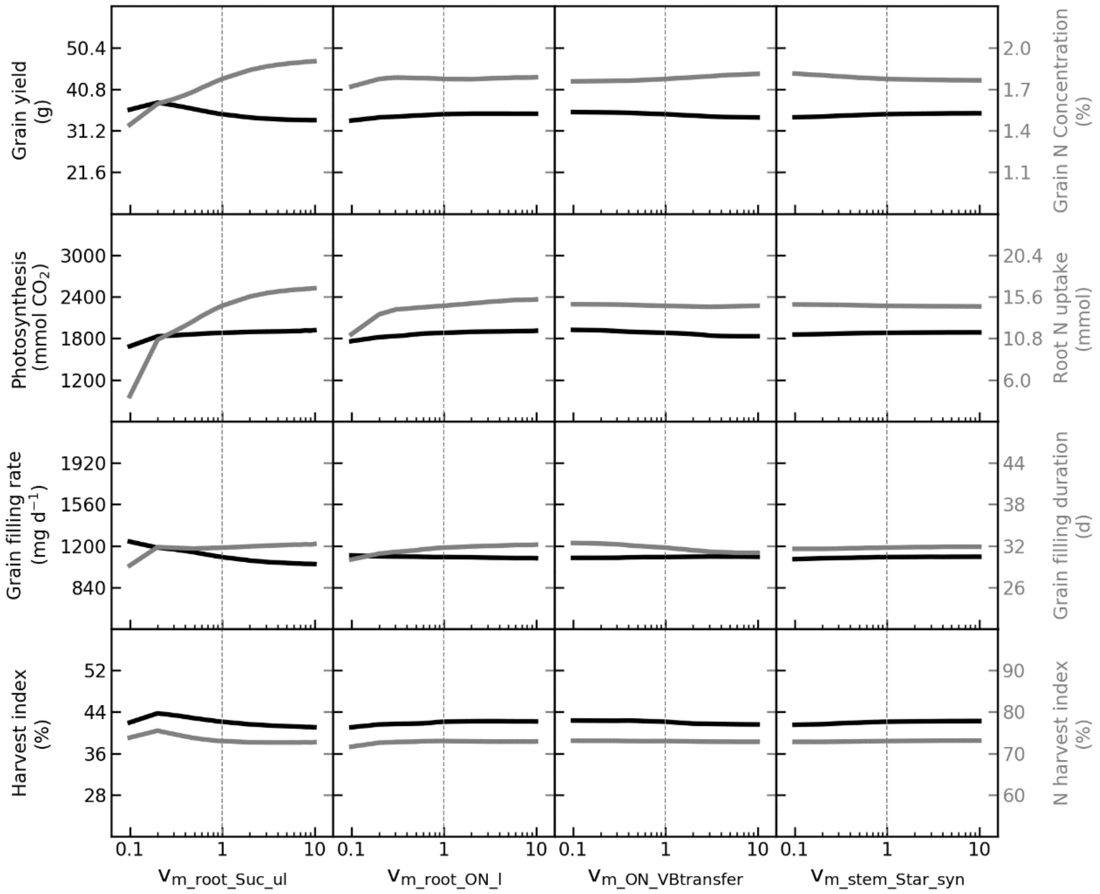


**Supplementary Figure 12. The responses of major agronomic traits to variation of model parameters $V_{m\_root\_Suc\_ul}$, $V_{m\_root\_ON\_l}$, $V_{m\_ON\_Vbtransfer}$, $V_{m\_stem\_Star\_syn}$, respectively.**

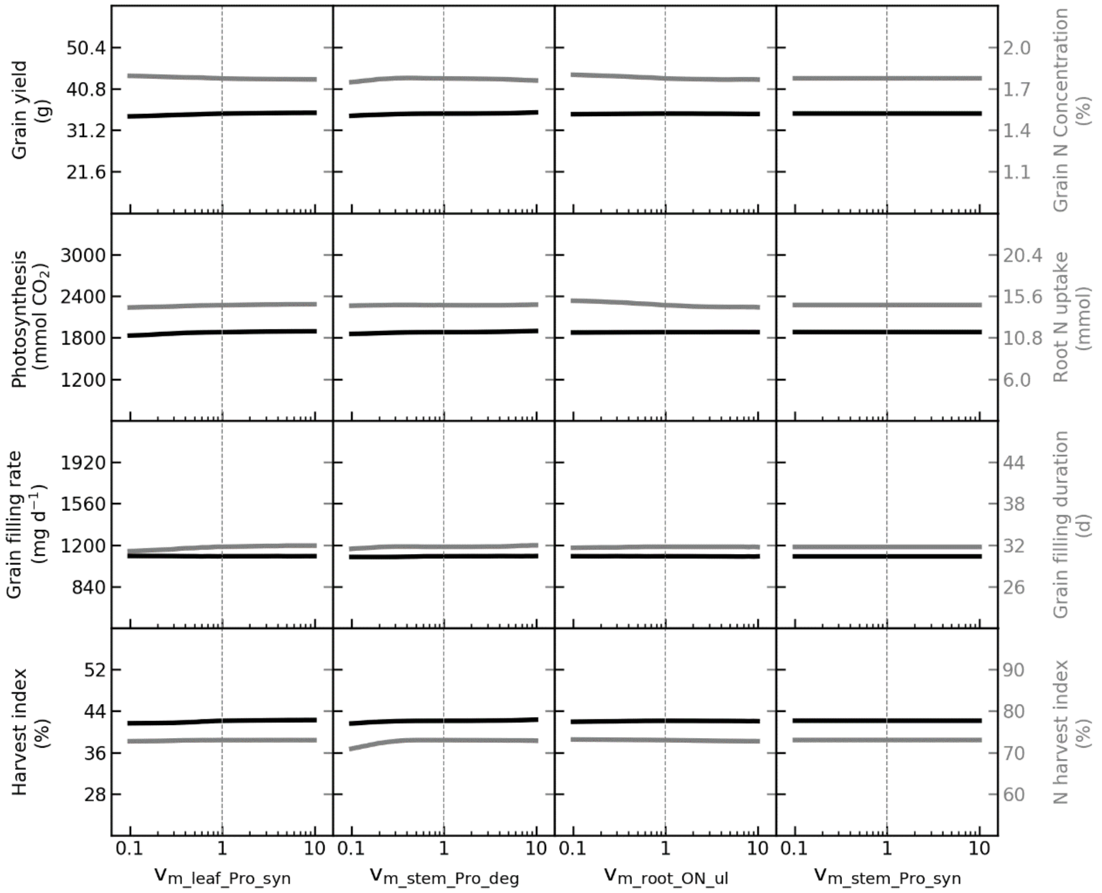


**Supplementary Figure 13. The responses of major agronomic traits to variation of model parameters $V_{m\_leaf\_Pro\_syn}$, $V_{m\_stem\_Pro\_deg}$, $V_{m\_root\_ON\_ul}$, $V_{m\_stem\_Pro\_syn}$, respectively.**

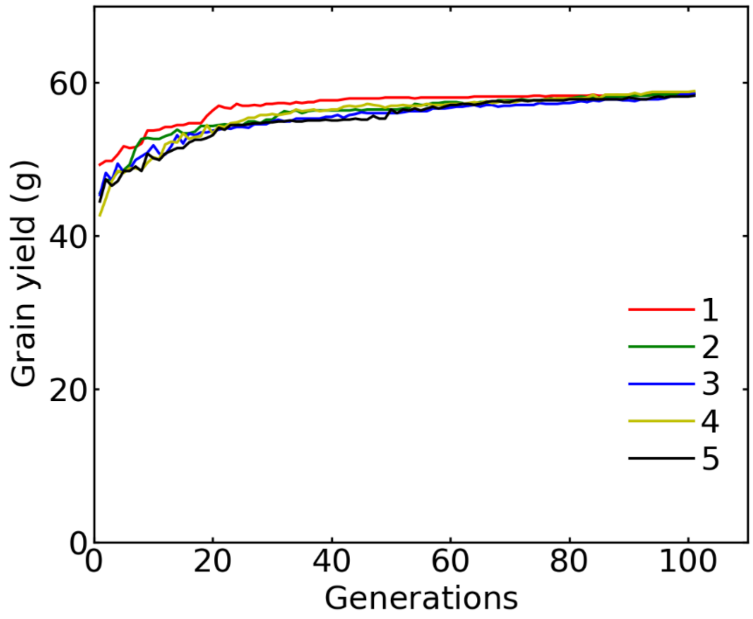


**Supplementary Figure 14. The evolution of total grain yield throughout generations in the five *in silico* "evolutionary populations".**

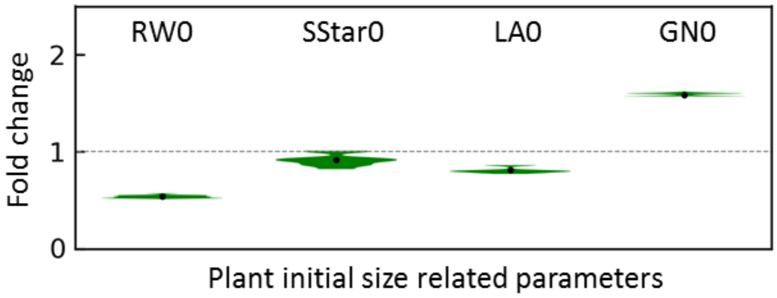


**Supplementary Figure 15. Distribution of relative value of plant initial size related parameters at flowering for *in silico* high-yield "Elite" individuals.** See parameter description in **Supplementary Table 2**. See parameter values for these individuals in **Supplementary Datasets 1**.

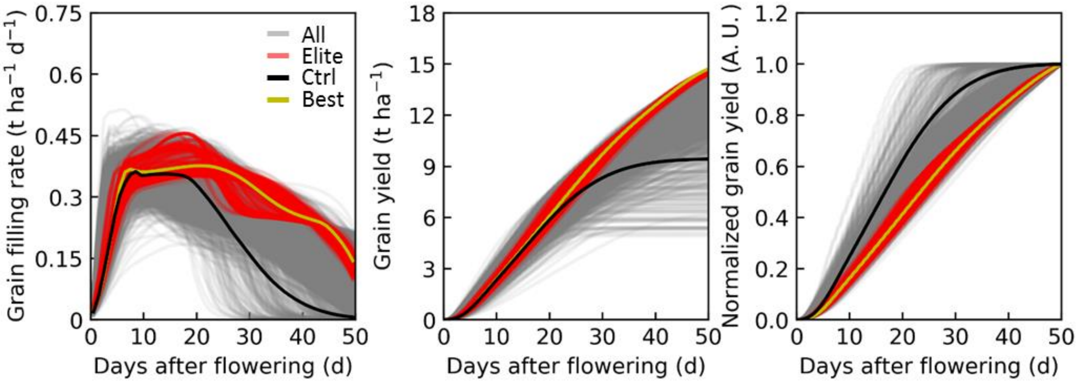


**Supplementary Figure 16. Grain filling patterns for plants in the *in silico* evolutionary populations.**

Abbreviations: “All”, all individuals in *in silico* evolutionary populations (n=50000); “Elite”, the top 1% individuals ranking by the final grain yield merged from five evolutionary populations (n=500); “Ctrl”, the individual with default parameters; “Best”, the most high-yield individual in *in silico* evolutionary populations.

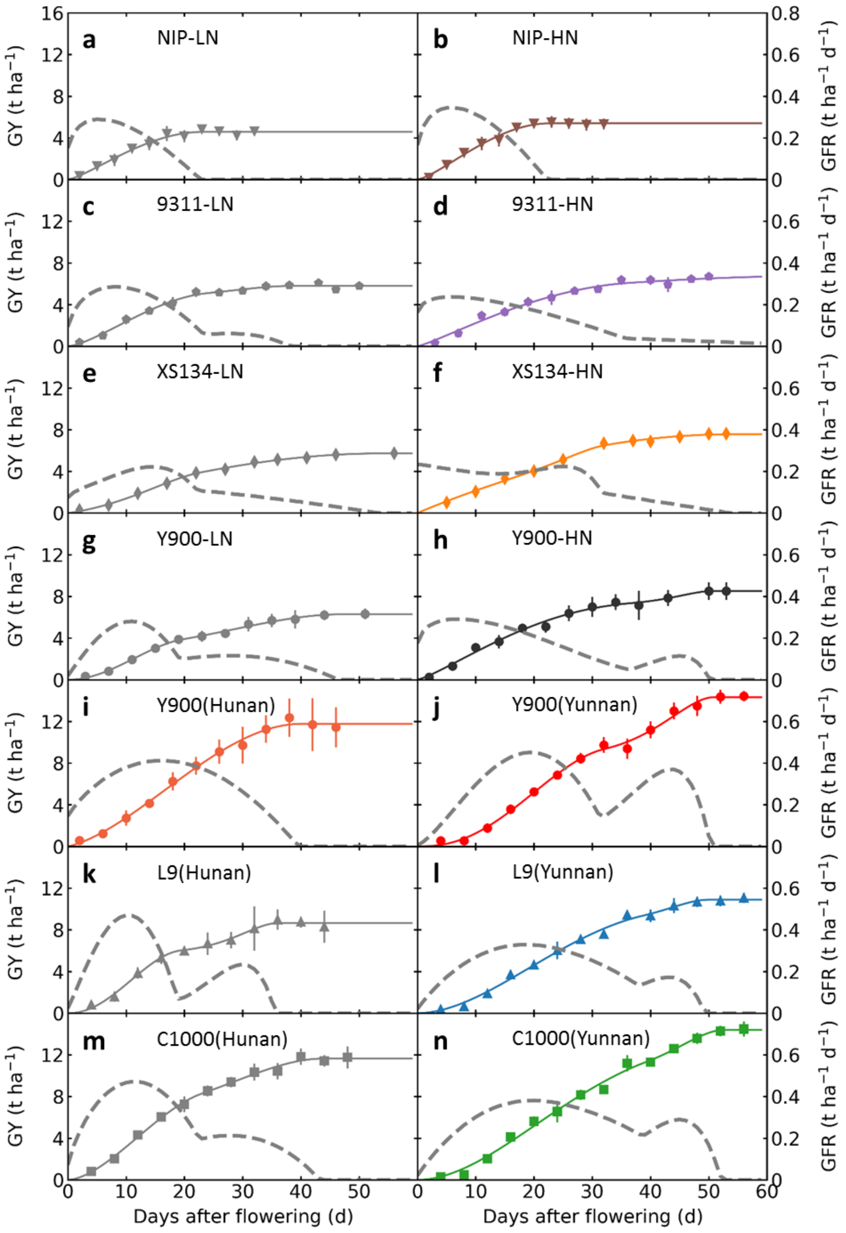


**Supplementary Figure 17. Grain filling pattern of different rice cultivars grown in different eco-zones.**

These experimental groups are: *Nipponbare* grown in Shanghai with low nitrogen (**a**) and high nitrogen (**b**); *9311* grown in Shanghai with low nitrogen (**c**) and high nitrogen (**d**); *Xiu-Shui 134* grown in Shanghai with low nitrogen (**e**) and high nitrogen (**f**); *Y-Liang-You 900* grown in Shanghai with low nitrogen (**g**) and high nitrogen (**h**); *Y-Liang-You 900* grown in Hunan (**i**) and in Yunnan (**j**); *Liang-You-Pei-Jiu* grown in Hunan (**k**) and in Yunnan (**l**); *Chao-You 1000* grown in Hunan (**m**) and in Yunnan (**n**).

The filled markers are measured data (mean ± sd, n=9) of grain yields with time after flowering; the solid lines are fitted values of grain yields with time after flowering; and the dashed lines are fitted values of grain filling rates with time after flowering.

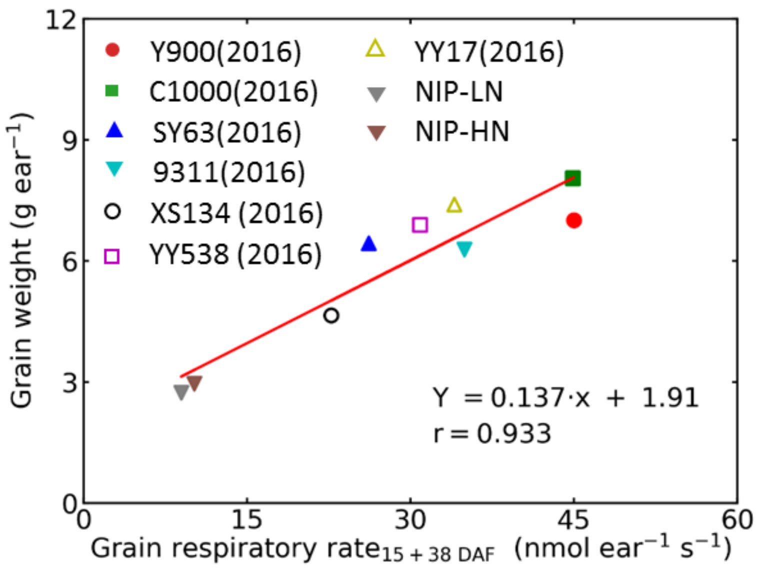


**Supplementary Figure 18. Relation between grain dry weight at harvest and a sum of grain respiratory rates at 15 and 38 days after flowering.**

See detailed information of the cultivars used in the **Methods**.

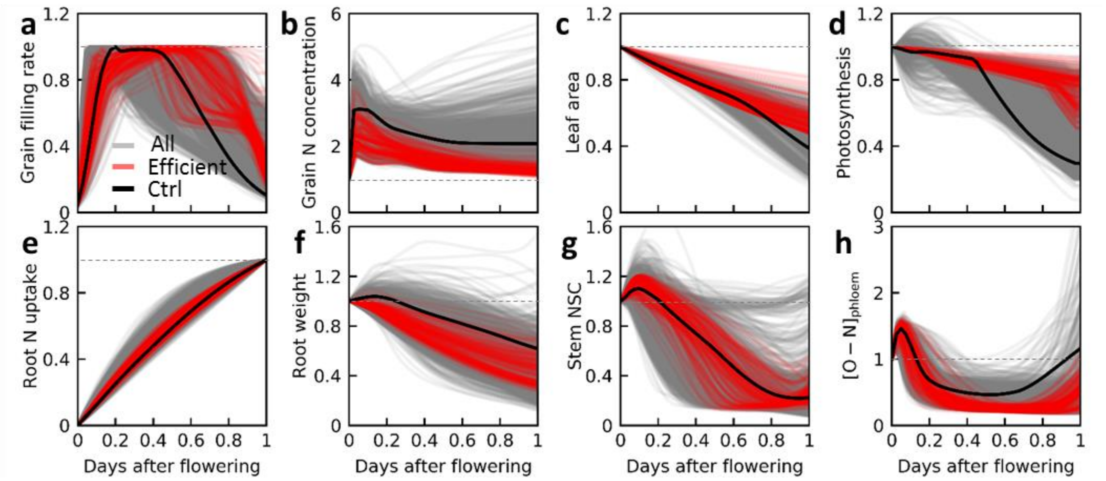


**Supplementary Figure 19. Comparison of macroscopic trait changes of the *in silico* rices.**

"Ctrl" is the individual with default model parameters, "Efficient" individuals have highest grain yields with different potential photosynthate accumulation, "All" are all individuals generated during *in silico* evolution.

Macroscopic traits include single-grain filling rate (**a**), grain nitrogen concentration (**b**), leaf area (**c**), photosynthetic rate (**d**), total accumulated root nitrogen uptake (**e**), root weight (**f**), amount of stem non-structural carbohydrates (NSC; **g**), and phloem organic nitrogen concentration ($[O\text{-}N]_{phloem}$; **h**). Notice that the harvesting time, maximum grain filling rate, initial grain nitrogen concentration, initial leaf area, initial photosynthetic rate, final accumulated root nitrogen uptake at harvest, initial root weight, initial stem non-structural carbohydrates, and initial phloem organic nitrogen concentration were normalized to 1.

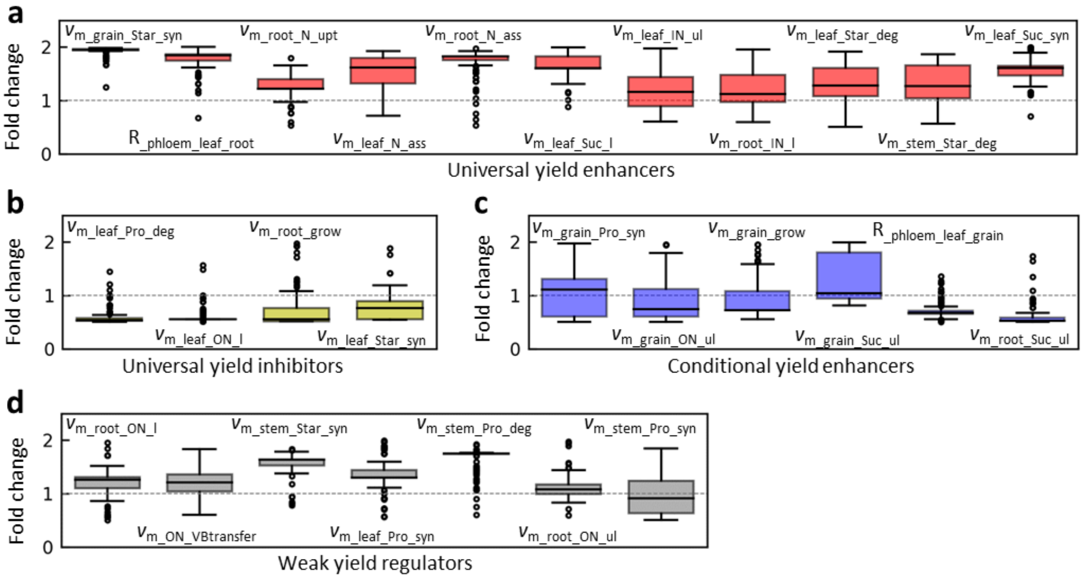


**Supplementary Figure 20. Distribution of relative value of each biochemical/physiological parameter for the *in silico* "Efficient" individuals.** The relative values of parameters refer to fold change from their default values, including universal yield enhancers (**a**), universal yield inhibitors (**b**), conditional yield enhancers (**c**) and weak yield regulators (**d**).

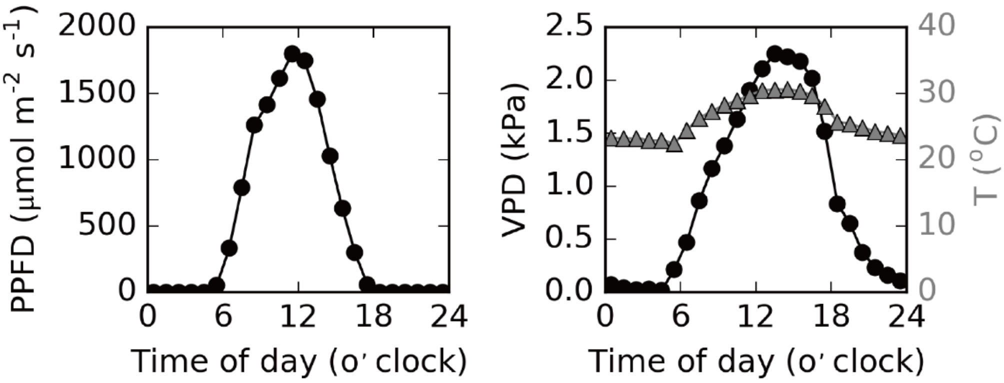


**Supplementary Figure 21. Daily weather data of the normal condition used in the model.**

Left, photosynthetic photon flux density (PPFD). Right, vapor pressure deficit (VPD, black line and circles) and temperature (T, grey line and triangles). As a simplification, we assumed that the other days during grain filling have the same diurnal PPFD, T and VPD pattern.

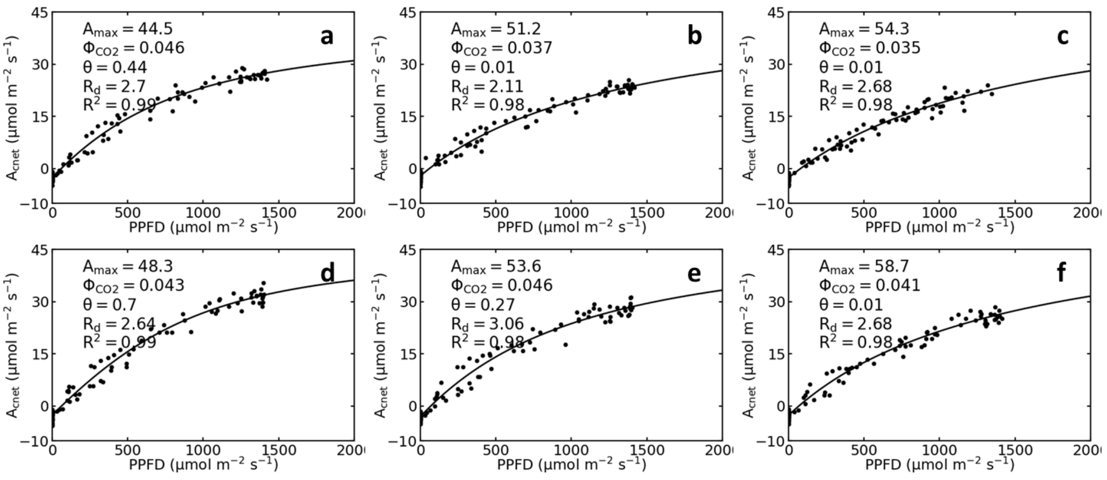


**Supplementary Figure 22. Canopy photosynthetic light response patterns at the flowering stage for rice cultivar *XS134* grown under low and high nitrogen conditions in Shanghai, 2015.**

(**a-c**) Three biological replicates for the low nitrogen condition.

(**d-e**) Three biological replicates for the high nitrogen condition.

**Supplementary Table 1. A list of the sub-models used in WACNI.**

| Module | Process | Sub-model annotation | Source |
|---|---|---|---|
| ***Root*** | I-N uptake | Root HATS and LATS | (Malagoli et al., 2004) |
| | | Feedback inhibitory effect from root N to N uptake | (Glass et al., 2002) |
| | | Promotive effect from root sugar to N uptake | (Monod, 1949); this study |
| | I-N assimilation | Sugar facilitated root N assimilation | This study |
| | Growth | Sugar facilitated root growth | (Radin et al., 1978); this study |
| | Senescence | Aging and carbon starvation driven root senescence | (Asseng et al., 1997); this study |
| ***Leaf*** | Photosynthesis | FvCB model of $C_3$ photosynthesis | (Von Caemmerer, 2000a) |
| | | Sun-shade model of canopy photosynthesis | (De Pury & Farquhar, 1997) |
| | | Stomatal conductance | (Leuning, 1995) |
| | | Promotive effect from protein to photosynthesis | (San-oh et al., 2006); this study |
| | | Feedback inhibitory effect from NSC to photosynthesis | (Azcón-Bieto, 1983); this study |
| | N assimilation | Photosynthetic N assimilation | (Walker et al., 2014); this study |
| | Sucrose and starch synthesis | Photosynthetic sucrose and starch synthesis | (Zhu et al., 2007); this study |
| | | Night leaf starch degradation | (Seaton et al., 2014); this study |
| | Protein synthesis and degradation | Leaf protein synthesis and degradation | This study |
| | Senescence | Aging and nitrogen starvation driven leaf senescence | (X. Yin & H. van Laar, 2005); this study |
| ***Grain*** | Volume expansion | Grain growth | This study |
| | Starch and protein storage | Grain starch and protein synthesis | This study |
| ***Stem*** | Sucrose and starch homeostasis | Stem starch synthesis and degradation | This study |
| | O-N and protein homeostasis | Stem protein synthesis and degradation | This study |
| ***Transport*** | Flux between different phloem compartments | Phloem long distance transport | (Lalonde et al., 2003; Minchin, Thorpe, & Farrar, 1993) |
| | Loading and unloading of metabolite | Metabolite transmembrane transport | (Y. Wang et al., 2014) |
| | Phloem-stem diffusion of metabolite | Metabolite diffusion | (Y. Wang et al., 2014) |
| ***Respiration*** | Organ respiration | Respiratory loss | (Cannell & Thornley, 2000; Thornley & Cannell, 2000) |

**Supplementary Table 2. The description and values of parameters used in WACNI.**

| Parameters | Units | Description | Range | Default | Source |
|---|---|---|---|---|---|
| ***Plant*** | | | | | |
| TillerNum | | Tiller number per plant | 2-31 | 12 | (Qu et al., 2017; Yoshida, 1981) |
| PlantDens | $m^{-2}$ | Planting density | 16-44 | 25 | (Baloch, Soomro, Javed, Ahmed, & Mastoi, 2002; Fukushima, Shiratsuchi, Yamaguchi, & Fukuda, 2011; Zheng et al., 2020) |
| ***Root*** | | | | | |
| RW0 | g | Root dry weight at flowering | 0.8-6.1 | 6 | (G. P. Chen et al., 2006; L. J. Liu et al., 2013; Tang et al., 2010; C. S. Zhang et al., 2005) |
| $c_{root\_strucN}$ | $g\ g^{-1}$ | Root structural N concentration | 0.0041-0.0054 | 0.005 | (H. Liu, 2009) |
| $C_{root\_FW2V}$ | $m^3\ g^{-1}$ | Conversion factor from root fresh weight to root aqueous space volume | - | $0.25*10^{-6}$ | Assumed |
| $C_{root\_FW2DW}$ | $g\ g^{-1}$ | Conversion factor from root fresh weight to root dry weight | 0.06-0.4 | 0.1 | D. Hong and Ichii (1996) |
| $[N]_{inhibit\_up}$ | $mol\ m^{-3}$ | Upper limit of total root N (i.e., I-N and O-N) concentration above which N uptake by HATS ceases | - | 100 | Assumed |
| $K_{m1}$ | $mol\ m^{-3}$ | Michaelis-constant | - | 10 | Assumed |
| $v_{m\_root\_HATS}$ | $mol\ g^{-1}DW\ s^{-1}$ | Maximal I-N uptake rate by root HATS | $2.4\text{-}61*10^{-9}$ | $3.65*10^{-9}$ | (L. Sun, Lu, Yu, Kronzucker, & Shi, 2016; M. Y. Wang, Siddiqi, Ruth, & Glass, 1993; Youngdahl, Pacheco, Street, & Vlek, 1982)[1] |
| $[I\text{-}N]_{soil}$ | $mol\ m^{-3}$ | Soil I-N concentration | 0-0.43 | 0.14 | (Yamakawa, Saigusa, Okada, & Kobayashi, 2004) |
| $K_{m2}$ | $mol\ m^{-3}$ | Michaelis-constant | 0.032-0.19 | 0.075 | (L. Sun et al., 2016; Teo, Beyrouty, & Gbur, 1992; M. Y. Wang et al., 1993; Youngdahl et al., 1982) |
| $k_{root\_LATS}$ | - | Slope of uptake rate versus N concentration for LATS | $4.7\text{-}10.7*10^{-10}$ | $5.5*10^{-10}$ | E.F. M. Y. Wang et al. (1993)[1] |
| $v_{m\_root\_N\_ass}$ | mol | Maximal I- | 15- | $1.25*10^{-}$ | Marwaha and |

| | | | | | |
|---|---|---|---|---|---|
| | $g^{-1}DW\ s^{-1}$ | N assimilation rate in root | $23*10^{-9}$ | 9 | Juliano (1976)[2] |
| $K_{m3}$ | $mol\ m^{-3}$ | Michaelis-constant | - | 10 | Set to be equal to $K_{m1}$ |
| $K_{m4}$ | $mol\ m^{-3}$ | Michaelis-constant | - | 20 | Assumed |
| $v_{m\_root\_grow}$ | $g\ g^{-1}\ d^{-1}$ | Root maximal relative growth rate | 0.13-0.28 | 0.2 | Tilman (1991) |
| $[Suc]_{grow\_low}$ | $mol\ m^{-3}$ | Lower limit of root sucrose concentration under which growth ceases | - | 10 | Assumed |
| $K_{m5}$ | $mol\ m^{-3}$ | Michaelis-constant | - | 40 | Assumed |
| $\alpha_{root\_sene}$ | $g\ g^{-1}\ d^{-1}$ | Root minimal senescence rate due to aging | - | 0.03 | Set to be $2*\alpha_{sene\_leaf}$[3] |
| $\beta_{root\_sene}$ | $m^3\ mol^{-1}$ | Coefficient in root senescence model | - | 0.24 | Assumed |
| $[Suc]_{sene\_up}$ | $mol\ m^{-3}$ | Upper limit of root sucrose concentration above which C starvation induced senescence relieves | - | 8 | Assumed |
| ***Leaf*** | | | | | |
| LA0 | $m^2$ | Plant total leaf area at flowering | 0.15-0.41 | 0.24 | (Haque, Pramanik, Biswas, Iftekharuddaula, & Hasanuzzaman, 2015; G. Li et al., 2009; Shi et al., 2019) |
| $Th_{leaf}$ | m | Leaf thickness | $1.1-4.2*10^{-4}$ | $1.75*10^{-4}$ | (Qu et al., 2017) |
| $C_{leaf\_FW2V}$ | $m^3\ g^{-1}$ | Conversion factor from leaf fresh weight to mesophyll aqueous space volume | - | $0.25*10^{-6}$ | Set to be equal to $C_{root\_FW2V}$ |
| SLA | $m^2\ g^{-1}$ | Specific leaf area | 0.019-0.037 | 0.02 | (G. Li et al., 2009; Xiong et al., 2016) |
| $c_{leaf\_strucN}$ | $g\ g^{-1}$ | Leaf structural N concentration | - | 0.005 | (De Pury & Farquhar, 1997) |
| LPro0 | $g\ N\ g^{-1}$ | Leaf protein content at flowering | 0.014-0.041 | 0.035 | (Zhao et al., 2015)[4] |
| $[Pro]_{promote\_up}$ | $mol\ N\ m^{-3}$ | Upper limit of leaf mesophyll aqueous space protein concentration above which photosynthesis | $2.2-2.8*10^3$ | $2.2*10^3$ | (Shi et al., 2019)[5] |

| | | | | | |
|---|---|---|---|---|---|
| | | will not be further promoted | | | |
| $[NSC]_{inhibit\_low}$ | mol Suc $m^{-3}$ | Lower limit of carbohydrates concentration under which photosynthesis will not be further inhibited | - | 400 | Assumed |
| $[NSC]_{inhibit\_up}$ | mol Suc $m^{-3}$ | Upper limit of carbohydrates concentration above which photosynthesis will be completely blocked | - | 1900 | Assumed |
| $v_{cmax0}$ | mol C $m^{-2}$ $s^{-1}$ | Maximal Rubisco carboxylation rate | 80-200*$10^{-6}$ | 80*$10^{-6}$ | (Chang et al., 2017; Gu, Yin, Stomph, & Struik, 2014; Von Caemmerer, 2000a) |
| $\Gamma$ | μbar | $CO_2$ compensation point in the absence of mitochondrial respiration | - | 38.6 | (Von Caemmerer, 2000a) |
| $k_{b0}$ | - | Canopy beam radiation extinction coefficient | 0.27-0.81 | 0.37 | Kiniry, McCauley, Xie, and Arnold (2001) |
| $K_{m6}$ | μbar | Michaelis-constant | 418 | 418 | (Von Caemmerer, 2000a) |
| $J_{max0}$ | mol $e^{-}$ $m^{-2}$ $s^{-1}$ | Maximal linear photosynthetic electron transport rate | 137-227*$10^{-6}$ | 160*$10^{-6}$ | (Chang et al., 2017; Gu et al., 2014; Von Caemmerer, 2000a) |
| $\varphi_0$ | - | Proportion of light absorbed by PS II of total incident light | 0.272-0.369 | 0.3612 | (Gu et al., 2014; Von Caemmerer, 2000a) |
| $\theta_0$ | - | Empirical curvature factor | 0.15-0.916 | 0.7 | (Gu et al., 2014; Shi et al., 2019) |
| $f_a$ | - | Fraction of diffuse irradiance | - | 0.425 | Pury and Farquhar (1997) |
| $a$ | - | Atmospheric transmission coefficient of PAR | - | 0.75 | Pury and Farquhar (1997) |
| $k_d$ | - | Diffuse PAR extinction coefficient | - | 0.78 | Pury and Farquhar (1997) |
| $\rho_{cb}$ | - | Canopy reflection coefficient for beam PAR | - | 0.029 | Pury and Farquhar (1997) |
| $\rho_{cd}$ | - | Canopy reflection coefficient for diffuse PAR | - | 0.036 | Pury and Farquhar (1997) |

| | | | | | |
|---|---|---|---|---|---|
| $\sigma$ | - | Leaf scattering coefficient of PAR | - | 0.15 | Pury and Farquhar (1997) |
| $[CO_2]_a$ | μbar | Ambient $CO_2$ concentration | 380-420 | 400 | Current atmosphere status |
| $g_0$ | mol $m^{-2}$ $s^{-1}$ | Residual stomatal conductance | - | 0.01 | Leuning (1995) |
| $a_1$ | - | Empirical correction factor | - | 20 | Leuning (1995) |
| $VPD_0$ | kPa | Empirical correction factor | - | 0.35 | Leuning (1995) |
| $v_{m\_leaf\_N\_ass}$ | mol $m^{-2}$ $s^{-1}$ | Maximal I-N assimilation rate | $0.9\text{-}7.3*10^{-7}$ | $6.2*10^{-7}$ | (Marwaha & Juliano, 1976; Walker et al., 2014)[6] |
| $K_{m7}$ | mol $m^{-3}$ | Michaelis-constant | - | 150 | Assumed |
| $v_{m\_leaf\_Suc\_syn}$ | mol C12 $m^{-2}$ $s^{-1}$ | Maximal leaf sucrose synthesis rate | $3.3\text{-}5.3*10^{-6}$ | $3.3*10^{-6}$ | Gesch, Vu, Boote, Allen, and Bowes (2002)[7] |
| $K_{e1}$ | - | Substrates feedback regulatory factor | - | 300 | Assumed |
| $K_{m8}$ | mol $m^{-3}$ | Michaelis-constant | - | 2 | Assumed |
| $v_{m\_leaf\_Star\_syn}$ | mol C12 $m^{-2}$ $s^{-1}$ | Maximal TP-to-starch conversion rate | $4.7\text{-}15.4*10^{-7}$ | $8.3*10^{-7}$ | Okamura et al. (2013)[8] |
| $K_{m9}$ | mol $m^{-3}$ | Michaelis-constant | - | 2 | Assumed |
| $v_{m\_leaf\_Star\_deg}$ | mol C12 $m^{-2}$ $s^{-1}$ | Maximal leaf night starch-to-sucrose conversion rate | - | $8.3*10^{-6}$ | Set to be 10* $v_{m\_leaf\_Star\_syn}$ |
| $[Suc]_{leaf\_Star\_deg\_low}$ | mol $m^{-3}$ | Lower limit of night leaf sucrose concentration below which starch decomposes | - | 120 | Assumed |
| $K_{m10}$ | mol $m^{-3}$ | Michaelis-constant | - | 15 | Assumed |
| $v_{m\_leaf\_Pro\_syn}$ | mol $m^{-2}$ $s^{-1}$ | Maximal leaf protein synthesis rate | - | $6.2*10^{-7}$ | Set to be equal to $v_{m\_N\_ass\_leaf}$ |
| $[O\text{-}N]_{leaf\_Pro\_syn\_low}$ | mol $m^{-3}$ | Lower limit of leaf O-N concentration below which protein synthesis ceases and starts to decompose | - | 100 | Assumed |
| $K_{m11}$ | mol $m^{-3}$ | Michaelis-constant | - | 45 | Assumed |
| $k_{leaf\_Pro\_turno}$ | $day^{-1}$ | Leaf protein turnover rate | - | 0.1 | Yoshida (1981) |
| $v_{m\_leaf\_Pro\_deg}$ | mol $m^{-2}$ $s^{-1}$ | Maximal leaf protein decomposition | - | $9.2*10^{-8}$ | Assumed |

| | | | | | |
|---|---|---|---|---|---|
| | | rate | | | |
| $\alpha_{leaf\_sene}$ | $m^2\ m^{-2}\ d^{-1}$ | Leaf minimal senescence rate due to aging | - | 0.015 | X. Yin and H. H. van Laar (2005)[9] |
| $\beta_{leaf\_sene}$ | - | Constant coefficient | - | 0.0015 | Assumed |
| $[N]_{sene\_up}$ | $mol\ N\ m^{-3}$ | Upper limit of total leaf N (i.e., I-N, O-N and protein) concentration above which N starvation induced senescence relieves | - | 2000 | Assumed |
| ***Grain*** | | | | | |
| GN0 | - | Total grain number per plant | 586-1927 | 1920 | (L. J. Liu et al., 2013; Smidansky et al., 2003; E. Wang et al., 2008) |
| GSR | - | Grain setting rate, i.e., filled grains to total grains ratio at harvest | 0.75-0.96 | 0.875 | (L. J. Liu et al., 2013; J. Yang & Zhang, 2010) |
| $c_{grain\_strucN}$ | $g\ g^{-1}$ | Grain structural N concentration | - | 0.04 | Assumed |
| $Wid_{grain}$ | m | Grain width | $1.2\text{-}3.8*10^{-3}$ | $2.4*10^{-3}$ | (Fitzgerald, McCouch, & Hall, 2009; X. Huang et al., 2010) |
| $Th_{grain}$ | m | Grain thickness | $1.5\text{-}2.3*10^{-3}$ | $2.4*10^{-3}$ | (C. Fan et al., 2006) |
| $LWR_{grain}$ | - | Grain length width ratio | 2-3.96 | 2.8 | (C. Fan et al., 2006; Mottaleb & Mishra, 2016; S. Wang et al., 2012) |
| $V_{glume}$ | $m^3$ | Glume volume for one spikelet | - | $2.03*10^{-8}$ | Set to be $4\pi/3*(Wid_{grain}*LWR_{grain}/2)*(Wid_{grain}/2)*(Th_{grain}/2)$ [10] |
| HW | g | Husk dry weight for one spikelet | - | $5.1*10^{-3}$ | Set to be $V_{glume}/0.8*0.2*10^6$ [11] |
| $c_{husk\_strucN}$ | $g\ g^{-1}$ | Husk structural N concentration | - | 0.005 | Assumed |
| $\varepsilon$ | - | Empirical coefficient, for which $(1-\varepsilon)\cdot V_{glume}$ is a upper limit of grain volume above which grain growth will be more and more physically restrained | - | 0.3 | Assumed |
| $v_{m\_grain\_grow}$ | $m^3\ m^{-2}\ s^{-1}$ | Maximal grain volume expansion rate | - | $4*10^{-9}$ | E.F. J. Yang, Zhang, Wang, Liu, and Wang (2006)[12] |
| $K_{m12}$ | mol | Michaelis- | - | 50 | Assumed |

| | | | | | |
|---|---|---|---|---|---|
| | $m^{-3}$ | constant | | | |
| $K_{m13}$ | mol $m^{-3}$ | Michaelis-constant | - | 80 | Assumed |
| $\rho_{St}$ | kg $m^{-3}$ | Density of grain storage starch | - | $1*10^{3}$ | Simplification |
| $\rho_{Pro}$ | kg $m^{-3}$ | Density of grain storage protein | - | $1*10^{3}$ | Simplification |
| $v_{m\_grain\_Star\_syn}$ | mol C12 $m^{-3}$ $s^{-1}$ | Maximal grain starch synthesis rate | $0.63\text{-}25*10^{-3}$ | $9.5*10^{-3}$ | (S. Chen, Zhou, Zeng, & Zhang, 2011; Devi, Sarla, Siddiq, & Sirdeshmukh, 2010; Perez, Perdon, Resurreccion, Villareal, & Juliano, 1975)[13] |
| $K_{m14}$ | mol $m^{-3}$ | Michaelis-constant | - | 160 | Assumed |
| $v_{m\_grain\_Pro\_syn}$ | mol N $m^{-3}$ $s^{-1}$ | Maximal grain protein synthesis rate | $0.05\text{-}5.7*10^{-3}$ | $3.4*10^{-3}$ | E.F. (Carlile, Raviv, & Prasad, 2019; Martin & Fitzgerald, 2002; Peng et al., 2014; Y. Yang et al., 2019)[14] |
| $K_{m15}$ | mol $m^{-3}$ | Michaelis-constant | - | 100 | Assumed |
| ***Stem*** | | | | | |
| SSW0 | g | Stem structural dry weight | 14.2-38.9 | 36 | (K. Liu et al., 2020; J. Yang, Zhang, Wang, Zhu, & Wang, 2001)[15] |
| SV0 | $m^{3}$ | Total stem aqueous space volume | $0.85\text{-}2.33*10^{-5}$ | $2.16*10^{-5}$ | E.F. SSW0[16] |
| SStar0 | g | Stored starch content at flowering | 0.8-14.6 | 9 | E.F. SSW0 and J. Yang et al. (2001)[17] |
| SPro0 | g | Stored protein content at flowering | - | 0.24 | Assumed |
| $c_{stem\_strucN}$ | g $g^{-1}$ | Stem structural N concentration | - | 0.002 | Assumed |
| $v_{m\_stem\_Star\_syn}$ | mol C12 $m^{-3}$ $s^{-1}$ | Maximal stem starch synthesis rate | $0.21\text{-}1.67*10^{-2}$ | $1.25*10^{-2}$ | E.F. Okamura et al. (2013)[18] |
| $v_{m\_stem\_Star\_deg}$ | mol C12 $m^{-3}$ $s^{-1}$ | Maximal stem starch degradation rate | - | $1*10^{-2}$ | Set to be close to $v_{m\_stem\_Star\_syn}$ |
| $[Suc]_{stem\_Star\_syn\_low}$ | mol $m^{-3}$ | Lower limit of stem sucrose concentration below which starch synthesis ceases and starts to decompose | - | 500 | Assumed |
| $K_{m16}$ | mol $m^{-3}$ | Michaelis-constant | - | 160 | Set to be equal to $K_{m14}$ |
| $K_{m17}$ | mol $m^{-3}$ | Michaelis-constant | - | 600 | Assumed |
| $v_{m\_stem\_Pro\_syn}$ | mol $m^{-3}$ $s^{-1}$ | Maximal stem protein synthesis rate | | $3.4*10^{-3}$ | Set to be equal to $v_{m\_grain\_Pro\_syn}$ |
| $v_{m\_stem\_Pro\_deg}$ | mol | Maximal | - | $2.1*10^{-3}$ | Set to be equal to |

| | | | | | |
|---|---|---|---|---|---|
| | $m^{-3}\ s^{-1}$ | stem protein decomposition rate | | | $v_{m\_leaf\_Pro\_deg}$ |
| $[O\text{-}N]_{stem\_Pro\_syn\_low}$ | mol $m^{-3}$ | Lower limit of stem O-N concentration below which protein synthesis ceases and starts to decompose | - | 300 | Assumed |
| $K_{m18}$ | mol $m^{-3}$ | Michaelis-constant | - | 200 | Assumed |
| $K_{m19}$ | mol $m^{-3}$ | Michaelis-constant | - | 200 | Assumed |
| ***Xylem and phloem transport*** | | | | | |
| XV0 | $m^3$ | Xylem volume | 0.085-10.6*$10^{-6}$ | 1.2*$10^{-6}$ | E.F. (H. Huang, 1998; B. H. Zhao, P. Wang, H. Zhang, Q. S. Zhu, & J. C. Yang, 2006)[19] |
| $PV0_{\_leaf}$ | $m^3$ | Volume of phloem at leaf terminal | - | 4*$10^{-7}$ | E.F. (H. Huang, 1998; B. H. Zhao et al., 2006)[19] |
| $PV0_{\_root}$ | $m^3$ | Volume of phloem at root terminal | - | 4*$10^{-7}$ | Assumed to be equal to $PV0_{\_leaf}$ |
| $PV0_{\_grain}$ | $m^3$ | Volume of phloem at grain terminal | - | 4*$10^{-7}$ | Assumed to be equal to $PV0_{\_leaf}$ |
| $R_{\_phloem\_leaf\_root}$ | mol s $m^{-6}$ | Phloem transport resistance between leaf and root | 0.5-10*$10^{13}$ | 7*$10^{13}$ | Minchin et al. (1993) |
| $R_{\_phloem\_leaf\_grain}$ | mol s $m^{-6}$ | Phloem transport resistance between leaf and grain | 0.5-10*$10^{13}$ | 1.75*$10^{13}$ | Set to be 1/4* $R_{\_phloem\_leaf\_root}$ |
| *Root (un)loading* | | | | | |
| $v_{m\_root\_IN\_l}$ | mol $g^{-1}DW\ s^{-1}$ | Root maximal I-N loading rate | - | 2.5*$10^{-9}$ | Set to be $v_{m\_root\_HATS}$ - $v_{m\_root\_N\_ass}$ |
| $K_{e\_root\_IN\_l}$ | - | Substrates feedback regulatory factor | - | 1 | Assumed |
| $K_{m\_root\_IN\_l}$ | mol $m^{-3}$ | Michaelis-constant | - | 10 | Assumed |
| $v_{m\_root\_ON\_l}$ | mol $g^{-1}DW\ s^{-1}$ | Root maximal O-N loading rate | - | 3.2*$10^{-9}$ | Set to be equal to $v_{m\_root\_ON\_ul}$ |
| $K_{e\_root\_ON\_l}$ | - | Substrates feedback regulatory factor | - | 1 | Assumed |
| $K_{m\_root\_ON\_l}$ | mol $m^{-3}$ | Michaelis-constant | - | 10 | Assumed |
| $v_{m\_root\_ON\_ul}$ | mol $g^{-1}DW\ s^{-1}$ | Root maximal O-N unloading rate | - | 3.2*$10^{-9}$ | Set to be 0.32* $v_{m\_root\_Suc\_ul}$ [20] |
| $K_{e\_root\_ON\_ul}$ | - | Substrates | - | 0.1 | Assumed |

| | | | | | |
|---|---|---|---|---|---|
| | | feedback regulatory factor | | | |
| $K_{m\_root\_ON\_ul}$ | mol m$^{-3}$ | Michaelis-constant | - | 100 | Assumed |
| $v_{m\_root\_Suc\_ul}$ | mol g$^{-1}$DW s$^{-1}$ | Root maximal sucrose unloading rate | 0.58-2.4*10$^{-8}$ | 1*10$^{-8}$ | E.F. Tilman (1991), Farrar and Williams (1991)[21] |
| $K_{e\_root\_Suc\_ul}$ | - | Substrates feedback regulatory factor | - | 0.1 | Assumed |
| $K_{m\_root\_Suc\_ul}$ | mol m$^{-3}$ | Michaelis-constant | - | 300 | Assumed |
| *Leaf (un)loading* | | | | | |
| $v_{m\_leaf\_IN\_ul}$ | mol m$^{-2}$ s$^{-1}$ | Leaf maximal I-N unloading rate | - | 6*10$^{-8}$ | E.F. $v_{m\_root\_IN\_l}$ [22] |
| $K_{e\_leaf\_IN\_ul}$ | - | Substrates feedback regulatory factor | - | 10 | Assumed |
| $K_{m\_leaf\_IN\_ul}$ | mol m$^{-3}$ | Michaelis-constant | - | 10 | E.F. Y. G. Li et al. (2015) |
| $v_{m\_leaf\_ON\_ul}$ | mol m$^{-2}$ s$^{-1}$ | Leaf maximal O-N unloading rate | - | 7.5*10$^{-8}$ | E.F. $v_{m\_root\_ON\_l}$ [23] |
| $K_{e\_leaf\_ON\_ul}$ | - | Substrates feedback regulatory factor | - | 15 | Assumed |
| $K_{m\_leaf\_ON\_ul}$ | mol m$^{-3}$ | Michaelis-constant | - | 10 | Assumed |
| $v_{m\_leaf\_ON\_l}$ | mol m$^{-2}$ s$^{-1}$ | Leaf maximal O-N loading rate | - | 4.8*10$^{-7}$ | Set to be 0.4* $v_{m\_leaf\_Suc\_l}$ [20] |
| $K_{e\_leaf\_ON\_l}$ | - | Substrates feedback regulatory factor | - | 2 | Assumed |
| $K_{m\_leaf\_ON\_l}$ | mol m$^{-3}$ | Michaelis-constant | - | 50 | Assumed |
| $v_{m\_leaf\_Suc\_l}$ | mol m$^{-2}$ s$^{-1}$ | Leaf maximal sucrose loading rate | - | 1.2*10$^{-6}$ | E.F. (Eom et al., 2011; L. Wang, Lu, Wen, & Lu, 2015) [24] |
| $K_{e\_leaf\_Suc\_l}$ | - | Substrates feedback regulatory factor | - | 3 | Assumed |
| $K_{m\_leaf\_Suc\_l}$ | mol m$^{-3}$ | Michaelis-constant | - | 200 | Assumed |
| *Grain unloading* | | | | | |
| $v_{m\_grain\_ON\_ul}$ | mol m$^{-2}$ s$^{-1}$ | Grain maximal O-N unloads (from phloem) rate based on grain surface area | - | 2.0*10$^{-6}$ | Set to be 0.4* $v_{m\_grain\_Suc\_ul}$ [20] |
| $K_{e\_grain\_ON\_ul}$ | - | Substrates feedback regulatory factor | - | 1 | Assumed |
| $K_{m\_grain\_ON\_ul}$ | mol m$^{-3}$ | Michaelis-constant | - | 100 | Assumed |
| $v_{m\_grain\_Suc\_ul}$ | mol m$^{-2}$ s$^{-1}$ | Grain maximal sucrose unloads (from phloem) rate based on grain surface | - | 5.0*10$^{-6}$ | E.F. N. Wang and Fisher (1994)[25] |

| | | | | | |
|---|---|---|---|---|---|
| | | area | | | |
| $K_{e\_grain\_Suc\_ul}$ | - | Substrates feedback regulatory factor | - | 1 | Assumed |
| $K_{m\_grain\_Suc\_ul}$ | mol m$^{-3}$ | Michaelis-constant | - | 400 | Assumed |
| *Phloem-stem diffusion* | | | | | |
| $R_{\_phloem\_stem\_Suc}$ | s m$^{-3}$ | Phloem-to-stem sucrose diffusion resistance | - | 7.4*10$^{7}$ | E.F. Y. Wang et al. (2014)[26] |
| $R_{\_phloem\_stem\_ON}$ | s m$^{-3}$ | Phloem-to-stem sucrose diffusion resistance | - | 3.2*10$^{7}$ | E.F. Y. Wang et al. (2014)[26] |
| *Xylem-phloem transfer* | | | | | |
| $v_{m\_ON\_VBtransfer}$ | mol s$^{-1}$ | Maximal xylem-to-phloem O-N transfer rate | - | 1.2*10$^{-8}$ | E.F. $v_{m\_root\_IN\_l}$ [27] |
| $K_{e\_ xylem\_ON\_l}$ | - | Substrates feedback regulatory factor | - | 45 | Assumed |
| $K_{m\_ xylem\_ON\_l}$ | mol m$^{-3}$ | Michaelis-constant | - | 10 | Assumed |
| ***Respiration*** | | | | | |
| $\eta_{load}$ | mol mol$^{-1}$ | Sucrose equiv. C consumed per mole material loaded to phloem/xylem | - | 0.06 | Cannell and Thornley (2000) |
| $\eta_{unload}$ | mol mol$^{-1}$ | Sucrose equiv. C consumed per mole material unloaded from xylem/phloem | - | 0.06 | Set to be equal to $\eta_{load}$ |
| $\eta_{N\_abs}$ | mol mol$^{-1}$ | Sucrose equiv. C consumed per mole I-N uptake from soil (average value of $NH_4^+$ and $NO_3^-$ uptake) | - | 0.025 | Cannell and Thornley (2000) |
| $\eta_{N\_ass}$ | mol mol$^{-1}$ | Sucrose equiv. C consumed per mole I-N assimilated in dark (average value of $NH_4^+$ and $NO_3^-$ assimilation) | - | 0.11 | Cannell and Thornley (2000) |
| $\eta_{grow}$ | g g$^{-1}$ | Sucrose consumed for dry matter production per gram sucrose | - | 0.23 | Cannell and Thornley (2000) |
| $\eta_{store}$ | mol mol$^{-1}$ | Sucrose equiv. C consumed | 0.017-0.05 | 0.03 | E.F. Plaxton (1996)[28] |

| | | | | | |
|---|---|---|---|---|---|
| | | between sucrose and starch conversion | | | |
| $\eta_{Pro\_decomp}$ | mol mol$^{-1}$ N | Sucrose equiv. C consumed per mole N equiv. protein decomposed | - | 0.05 | De Visser, Spitters, and Bouma (1992) |
| $\eta_{Pro\_syn}$ | mol mol$^{-1}$ N | Sucrose equiv. C consumed per mole N equiv. protein synthesized | 0.3 | 0.3 | De Visser et al. (1992) |
| $\gamma_{residual}$ | mol C12 m$^{-3}$ s$^{-1}$ | Other maintenance process costs rate of leaf, stem and grains, based on aqueous space volume of organ | - | $6.9*10^{-5}$ | Assumed |
| $\gamma_{residual\_root}$ | mol C12 m$^{-3}$ s$^{-1}$ | Other maintenance process costs rate of root, based on aqueous space volume of root | - | $1.39*10^{-4}$ | Set to be 2*$\gamma_{residual}$ |

Note:

[1] A lower value of this parameter was used as during grain filling period, root nutrient uptake was not as active as that in the tillering stage (Yoshida, 1981).

[2] It has been shown in Marwaha and Juliano (1976) that for rice seedlings, *in vitro* nitrate reductase activity in root is in trace amount, which means almost all the nitrate is assimilated in shoot (~100%); in 10-day-old seedlings under hypodronic condition, glutamine synthetase activity in root is 0.37-0.56 μmol min$^{-1}$ g$^{-1}$ fresh weight, which equals to 15-23*10$^{-9}$ mol g$^{-1}$DW s$^{-1}$ with assuming root fresh weight to dry weight ratio of 2.5 (D. Hong & Ichii, 1996). Here we further assumed a much lower assimilation activity in root of after-flowering mature plants grown in the paddy field.

[3] It has been shown that natural root senescence rate is faster than that of a leaf (D. Wei, Ning, & Lin, 2004).

[4] It can be calculated from Zhao et al. (2015) that leaf protein-derived amino acid concentration is 0.088-0.13 g g$^{-1}$, which equals to nitrogen concentration of 0.014-0.041 g N g$^{-1}$ with a nitrogen-to-protein conversion factor ranging from 3.28 to 6.25 (Yeoh & Wee, 1994).

[5] It can be estimated from Shi et al. (2019) that rice leaf photosynthetic rate does not increase further when total leaf N (TN) exceeds 1.9 g N m$^{-2}$. Leaf structural N (SN) is estimated to be 0.15-0.3 g N m$^{-2}$; leaf free amino acids (AN) concentration is 0.007-0.030 g N m$^{-2}$ (Zhao et al., 2015) with a nitrogen-to-protein conversion factor ranging

from 3.28 to 6.25 (Yeoh & Wee, 1994); leaf inorganic N (IN) concentration in barley is 1.7-6.8 folds of amino acids (Winter, Robinson, & Heldt, 1993), which equals to 0.012-0.2 g N $m^{-2}$. Therefore, $Pro]_{promote\_up}$=2.2-2.8*$10^3$ mol $m^{-3}$ [(TN-SN-AN-IN)/14/($Th_{leaf}$ * $C_{leaf\_FW2V}$)].

[6] It has been shown in Marwaha and Juliano (1976) that for rice seedlings, *in vitro* nitrate reductase activity in leaf is 31-251 nmol $min^{-1}$ $g^{-1}$ fresh weight, which equals to 0.9-7.3*$10^{-7}$ mol $m^{-2}$ $s^{-1}$ with assuming leaf fresh weight to dry weight ratio of 3.5. Furthermore, it is shown in Walker et al. (2014) that leaf photosynthetic nitrogen assimilation rate is 0.17 μmol $m^{-2}$ $s^{-1}$ when leaf photosynthetic rate is 22 μmol $m^{-2}$ $s^{-1}$. Thus we set the $v_{m\_leaf\_N\_ass}$=6.18*$10^{-7}$ mol $m^{-2}$ $s^{-1}$ (0.17/22* $v_{cmax0}$).

[7] It is shown that sucrose-phosphate synthase activity in mature rice leaves is 6.5-10.5 μmol $m^{-2}$ $s^{-1}$, which equals to 3.3-5.3*$10^{-6}$ mol C12 $m^{-2}$ $s^{-1}$.

[8] It is shown that ADP-glucose pyrophosphorylase activity in mature rice leaves is 600-1000 nmol $min^{-1}$ $g^{-1}$ fresh weight, which equals to 8.8-14.6*$10^{-7}$ mol C12 $m^{-2}$ $s^{-1}$ with assuming leaf fresh weight to dry weight ratio of 3.5 and specific leaf area of 0.019-0.037.

[9] X. Yin and H. H. van Laar (2005) set $\alpha_{sene\_leaf}$=0.03 for wheat, which usually has shorter grain filling duration than rice. Therefore, here we set a smaller $\alpha_{sene\_leaf}$ (0.015) for rice.

[10] Rice glume volume is calculated by assuming the white grain as an ellipsoid.

[11] Rice husk accounts for 20% of the total weight of the rough rice grain (Carlile et al., 2019). We further assume total weight of the rough rice grain equals to $V_{glume}$*$10^6$.

[12] It is shown in J. Yang et al. (2006) that the fastest endosperm cell division rate is 50*$10^3$ $d^{-1}$ $grain^{-1}$ for rice cultivar LY9, which is reached at 5 days after flowering when grain volume accounts for 5/12 of the final grain volume at harvest. The final endosperm cell number is 2.4*$10^5$ and the endosperm cell size of rice is about 80*$10^{-15}$ $m^3$ $cell^{-1}$ (J. Yang et al., 2002), based on which the surface area is estimated to be 2.24*$10^{-5}$ $m^2$ at 5 days after flowering. Thus the grain volume expansion rate is 50*$10^3$*80*$10^{-15}$/86400/2.24*$10^{-5}$ = 2.1*$10^{-9}$ $m^3$ $m^{-2}$ $s^{-1}$. Finally we set the maximal potential of grain volume expansion $v_{m\_grain\_grow}$ to be 4*$10^{-9}$ $m^3$ $m^{-2}$ $s^{-1}$.

[13] ADP-glucose pyrophosphorylase (AGP) is the rate-limiting enzyme in starch synthesis. It has been shown the maximal AGP activity in grain is reached at 10 days after flowering (Smidansky et al., 2003). Moreover, it is reported that the AGP activity is 0.3-12 nmol $grain^{-1}$ $min^{-1}$ around 10 days after flowering (S. Chen et al., 2011; Devi et al., 2010; Perez et al., 1975), when the fresh weight of grain is around 8 mg, and water content to be 50% (Devi et al., 2010), then we can estimate the potential starch synthesis rate $v_{m\_grain\_Star\_syn}$=0.3-12*$10^{-9}$/(8*$10^{-9}$*0.5)/60=0.0013-0.05 mol C6 $m^{-3}$ $s^{-1}$=0.63-25*$10^{-3}$ mol C12 $m^{-3}$ $s^{-1}$.

[14] It is known that in rice grain, starch accounts for 70-95% and protein accounts for

5-11% of the total dry weight, respectively (Carlile et al., 2019; Martin & Fitzgerald, 2002; Peng et al., 2014; Y. Yang et al., 2019). Namely, grain starch content (GSC) is 70-95% and grain N content (GNC) is 5-11%/6.25=0.8-1.8%. Thus, $v_{m\_grain\_Pro\_syn}$ is estimated to be (GNC/14)/(GSC/324)* $v_{m\_grain\_Star\_syn}$=0.05-5.7*$10^{-3}$ mol N $m^{-3}$ $s^{-1}$.

[15] It has been reported that rice straw structural dry weight is 21.3-25.7 g $plant^{-1}$ (J. Yang et al., 2001); and it has been reported that rice straw total dry weight is 34.6-68.6 g $plant^{-1}$ (K. Liu et al., 2020), which is equivalent to structural dry weight of 29.4-58.3 g $plant^{-1}$ if assuming 15% is non-structural matter. With a further assumption of 2/3 of the straw structural dry weight belongs to stem and the other 1/3 belongs to leaves, the straw structural dry weight of 21.3-58.3 g $plant^{-1}$ is equivalent to stem structural dry weight of 14.2-38.9 g $plant^{-1}$.

[16] If we assume the dry:fresh weight ratio of stem to be 1:3, and further we assume that the stem aqueous space accounts for 20% volume of the total stem, then $V_{stem\_aq}$=14.2-38.9*$10^{-6}$/(1/3)*20%=0.85-2.33*$10^{-5}$ $m^{-3}$.

[17] It is shown that starch accounts for 5.1-27.3% of the total stem weight at flowering (Arai-Sanoh et al., 2011). Thus, CStar0= SSW0/(72.7-94.9%)*(27.3-5.1%)=0.8-14.6 g.

[18] ADP-glucose pyrophosphorylase (AGP) is the rate-limiting enzyme in starch synthesis. It has been shown that stem AGP activity is 50-400 nmol $min^{-1}$ g fresh $weight^{-1}$ (Hashida et al., 2018; Okamura et al., 2013). If we assume that the stem aqueous space accounts for 20% volume of the total stem, then $v_{m\_stem\_Star\_syn}$=50-400*$10^{-9}$/60*$10^{6}$/20%=0.004-0.033 mol C6 $m^{-3}$ $s^{-1}$=0.21-1.67*$10^{-2}$ mol C12 $m^{-3}$ $s^{-1}$.

[19] It has been reported that phloem area (PA) and xylem area (XA) consisting peduncle vascular bundles at flowering are 0.048-0.225 $mm^2$ and 0.071-0.286 $mm^2$ per culm, respectively. As tiller number (TN) is 2-31 (Qu et al., 2017; Yoshida, 1981), plant height (PH) is 60-120 cm (X. Huang et al., 2015), the total xylem volume XV0=XA*TN*PH=0.085-10.6*$10^{-6}$ $m^3$ $plant^{-1}$; and the total phloem volume PV0=0.058-8.4*$10^{-6}$ $m^3$ $plant^{-1}$.

[20] According to Hayashi and Chino (1990), concentrations of sucrose and amino acids in phloem of one-week-after-flowering rice plant are 124.8±25.6 and 573.8±123.1 mol $m^{-3}$, respectively. Glutamine and Asparagine, which carry 2 N molecules, are two major forms of the amino acids. Therefore, phloem nitrogen-to-sucrose molar ratio at flowering is ~0.4 (124.8*2/573.8), from which we set $v_{m\_root\_ON\_ul}$=0.32*$v_{m\_root\_Suc\_ul}$, $v_{m\_grain\_ON\_ul}$=0.4*$v_{m\_grain\_Suc\_ul}$ and $v_{m\_leaf\_ON\_l}$=0.4*$v_{m\_leaf\_Suc\_l}$.

[21] According to Tilman (1991), maximal root relative growth rate is 0.13-0.28 g $g^{-1}$ $d^{-1}$, which costs 0.13-0.28*$\eta_{grow}$=0.17-0.36 g suc $g^{-1}$ $d^{-1}$=5.8-12.2*$10^{-9}$ mol $g^{-1}$ $s^{-1}$; according to Farrar and Williams (1991), 15-day old barley root sucrose import rate can reach 1mg $h^{-1}$=9.5*$10^{-10}$ mol $s^{-1}$, when the root dry weight is about 0.04 g per plant (Crossett & Campbell, 1975), which is 9.5*$10^{-10}$/0.04=2.4*$10^{-8}$ mol $g^{-1}$ $s^{-1}$.

[22] We assume leaf I-N unloading capacity should match root I-N loading capacity. Namely, $v_{m\_leaf\_IN\_ul}$= $v_{m\_root\_IN\_l}$*RW0/LA0=6*10$^{-8}$ mol m$^{-2}$ s$^{-1}$.

[23] We assume leaf O-N unloading capacity should match root O-N loading capacity. Namely, $v_{m\_leaf\_ON\_ul}$= $v_{m\_root\_ON\_l}$*RW0/LA0=8*10$^{-8}$ mol m$^{-2}$ s$^{-1}$.

[24] As mentioned above, the specific leaf area (SLA) is 0.019-0.037 m$^2$ g$^{-1}$ (G. Li et al., 2009; Xiong et al., 2016). Based on the difference between the non-structural carbohydrates contents measured at end-of-day and end-of-night from Eom et al. (2011), it can be estimated that leaf average night sucrose export rate is 0.08-0.15 μmol Suc m$^{-2}$ s$^{-1}$ [36*3.5/SLA/(12*3600)] by assuming leaf fresh weight to dry weight ratio of 3.5. Using the same strategy, it can be estimated from L. Wang et al. (2015) that leaf average night sucrose export rate is 0.25-0.48 μmol Suc m$^{-2}$ s$^{-1}$. We finally assume $v_{m\_leaf\_Suc\_l}$ to be about five times of the average night sucrose export rate (0.25 μmol Suc m$^{-2}$ s$^{-1}$), i.e., 1.2*10$^{-6}$ mol Suc m$^{-2}$ s$^{-1}$.

[25] According to N. Wang and Fisher (1994), grain sucrose unloading rate of wheat can reach 262 nmol h$^{-1}$ grain$^{-1}$ around 20-day after flowering. If we assume the grain width and length to be 3.3 mm and 7 mm, respectively, we can get a sucrose unloading rate of 4.2*10$^{-6}$ mol m$^{-2}$ s$^{-1}$ on a grain surface area base. We set $v_{m\_grain\_Suc\_ul}$=5.0*10$^{-6}$ mol m$^{-2}$ s$^{-1}$ for rice.

[26] Following the way describing metabolites exchange between cells through the plasmodesmata by Y. Wang et al. (2014), the diffusion rate between leaf-phloem and stem can be written as

$$v = D/L*S*\varphi*([A]_{LP} - [A]_{stem})$$

where diffusion coefficient D is set as 3.4*10$^{-10}$ m s$^{-1}$ and 7.7*10$^{-10}$ m s$^{-1}$ for sucrose and amino acids here, respectively. L=4*10$^{-7}$ m and φ=0.03 are used following Y. Wang et al. (2014). The interface area S is estimated to be 5.3*10$^{-4}$ m$^{-2}$ (B. Zhao, P. Wang, H.-x. Zhang, Q. Zhu, & J. Yang, 2006), so that $R_{Suc\_LP\_stem}$=1/( D/L*S*φ)=7.4*10$^7$ s m$^{-3}$, $R_{AA\_LP\_stem}$=3.2*10$^7$ s m$^{-3}$.

[27] We assume xylem-to-phloem O-N transfer capacity is about 0.6-fold of the root O-N loading capacity. Namely, $v_{m\_ON\_VBtransfer}$= $v_{m\_root\_IN\_l}$*RW0*0.6=1.2*10$^{-8}$ mol m$^{-2}$ s$^{-1}$.

[28] According to Plaxton (1996), it may cost 1 to 3 ATPs with different metabolism pathway to synthesize 1 molecule of sucrose from starch. A relationship of 1 Suc=60 ATPs is used throughout the model, which gives $\eta_{store}$=1-3/60=0.017-0.05.

Abbreviations: E.F., estimated from; DW, dry weight; N, nitrogen; C, carbon.

**Supplementary Table 3. A list of key model parameters for high yield, and reported related genes or quantitative trait loci (QTLs) controlling capacities of the underlying processes in rice.**

| Parameter | Gene/QTL | Functional annotation | Source |
|---|---|---|---|
| $v_{m\_grain\_Star\_syn}$ | *OsPK2* | Encodes a plastidic pyruvate kinase involved in rice | (Cai et al., 2018) |

| | | | |
|---|---|---|---|
| | | endosperm starch synthesis | |
| | *OsAGPL2*, *OsAGPL3*, *OsAGPS2b* | Encode subunits of a starch biosynthesis enzyme ADP-glucose pyrophosphorylase (AGP), and are preferentially expressed in the endosperm | (Lee et al., 2007; Okamura et al., 2013; X. Wei et al., 2017) |
| $R_{\_phloem\_leaf\_root}$ | – | | |
| $v_{m\_root\_N\_upt}$ | *NRT1.1B* | Encodes a nitrate-transporter | (Hu et al., 2015) |
| | *OsNAR2.1*, *OsNRT2.1* | *OsNRT2.1* encodes a high-affinity nitrate transporter, which requires the protein encoded by *OsNAR2.1* for nitrate uptake | (J. Chen et al., 2017; J. Chen et al., 2016) |
| | *OsNRT2.3b* | Encodes a pH-sensitive nitrate transporter | (X. Fan et al., 2016) |
| $v_{m\_leaf\_Suc\_l}$ | *OsSUT2* | Encodes a tonoplast-localized sucrose transporter for sucrose uptake from the vacuole, and plays an essential role in sugar export from leaves | (Eom et al., 2011) |
| $v_{m\_leaf\_Suc\_syn}$ | *OsSPS1*, *OsSPS11* | Encode sucrose phosphate synthase, a key enzyme in sucrose biosynthesis | (Hashida et al., 2016) |
| $v_{m\_leaf\_Pro\_deg}$ | *OsNAP* | Encodes a plant-specific NAC transcriptional activator, which positively regulates leaf senescence by targeting genes related to chlorophyll degradation and other genes associated with senescence | (Liang et al., 2014) |
| | *OsNAC2* | Encodes a rice NAC transcription factor that participates in ABA-induced leaf senescence by directly activating expression of chlorophyll degradation genes | (Mao et al., 2017) |
| | *OsMTS1* | Encodes an O-methyltransferase in the melatonin biosynthetic pathway, which is associated with trigger of leaf senescence | (Y. Hong et al., 2018) |
| $v_{m\_leaf\_ON\_l}$ | – | | |
| $v_{m\_grain\_grow}$ | *OsNF-YB1* | Is an endosperm-specific expressed gene, encodes a cell cycle regulator and plays a role in maintaining endosperm cell proliferation | (X. Sun et al., 2014) |
| | *OsKRP1* | Encodes a KIP-related protein, a cyclin-dependent kinase (CDK) inhibitor, and controls cell cycle | (Barrôco et al., 2006) |
| | *OsCycB1;1*, | Encodes a B type cyclin, which is critical for endosperm formation | (Guo, Wang, Song, Sun, & Zhang, 2010) |
| | *OsCCS52A* | Is an anaphase-promoting complex activator that is involved in maintaining normal seed size formation by mediating the exit from mitotic cell division to enter the endoreduplication cycles in rice endosperm | (Su'Udi et al., 2012) |
| | *OsSPL16* | Encodes a positive regulator of cell proliferation | (S. Wang et al., 2012) |
| $v_{m\_root\_Suc\_ul}$ | – | | |
| $v_{m\_stem\_Star\_syn}$ | *OsAGPL1* | Encodes a large subunit of ADP-glucose pyrophosphorylase (AGP) and are preferentially expressed in the stem | (Okamura et al., 2013) |

#### References


Allen, R. G., Pereira, L. S., Raes, D., & Smith, M. (1998). Crop evapotranspiration-Guidelines for computing crop water requirements-FAO Irrigation and drainage paper 56. *FAO, Rome, 300*(9), D05109.

Andrews, M. (1986). The partitioning of nitrate assimilation between root and shoot of higher plants. *Plant, Cell and Environment, 9*(7), 511-519.

Arai-Sanoh, Y., Ida, M., Zhao, R., Yoshinaga, S., Takai, T., Ishimaru, T., . . . Kondo, M. (2011). Genotypic variations in non-structural carbohydrate and cell-wall components of the stem in rice, sorghum, and sugar cane. *Bioscience, Biotechnology, and Biochemistry, 75*(6), 1104-1112. doi:10.1271/bbb.110009

Asseng, S., Richter, C., & Wessolek, G. (1997). Modelling root growth of wheat as the linkage between crop and soil. *Plant and Soil, 190*(2), 267-277. doi:10.1023/A:1004228201299

Azcón-Bieto, J. (1983). Inhibition of photosynthesis by carbohydrates in wheat leaves. *Plant Physiology, 73*(3), 681-686.

Baloch, A. W., Soomro, A. M., Javed, M. A., Ahmed, M., & Mastoi, N. N. (2002). Optimum plant density for high yield in rice ( *Oryza sativa* L.). *Asian Journal of Plant Sciences, 1*(1).

Barrôco, R. M., Peres, A., Droual, A.-M., De Veylder, L., Nguyen, L. S. L., De Wolf, J., . . . Frankard, V. (2006). The cyclin-dependent kinase inhibitor Orysa;KRP1 plays an important role in seed development of rice. *Plant Physiology, 142*(3), 1053-1064. doi:10.1104/pp.106.087056

Cai, Y., Li, S., Jiao, G., Sheng, Z., Wu, Y., Shao, G., . . . Hu, P. (2018). *OsPK2* encodes a plastidic pyruvate kinase involved in rice endosperm starch synthesis, compound granule formation and grain filling. *Plant Biotechnology Journal, 16*(11), 1878-1891. doi:10.1111/pbi.12923

Cannell, M. G. R., & Thornley, J. H. M. (2000). Modelling the components of plant respiration: Some guiding principles. *Annals of Botany, 85*(1), 45-54. doi:DOI 10.1006/anbo.1999.0996

Carlile, W. R., Raviv, M., & Prasad, M. (2019). Chapter 8 - Organic soilless media components. In M. Raviv, J. H. Lieth, & A. Bar-Tal (Eds.), *Soilless Culture (Second Edition)* (pp. 303-378). Boston: Elsevier.

Chang, T. G., Xin, C. P., Qu, M. N., Zhao, H. L., Song, Q. F., & Zhu, X. G. (2017). Evaluation of protocols for measuring leaf photosynthetic properties of field-grown rice. *Rice Science, 24*(1), 1-9.

Chen, G. P., Cheng, L., Zhu, J. G., Pang, J., Xie, Z. B., & Zeng, Q. (2006). Effects of free-air $CO_2$ enrichment on root characteristics and C:N ratio of rice at the heading stage. *Rice Science, 13*(2), 120-124.

Chen, J., Fan, X., Qian, K., Zhang, Y., Song, M., Liu, Y., . . . Fan, X. (2017). *pOsNAR2.1:OsNAR2.1* expression enhances nitrogen uptake efficiency and grain yield in transgenic rice plants. *Plant Biotechnology Journal, 15*(10), 1273-1283. doi:10.1111/pbi.12714

Chen, J., Zhang, Y., Tan, Y., Zhang, M., Zhu, L., Xu, G., & Fan, X. (2016). Agronomic nitrogen-use efficiency of rice can be increased by driving *OsNRT2.1* expression with the *OsNAR2.1* promoter. *Plant Biotechnology Journal, 14*(8), 1705-1715. doi:10.1111/pbi.12531

Chen, S., Zhou, W., Zeng, F., & Zhang, G. (2011). Effect of planting method on grain quality and nutrient utilization for no-tillage rice. *Communications in Soil Science and Plant Analysis, 42*(11), 1324-1335. doi:10.1080/00103624.2011.571737

Crossett, R. N., & Campbell, D. J. (1975). The effects of ethylene in the root environment upon the development of barley. *Plant and Soil, 42*(2), 453-464.

De Pury, D., & Farquhar, G. (1997). Simple scaling of photosynthesis from leaves to canopies without the errors of big-leaf models. *Plant, Cell and Environment, 20*(5), 537-557.

De Visser, R., Spitters, C., & Bouma, T. (1992). Energy cost of protein turnover: theorectical calculation and experimental estimation from regression of respiration on protein concentration of fulle-grown leaves *Molecular, biochemical and physiological aspects of plant respiration* (pp. 493-508): Academic Publishing.

Devi, T. A., Sarla, N., Siddiq, E. A., & Sirdeshmukh, R. (2010). Activity and expression of adenosine diphosphate glucose pyrophosphorylase in developing rice grains: Varietal differences and implications on grain filling. *Plant Science, 178*(2), 123-129. doi:https://doi.org/10.1016/j.plantsci.2009.10.008

Eom, J.-S., Cho, J.-I., Reinders, A., Lee, S.-W., Yoo, Y., Tuan, P. Q., . . . Cho, M.-H. (2011). Impaired function of the tonoplast-localized sucrose transporter in rice, OsSUT2, limits the transport of vacuolar reserve sucrose and affects plant growth. *Plant Physiology, 157*(1), 109-119.

Fan, C., Xing, Y., Mao, H., Lu, T., Han, B., Xu, C., . . . Zhang, Q. (2006). GS3, a major QTL for grain length and weight and minor QTL for grain width and thickness in rice, encodes a putative transmembrane protein. *Theoretical and Applied Genetics, 112*(6), 1164-1171.

Fan, X., Tang, Z., Tan, Y., Zhang, Y., Luo, B., Yang, M., . . . Xu, G. (2016). Overexpression of a pH-sensitive nitrate transporter in rice increases crop yields. *Proceedings of the National academy of Sciences of the United States of America, 113*(26), 7118-7123. doi:10.1073/pnas.1525184113

Fanello, D. D., Bartoli, C. G., & Guiamet, J. J. (2017). Qualitative and quantitative modifications of root mitochondria during senescence of above-ground parts of *Arabidopis thaliana*. *Plant Science, 258*, 112-121. doi:10.1016/j.plantsci.2017.01.013

Farquhar, G. D., & Sharkey, T. D. (1982). Stomatal conductance and photosynthesis. *Annual Review of Plant Physiology, 33*(1), 317-345.

Farrar, J. F., & Williams, J. H. H. (1991). *Control of the rate of respiration in roots: compartmentation, demand and the supply of substrate.* Paper presented at the Seminar series - Society for Experimental Biology.

Feugier, F. G., & Satake, A. (2013). Dynamical feedback between circadian clock and sucrose availability explains adaptive response of starch metabolism to various photoperiods. *Frontiers in Plant Science, 3*, 305.

Fitzgerald, M. A., McCouch, S. R., & Hall, R. D. (2009). Not just a grain of rice: the quest for quality. *Trends in Plant Science, 14*(3), 133-139. doi:https://doi.org/10.1016/j.tplants.2008.12.004

Fukushima, A., Shiratsuchi, H., Yamaguchi, H., & Fukuda, A. (2011). Effects of nitrogen application and planting density on morphological traits, dry matter production and yield of large grain type rice variety Bekoaoba and strategies for super high-yielding rice in the Tohoku region of Japan. *Plant Production Science, 14*(1), 56-63.

Gesch, R. W., Vu, J. C. V., Boote, K. J., Allen, L. H., & Bowes, G. (2002). Sucrose-phosphate synthase activity in mature rice leaves following changes in growth $CO_2$ is unrelated to sucrose pool size. *New Phytologist, 154*(1), 77-84. doi:DOI 10.1046/j.1469-8137.2002.00348.x

Glass, A. D. M., Britto, D. T., Kaiser, B. N., Kinghorn, J. R., Kronzucker, H. J., Kumar, A., . . . Vidmar, J. J. (2002). The regulation of nitrate and ammonium transport systems in plants. *Journal of Experimental Botany, 53*(370), 855-864. doi:DOI 10.1093/jexbot/53.370.855

Gu, J., Yin, X., Stomph, T.-J., & Struik, P. C. (2014). Can exploiting natural genetic variation in leaf photosynthesis contribute to increasing rice productivity? A simulation analysis. *Plant, Cell and Environment, 37*(1), 22-34. doi:10.1111/pce.12173

Guo, J., Wang, F., Song, J., Sun, W., & Zhang, X. (2010). The expression of *Orysa;CycB1;1* is essential for endosperm formation and causes embryo enlargement in rice. *Planta, 231*(2), 293-303. doi:10.1007/s00425-009-1051-y

Haque, M. M., Pramanik, H. R., Biswas, J. K., Iftekharuddaula, K., & Hasanuzzaman, M. (2015). Comparative performance of hybrid and elite inbred rice varieties with respect to their source-sink relationship. *The Scientific World Journal, 2015*.

Hashida, Y., Hirose, T., Okamura, M., Hibara, K.-i., Ohsugi, R., & Aoki, N. (2016). A reduction of sucrose phosphate synthase (SPS) activity affects sucrose/starch ratio in leaves but does not inhibit normal plant growth in rice. *Plant Science, 253*, 40-49.

Hashida, Y., Kadoya, S., Okamura, M., Sugimura, Y., Hirano, T., Hirose, T., . . . Aoki, N. (2018). Characterization of sugar metabolism in the stem of Tachisuzuka, a whole-crop silage rice cultivar with high sugar content in the stem. *Plant Production Science, 21*(3), 233-243. doi:10.1080/1343943X.2018.1461016

Hayashi, H., & Chino, M. (1990). Chemical-composition of phloem sap from the uppermost internode of the rice plant. *Plant and Cell Physiology, 31*(2), 247-251.

Henry, L. T., & Raper Jr, C. D. (1991). Soluble carbohydrate allocation to roots, photosynthetic rate of leaves, and nitrate assimilation as affected by nitrogen stress and irradiance. *Botanical Gazette*, 23-33.

Hong, D., & Ichii, M. (1996). Studies on agronomic characters of short root mutants in rice (*Oryza sativa* L. ). *Chinese Journal of Rice Science, 10*(1), 57-61.

Hong, Y., Zhang, Y., Sinumporn, S., Yu, N., Zhan, X., Shen, X., . . . Cao, L. (2018). Premature leaf senescence 3, encoding a methyltransferase, is required for melatonin biosynthesis in rice. *The Plant Journal, 95*(5), 877-891. doi:10.1111/tpj.13995

Hu, B., Wang, W., Ou, S., Tang, J., Li, H., Che, R., . . . Chu, C. (2015). Variation in *NRT1.1B* contributes to nitrate-use divergence between rice subspecies. *Nature Genetics, 47*(7), 834-838. doi:10.1038/ng.3337

Huang, H. (1998). Relation between the tissue of the highest internode and the number of spikelets. *Acta Agronomica Sinica, 24*(2), 193-200.

Huang, X., Wei, X., Sang, T., Zhao, Q., Feng, Q., Zhao, Y., . . . Zhang, Z. (2010). Genome-wide association studies of 14 agronomic traits in rice landraces. *Nature Genetics, 42*(11), 961-967.

Huang, X., Yang, S., Gong, J., Zhao, Y., Feng, Q., Gong, H., . . . Xia, J. (2015). Genomic analysis of hybrid rice varieties reveals numerous superior alleles that contribute to heterosis. *Nature Communications, 6*, 6258.

Jarvis, P. (1976). The interpretation of the variations in leaf water potential and stomatal conductance found in canopies in the field. *Philosophical Transactions of the Royal Society of London B: Biological Sciences, 273*(927), 593-610.

Johnson, I. R., & Thornley, J. H. M. (1985). Dynamic model of the response of a vegetative grass crop to light, temperature and nitrogen. *Plant, Cell and Environment, 8*(7), 485-499. doi:10.1111/j.1365-3040.1985.tb01684.x

Kim, H., Lieffering, M., Miura, S., Kobayashi, K., & Okada, M. (2001). Growth and nitrogen uptake of $CO_2$ ‐ enriched rice under field conditions. *New Phytologist, 150*(2), 223-229.

Kiniry, J. R., McCauley, G., Xie, Y., & Arnold, J. G. (2001). Rice parameters describing crop performance of four US cultivars. *Agronomy Journal, 93*(6), 1354-1361.

Knoblauch, M., Knoblauch, J., Mullendore, D. L., Savage, J. A., Babst, B. A., Beecher, S. D., . . . Holbrook, N. M. (2016). Testing the Munch hypothesis of long distance phloem transport in plants. *eLife, 5*. doi:ARTN e15341

10.7554/eLife.15341

Kobata, T., Sugawara, M., & Takatu, S. (2000). Shading during the early grain filling period does not affect potential grain dry matter increase in rice. *Agronomy Journal, 92*(3), 411-417.

Lalonde, S., Tegeder, M., Throne ‐ Holst, M., Frommer, W., & Patrick, J. (2003). Phloem loading and unloading of sugars and amino acids. *Plant, Cell and Environment, 26*(1), 37-56.

Lee, S.-K., Hwang, S.-K., Han, M., Eom, J.-S., Kang, H.-G., Han, Y., . . . Jeon, J.-S. (2007). Identification of the ADP-glucose pyrophosphorylase isoforms essential for starch synthesis in the leaf and seed endosperm of rice (*Oryza sativa* L.). *Plant Molecular Biology 65*(4), 531-546. doi:10.1007/s11103-007-9153-z

Leuning, R. (1995). A critical appraisal of a combined stomatal-photosynthesis model for $C_3$ plants. *Plant, Cell and Environment, 18*(4), 339-355.

Li, G., Xue, L., Gu, W., Yang, C., Wang, S., Ling, Q., . . . Ding, Y. (2009). Comparison of yield components and plant type characteristics of high-yield rice between Taoyuan, a 'special eco-site' and Nanjing, China. *Field Crops Research, 112*(2), 214-221.

Li, Y. G., Ouyang, J., Wang, Y. Y., Hu, R., Xia, K. F., Duan, J., . . . Zhang, M. Y. (2015). Disruption of the rice nitrate transporter OsNPF2.2 hinders root-to-shoot nitrate transport and vascular development. *Scientific Reports, 5*. doi:Artn 9635

10.1038/Srep09635

Liang, C., Wang, Y., Zhu, Y., Tang, J., Hu, B., Liu, L., . . . Chu, J. (2014). OsNAP connects abscisic acid and leaf senescence by fine-tuning abscisic acid biosynthesis and directly targeting senescence-associated genes in rice. *Proceedings of the National academy of Sciences of the United States of America, 111*(27), 10013-10018.

Liu, H. (2009). Effect of water-saving irrigation on dry matter and nitrogen content of rice root. *Shandong Agricultural Sciences*.

Liu, K., Yang, R., Deng, J., Huang, L., Wei, Z., Ma, G., . . . Zhang, Y. (2020). High radiation use efficiency improves yield in the recently developed elite hybrid rice Y-liangyou 900. *Field Crops Research, 253*, 107804. doi:https://doi.org/10.1016/j.fcr.2020.107804

Liu, L. J., Chen, T. T., Wang, Z. Q., Zhang, H., Yang, J. C., & Zhang, J. H. (2013). Combination of site-specific nitrogen management and alternate wetting and drying irrigation increases grain yield and nitrogen and water use efficiency in super rice. *Field Crops Research, 154*, 226-235. doi:10.1016/j.fcr.2013.08.016

Malagoli, P., Lainé, P., Le Deunff, E., Rossato, L., Ney, B., & Ourry, A. (2004). Modeling nitrogen uptake in oilseed rape cv capitol during a growth cycle using influx kinetics of root nitrate transport systems and field experimental data. *Plant Physiology, 134*(1), 388. doi:10.1104/pp.103.029538

Mao, C., Lu, S., Lv, B., Zhang, B., Shen, J., He, J., . . . Ming, F. (2017). A Rice NAC Transcription Factor Promotes Leaf Senescence via ABA Biosynthesis. *Plant Physiology, 174*(3), 1747-1763. doi:10.1104/pp.17.00542

Martin, M., & Fitzgerald, M. A. (2002). Proteins in rice grains influence cooking properties. *Journal of Cereal Science, 36*(3), 285-294. doi:https://doi.org/10.1006/jcrs.2001.0465

Marwaha, R. S., & Juliano, B. O. (1976). Aspects of nitrogen metabolism in the rice seedling. *Plant Physiology, 57*(6), 923-927.

Masclaux‐Daubresse, C., Reisdorf‐Cren, M., & Orsel, M. (2008). Leaf nitrogen remobilisation for plant development and grain filling. *Plant Biology, 10*(s1), 23-36.

Miflin, B., & Lea, P. (1980). Ammonia assimilation. *The Biochemistry of Plants, 5*, 169-202.

Minchin, P., Thorpe, M., & Farrar, J. (1993). A simple mechanistic model of phloem transport which explains sink priority. *Journal of Experimental Botany, 44*(5), 947-955.

Monod, J. (1949). The growth of bacterial cultures. *Annual Reviews in Microbiology, 3*(1), 371-394.

Mottaleb, K. A., & Mishra, A. K. (2016). Rice consumption and grain-type preference by household: a Bangladesh case. *Journal of Agricultural and Applied Economics, 48*(3), 298-319. doi:10.1017/aae.2016.18

Muller, B., Pantin, F., Génard, M., Turc, O., Freixes, S., Piques, M., & Gibon, Y. (2011). Water deficits uncouple growth from photosynthesis, increase C content, and modify the relationships between C and growth in sink organs. *Journal of Experimental Botany*, erq438.

Nunes-Nesi, A., Fernie, A. R., & Stitt, M. (2010). Metabolic and signaling aspects underpinning the regulation of plant carbon nitrogen interactions. *Molecular Plant, 3*(6), 973-996. doi:10.1093/mp/ssq049

Ober, E. S., Setter, T. L., Madison, J. T., Thompson, J. F., & Shapiro, P. S. (1991). Influence of water deficit on maize endosperm development : enzyme activities and RNA transcripts of starch and zein synthesis, abscisic acid, and cell division. *Plant Physiology, 97*(1), 154-164. doi:10.1104/pp.97.1.154

Ohdan, T., Francisco, P. B., Jr, Sawada, T., Hirose, T., Terao, T., Satoh, H., & Nakamura, Y. (2005). Expression profiling of genes involved in starch synthesis in sink and source organs of rice. *Journal of Experimental Botany, 56*(422), 3229-3244. doi:10.1093/jxb/eri292

Okamura, M., Hirose, T., Hashida, Y., Yamagishi, T., Ohsugi, R., & Aoki, N. (2013). Starch reduction in rice stems due to a lack of OsAGPL1 or OsAPL3 decreases grain yield under low irradiance during ripening and modifies plant architecture. *Functional Plant Biology, 40*(11), 1137-1146. doi:10.1071/FP13105

Olsen, O. A. (2004). Nuclear endosperm development in cereals and *Arabidopsis thaliana*. *The Plant Cell, 16 Suppl*, S214-227. doi:10.1105/tpc.017111

Ou-Lee, T. M., & Setter, T. L. (1985). Enzyme activities of starch and sucrose pathways and growth of apical and basal maize kernels. *Plant Physiology, 79*(3), 848-851. doi:10.1104/pp.79.3.848

Paul, M. J., & Pellny, T. K. (2003). Carbon metabolite feedback regulation of leaf photosynthesis and development. *J Exp Bot, 54*(382), 539-547. doi:10.1093/jxb/erg052

Peng, B., Kong, H. L., Li, Y. B., Wang, L. Q., Zhong, M., Sun, L., . . . He, Y. Q. (2014). OsAAP6 functions as an important regulator of grain protein content and nutritional quality in rice. *Nature Communications, 5*. doi:Artn 4847

10.1038/Ncomms5847

Perez, C. M., Perdon, A. A., Resurreccion, A. P., Villareal, R. M., & Juliano, B. O. (1975). Enzymes of carbohydrate metabolism in the developing rice grain. *Plant Physiology, 56*(5), 579-583.

Pilkington, S. M., Encke, B., Krohn, N., Hoehne, M., Stitt, M., & PYL, E. T. (2015). Relationship between starch degradation and carbon demand for maintenance and growth in *Arabidopsis thaliana* in different irradiance and temperature regimes. *Plant, Cell and Environment, 38*(1), 157-171.

Pitman, M., & Cram, W. (2013). Regulation of inorganic ion transport in plants. *Ion Transport in Plants*, 465-481.

Plaxton, W. C. (1996). The organization and regulation of plant glycolysis. *Annual Review of Plant Physiology and Plant Molecular Biology, 47*, 185-214. doi:DOI 10.1146/annurev.arplant.47.1.185

Pury, D. d., & Farquhar, G. (1997). Simple scaling of photosynthesis from leaves to canopies without the errors of big‐leaf models. *Plant, Cell & Environment, 20*(5), 537-557.

Qu, M., Zheng, G., Essmine, J., Hamdani, S., Song, Q., Wang, H., . . . Zhu, X.-G. (2017). Leaf photosynthetic parameters related to biomass accumulation in a global rice diversity survey. *Plant Physiology*, pp. 00332.02017.

Radin, J. W., Parker, L. L., & Sell, C. R. (1978). Partitioning of sugar between growth and nitrate reduction in cotton roots. *Plant Physiology, 62*(4), 550-553.

San-oh, Y., Sugiyama, T., Yoshita, D., Ookawa, T., & Hirasawa, T. (2006). The effect of planting pattern on the rate of photosynthesis and related processes during ripening in rice plants. *Field Crops Research, 96*(1), 113-124.

Scialdone, A., Mugford, S. T., Feike, D., Skeffington, A., Borrill, P., Graf, A., . . . Howard, M. (2013). Arabidopsis plants perform arithmetic division to prevent starvation at night. *Elife, 2,* e00669.

Seaton, D. D., Ebenhöh, O., Millar, A. J., & Pokhilko, A. (2014). Regulatory principles and experimental approaches to the circadian control of starch turnover. *Journal of the Royal Society Interface, 11*(91), 20130979.

Shi, Z., Chang, T.-G., Chen, G., Song, Q., Wang, Y., Zhou, Z., . . . Zhu, X.-G. (2019). Dissection of mechanisms for high yield in two elite rice cultivars. *Field Crops Research, 241*, 107563. doi:10.1016/j.fcr.2019.107563

Sinclair, T., & De Wit, C. (1976). Analysis of the carbon and nitrogen limitations to soybean yield. *Agronomy Journal, 68*(2), 319-324.

Smidansky, E. D., Martin, J. M., Hannah, L. C., Fischer, A. M., & Giroux, M. J. (2003). Seed yield and plant biomass increases in rice are conferred by deregulation of endosperm ADP-glucose pyrophosphorylase. *Planta, 216*(4), 656-664. doi:10.1007/s00425-002-0897-z

Stitt, M. (1991). Rising $CO_2$ levels and their potential significance for carbon flow in photosynthetic cells. *Plant, Cell and Environment, 14*(8), 741-762.

Su'Udi, M., Cha, J. Y., Min, H. J., Ermawati, N., Han, C. D., Min, G. K., . . . Son, D. (2012). Potential role of the rice *OsCCS52A* gene in endoreduplication. *Planta, 235*(2), 387-397.

Sun, L., Lu, Y., Yu, F., Kronzucker, H. J., & Shi, W. (2016). Biological nitrification inhibition by rice root exudates and its relationship with nitrogen-use efficiency. *New Phytologist, 212*(3), 646-656. doi:10.1111/nph.14057

Sun, X., Ling, S., Lu, Z., Ouyang, Y.-d., Liu, S., & Yao, J. (2014). *OsNF-YB1*, a rice endosperm-specific gene, is essential for cell proliferation in endosperm development. *Gene, 551*(2), 214-221. doi:https://doi.org/10.1016/j.gene.2014.08.059

Tang, W.-B., Deng, H.-B., Xiao, Y.-H., Zhang, G.-L., Fan, K., Mo, H., & Chen, L.-Y. (2010). Root characteristics of high-yield C Liangyou rice combinations of two-line hybrid rice. *Scientia Agricultura Sinica, 43*(14), 2859-2868. doi:10.3864/j.issn.0578-1752.2010.14.004

Tashiro, T., & Wardlaw, I. F. (1989). A comparison of the effect of high temperature on grain development in wheat and rice. *Annals of Botany, 64*(1), 59-65.

Tazoe, Y., Noguchi, K., & Terashima, I. (2006). Effects of growth light and nitrogen nutrition on the organization of the photosynthetic apparatus in leaves of a $C_4$ plant, Amaranthus cruentus. *Plant, Cell and Environment, 29*(4), 691-700.

Teo, Y. H., Beyrouty, C. A., & Gbur, E. E. (1992). Nitrogen, phosphorus, and potassium influx kinetic parameters of three rice cultivars. *Journal of Plant Nutrition, 15*(4), 435-444. doi:10.1080/01904169209364331

Thornley, J., & Cannell, M. (2000). Modelling the components of plant respiration: representation and realism. *Annals of Botany, 85*(1), 55-67.

Tilman, D. (1991). Relative growth rates and plant allocation patterns. *The American Naturalist, 138*(5), 1269-1275.

Toyosawa, Y., Kawagoe, Y., Matsushima, R., Crofts, N., Ogawa, M., Fukuda, M., . . . Fujita, N. (2016). Deficiency of starch synthase IIIa and IVb alters starch granule morphology from polyhedral to spherical in rice endosperm. *Plant Physiology, 170*(3), 1255-1270. doi:10.1104/pp.15.01232

Von Caemmerer, S. (2000a). *Biochemical models of leaf photosynthesis*: CSIRO publishing.

Von Caemmerer, S. (2000b). *Biochemical models of leaf photosynthesis*: Csiro publishing.

Walker, B. J., Strand, D. D., Kramer, D. M., & Cousins, A. B. (2014). The response of cyclic electron flow around photosystem I to changes in photorespiration and nitrate assimilation. *Plant Physiology, 165*(1), 453-462.

Wang, E., Wang, J., Zhu, X., Hao, W., Wang, L., Li, Q., . . . Lin, H. (2008). Control of rice grain-filling and yield by a gene with a potential signature of domestication. *Nature Genetics, 40*(11), 1370-1374.

Wang, L., Lu, Q., Wen, X., & Lu, C. (2015). Enhanced sucrose loading improves rice yield by increasing grain size. *Plant Physiology*, *169*(4), 2848-2862. doi:10.1104/pp.15.01170

Wang, M. Y., Siddiqi, M. Y., Ruth, T. J., & Glass, A. D. (1993). Ammonium uptake by rice roots (II. Kinetics of $^{13}NH_4^+$ influx across the plasmalemma). *Plant Physiology*, *103*(4), 1259-1267.

Wang, N., & Fisher, D. B. (1994). Monitoring phloem unloading and post-phloem transport by microperfusion of attached wheat grains. *Plant Physiology*, *104*(1), 7-16. doi:Doi 10.1104/Pp.104.1.7

Wang, S., Wu, K., Yuan, Q., Liu, X., Liu, Z., Lin, X., . . . Fu, X. (2012). Control of grain size, shape and quality by *OsSPL16* in rice. *Nature Genetics, 44*(8), 950-954. doi:10.1038/ng.2327

Wang, Y., Long, S. P., & Zhu, X.-G. (2014). Elements required for an efficient NADP-malic enzyme type $C_4$ photosynthesis. *Plant Physiology*, *164*(4), 2231-2246.

Wei, D., Ning, S., & Lin, W. (2004). Relationship between wheat root activity and leaf senescence. *The Journal of Applied Ecology, 15*(9), 1565.

Wei, X., Jiao, G., Lin, H., Sheng, Z., Shao, G., Xie, L., . . . Hu, P. (2017). *GRAIN INCOMPLETE FILLING 2* regulates grain filling and starch synthesis during rice caryopsis development. *Journal of Integrative Plant Biology, 59*(2), 134-153. doi:10.1111/jipb.12510

Willaume, M., & Pagès, L. (2011). Correlated responses of root growth and sugar concentrations to various defoliation treatments and rhythmic shoot growth in oak tree seedlings (*Quercus pubescens*). *Annals of Botany*, mcq270.

Winter, H., Robinson, D. G., & Heldt, H. W. (1993). Subcellular volumes and metabolite concentrations in barley leaves. *Planta*, *191*(2), 180-190.

Xiong, D., Wang, D., Liu, X., Peng, S., Huang, J., & Li, Y. (2016). Leaf density explains variation in leaf mass per area in rice between cultivars and nitrogen treatments. *Annals of Botany*, *117*(6), 963-971. doi:10.1093/aob/mcw022

Yamakawa, Y., Saigusa, M., Okada, M., & Kobayashi, K. (2004). Nutrient uptake by rice and soil solution composition under atmospheric $CO_2$ enrichment. *Plant and Soil, 259*(1), 367-372. doi:10.1023/B:PLSO.0000020988.18365.b5

Yang, J., & Zhang, J. (2010). Grain-filling problem in 'super' rice. *Journal of Experimental Botany, 61*(1), 1-5.

Yang, J., Zhang, J., Huang, Z., Wang, Z., Zhu, Q., & Liu, L. (2002). Correlation of cytokinin levels in the endosperms and roots with cell number and cell division activity during endosperm development in rice. *Annals of Botany, 90*(3), 369-377. doi:10.1093/aob/mcf198

Yang, J., Zhang, J., Wang, Z., Liu, K., & Wang, P. (2006). Post-anthesis development of inferior and superior spikelets in rice in relation to abscisic acid and ethylene. *Journal of Experimental Botany, 57*(1), 149-160.

Yang, J., Zhang, J., Wang, Z., Zhu, Q., & Wang, W. (2001). Remobilization of carbon reserves in response to water deficit during grain filling of rice. *Field Crops Research, 71*(1), 47-55.

Yang, Y., Guo, M., Sun, S., Zou, Y., Yin, S., Liu, Y., . . . Yan, C. (2019). Natural variation of *OsGluA2* is involved in grain protein content regulation in rice. *Nature Communications, 10*(1), 1949. doi:10.1038/s41467-019-09919-y

Yang, Z., Van Oosterom, E. J., Jordan, D. R., & Hammer, G. L. (2009). Pre-anthesis ovary development determines genotypic differences in potential kernel weight in sorghum. *Journal of Experimental Botany*, erp019.

Yeoh, H.-H., & Wee, Y.-C. (1994). Leaf protein contents and nitrogen-to-protein conversion factors for 90 plant species. *Food Chemistry, 49*(3), 245-250. doi:https://doi.org/10.1016/0308-8146(94)90167-8

Yin, X., & van Laar, H. (2005). Crop Systems Dynamics: An Ecophysiological Model of Genotype-by-Environment Interactions (GECROS). *Wageningen Academic Pub., Wageningen*.

Yin, X., & van Laar, H. H. (2005). Crop systems dynamics: an ecophysiological model of genotype-by-environment interactions (GECROS);. Wageningen. *Summary*.

Yoshida, S. (1981). *Fundamentals of rice crop science*: Los Banos, Philippines: International Rice Research Institute.

Youngdahl, L. J., Pacheco, R., Street, J. J., & Vlek, P. L. G. (1982). The kinetics of ammonium and nitrate uptake by young rice plants. *Plant and Soil, 69*(2), 225-232. doi:Doi 10.1007/Bf02374517

Zhang, C. S., Wang, Y. L., Long, Y. C., Dong, G. C., Yang, L. X., & Huang, J. Y. (2005). Main root traits affecting yield level in conventional indica rice cultivars (*Oryza sativa* L.). *Acta Agronomica Sinica, 31*.

Zhang, J., & Davies, W. (1990). Changes in the concentration of ABA in xylem sap as a function of changing soil water status can account for changes in leaf conductance and growth. *Plant, Cell and Environment, 13*(3), 277-285.

Zhao, B., Wang, P., Zhang, H.-x., Zhu, Q., & Yang, J. (2006). Source-sink and grain filling characteristics of two-line hybrid rice Yangliangyou 6. *Rice Science, 13*(1), 34-42.

Zhao, B. H., Wang, P., Zhang, H., Zhu, Q. S., & Yang, J. C. (2006). Source-sink and grain-filling characteristics of two-line hybrid rice Yangliangyou 6. *Rice Science, 13*(1), 34-42.

Zhao, Y., Xi, M., Zhang, X., Lin, Z., Ding, C., Tang, S., . . . Ding, Y. (2015). Nitrogen effect on amino acid composition in leaf and grain of japonica rice during grain filling stage. *Journal of Cereal Science, 64*, 29-33.

Zheng, H., Chen, Y., Chen, Q., Li, B., Zhang, Y., Jia, W., . . . Tang, Q. (2020). High-density planting with lower nitrogen application increased early rice production in a double-season rice system. *Agronomy Journal, 112*(1), 205-214. doi:10.1002/agj2.20033

Zhu, X. G., de Sturler, E., & Long, S. P. (2007). Optimizing the distribution of resources between enzymes of carbon metabolism can dramatically increase photosynthetic rate: A numerical simulation using an evolutionary algorithm. *Plant Physiology, 145*(2), 513-526. doi:DOI 10.1104/pp.107.103713
